# The generation of a comprehensive spectral library for the analysis of the guinea pig proteome by SWATH-MS

**DOI:** 10.1101/514760

**Authors:** Pawel Palmowski, Rachael Watson, G. Nicholas Europe-Finner, Magdalena Karolczak-Bayatti, Andrew Porter, Achim Treumann, Michael J Taggart

## Abstract

Advances in liquid chromatography-mass spectrometry have facilitated the incorporation of proteomic studies to many biology experimental workflows. In particular, the introduction of Data-Independent Acquisition platforms, such as SWATH, offers several advantages for label-free quantitative assessment of complex proteomes over Data-Dependent Acquisition (DDA) approaches. However, SWATH data interpretation requires spectral libraries as a reference resource. This is often not available for many species of experimental models. The guinea pig *(cavia porcellus)* is an excellent experimental model for translation to many aspects of human physiology and disease yet there is limited experimental information regarding its proteome. In an effort to overcome this knowledge gap, we generated a comprehensive spectral library of the guinea pig proteome. Homogenates and tryptic peptide digests were prepared from 16 tissues and subjected to >200 DDA runs. Analysis of >250,000 peptide-spectrum matches resulted in the construction of a library of 73594 peptides corresponding to 7667 proteins. This spectral library furnishes the research community with the first comprehensive guinea pig proteomic resource that will facilitate future molecular-phenotypic studies using (re-engaging) the guinea pig as an experimental model of relevance to human biology. The guinea pig spectral library and MS data are freely accessible in the MassIVE repository (MSV000083199).

## Introduction

Many advances in the speed, accuracy and throughput of liquid chromatography mass spectrometry (LCMS) systems in the last decade have brought proteomic workflows – where the assessment of abundance of several hundreds of proteins can reliably be made from complex biological samples –to the mainstream of biological experimentation. Data-Dependent Acquisition MS modes (DDA), involving the most abundant eluted parent ions of an MS1 scan being selected for fragmentation in MS2 to enable peptide identification and quantification, have contributed considerably in this regard. However, the stochastic nature of parent ion selection can introduce variability to peptide identification outputs, hinder quantification between sample runs and thus necessitate lengthy and costly procedures such as sample fractionation (to reduce input complexity) and injection replicates [1-3]. The more recent adoption of Data-Independent Acquisition (DIA) MS modes such as SWATH (Sequential Window Acquisition of all Theoretical fragment ion spectra), whereby MS2 fragment ion spectra are collected for each parent ion observed in a series of mass-to-charge isolation windows, has presented the opportunity to overcome these issues and obtain deep, label-free proteomic coverage of complex samples in a timely manner without fractionation [3-6]. Interpretation of the SWATH MS2 spectra however, requires reference to a spectral library of peptide sequence matching (including established m/z and LC retention time coordinates) itself obtained from the outcomes of multiple DDA runs. Spectral libraries of notable depth are available for only a few species (or specialised cells/tissues) -including human [7], mouse [8-9], a few microbiota [10-11], drosophila and tomato [12], zebrafish [13] and yeast [4] – that, at present, limits the breadth of uptake of SWATH and the application of its benefits for label-free proteome analyses.

The guinea pig is an excellent experimental model for many aspects of human physiology and pathophysiology - including maternal and fetal adaptations to pregnancy [14-17] cardiac excitation-contraction coupling [18], asthma and airway drug responsiveness [19-20], auditory somatosensory processes [21], type 2 diabetes [22] and vitamin C deficiency [23] – yet there is limited experimental information regarding the proteome available for this species. In an effort to overcome this obstacle, we sought to generate, via numerous LC-MS/MS measurements, a spectral library of the guinea pig proteome.

## Materials and methods

The overall experimental workflow is displayed in Figure 1. Homogenates were prepared from 16 tissues (brain, colon, duodenum, adipose, kidney, large intestine, liver, lung, ovaries, pancreas, placenta, skeletal muscle, small intestine, stomach, heart, uterus) isolated from guinea pigs (fetal- and adult) sacrificed according to the Animals (Scientific Procedures) Act 1986 under UK Home Office project license approval (PPL 60/4312). The study was approved by Newcastle University ethics review process. These homogenates were trypsin digested and subjected to >200 LC-MS/MS runs, utilizing a range of LC/MS platforms (i.e. Q-Exactive, TripleTOF 6600), chromatographic gradients and additional pre-processing steps (i.e. SDS page fractionation, acetylation enrichment). For more details please refer to Supplementary Table 1.

**Figure 1.**
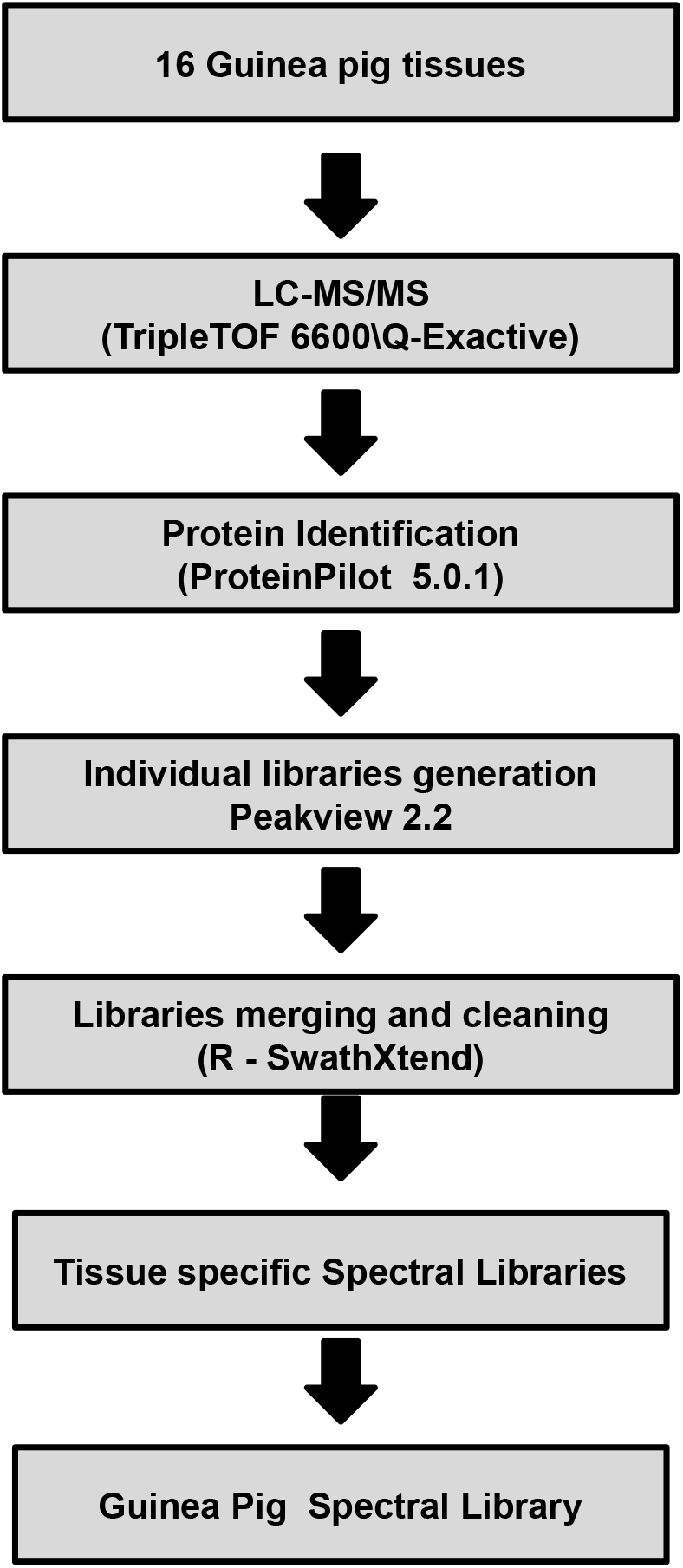
Schematic of the experimental workflow for spectral library generation.

The acquired MS/MS data were searched against the Uniprot guinea pig proteome (version: January 2016) re-annotated by combining the original annotation (if present) and annotation of homologous sequences from BLAST (version 2.2.30), searching against Swiss-Prot mammalian sequences. Consistency of annotation (i.e. elimination of synonyms) was achieved by mapping to the HGNC database [24].

The Q-Exactive *.raw files were first converted to *.mgf format using MSConvert (ProteoWizard package). TripleTOF 6600 *.wiff files were searched directly using Protein Pilot 5 (parameters used: cysteine alkylation: iodoacetamide, digestion enzyme: trypsin, search effort: thorough, instrument: TripleTOF 6600/ Orbi MS, Orbi MS/MS, default settings). Joint searches were performed for LC-MS runs of the same tissue, analysed within the same experimental batch (same instrument and setup, acquired the same day) and showing the same peptide retention time profile (assed by visual inspection).

The individual search results were exported (using PeakView 2.1), in a spectral library format, as *.tsv files and sequentially joined into 16 tissue-specific libraries using SwathXtend R package. Prior to joining, the libraries were cleaned to only contain unmodified peptides identified with FDR<0.01 with at least 5 corresponding fragment ions present. The confidence cut-off representative to FDR<0.01 was extracted from and applied individually to each search result file. At each joining step the retention times of the base and the add-on library were aligned (Figure 2A) and the correctness (linearity) of the alignment was visually inspected. On occasions, when the RT correlation was non-linear, the gradient was manually divided into linear fragments, which were pre-aligned, reassembled and then submitted to the joining algorithm, while fragments of poor correlation were discarded (Figure 2B and 2C). For final progression to inclusion in the concatenation steps, only correlation R^2^ values >0.9 were deemed acceptable (the actual range being 0.94-1, typically >0.98). The assembly of the total consensus spectral library was achieved analogically, by one-by-one joining of the individual tissue-specific spectral libraries to the base spectral library (starting with brain). All the steps of library assembly were carried out using R scripts which are attached as supplementary data.

**Figure 2.**
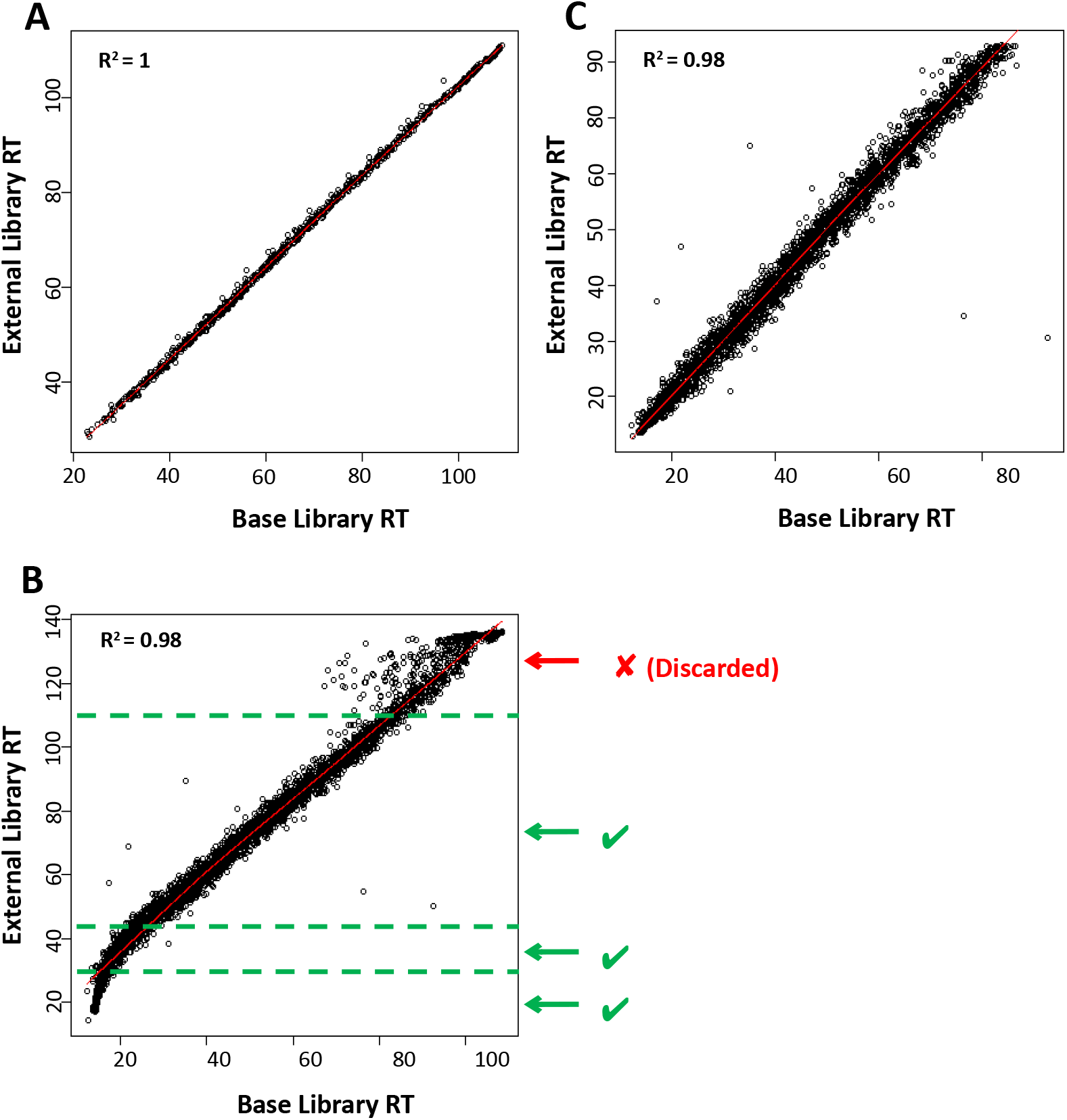
Concatenation of tissue-specific spectral libraries. A, liquid chromatography peptide retention time correlations between tissue-specific libraries. Panel A indicates excellent retention time correlation. Panel B indicates a situation where additional linearization was required. The external Library was manually divided into 4 parts, 3 of which are the linear fragments of the plot and the 4^th^, noisy fragment, which was discarded. For each of the 3 linear fragments, the linear regression was calculated and the resulting parameters were used to adjust peptide retention times to match the base library. Subsequently, the fragments with corrected retention times were combined and used for library building. Panel C is showing the outcome following normalization.

## Results and discussion

Analysis of >250,000 peptide-spectrum matches resulted in the construction of a library of 73594 peptides (unique to individual proteins) that corresponded to 7667 proteins. The majority of proteins were identified with more than one peptide (Figure 3A) and a minimum of 5 fragment ions per peptide (Figure 3B). The contribution of tissue-specific libraries to the total library varied roughly in accordance to the number of peptides, reflecting the biological properties, the number of biological replicates and repeat injections and the level of fractionation that was carried out for different tissues (directed by the core research interests of our group) (Figure 3C). The overlap between tissue-specific libraries is shown in Supplementary Figure 1, alongside normalized peptide counts plotted for proteins shared between the individual libraries, providing an indication of how similar/dissimilar different tissues are in terms of protein composition. Peptide retention time correlations and the corresponding correlation residuals plots are provided as Supplementary Figure 2.

**Figure 3.**
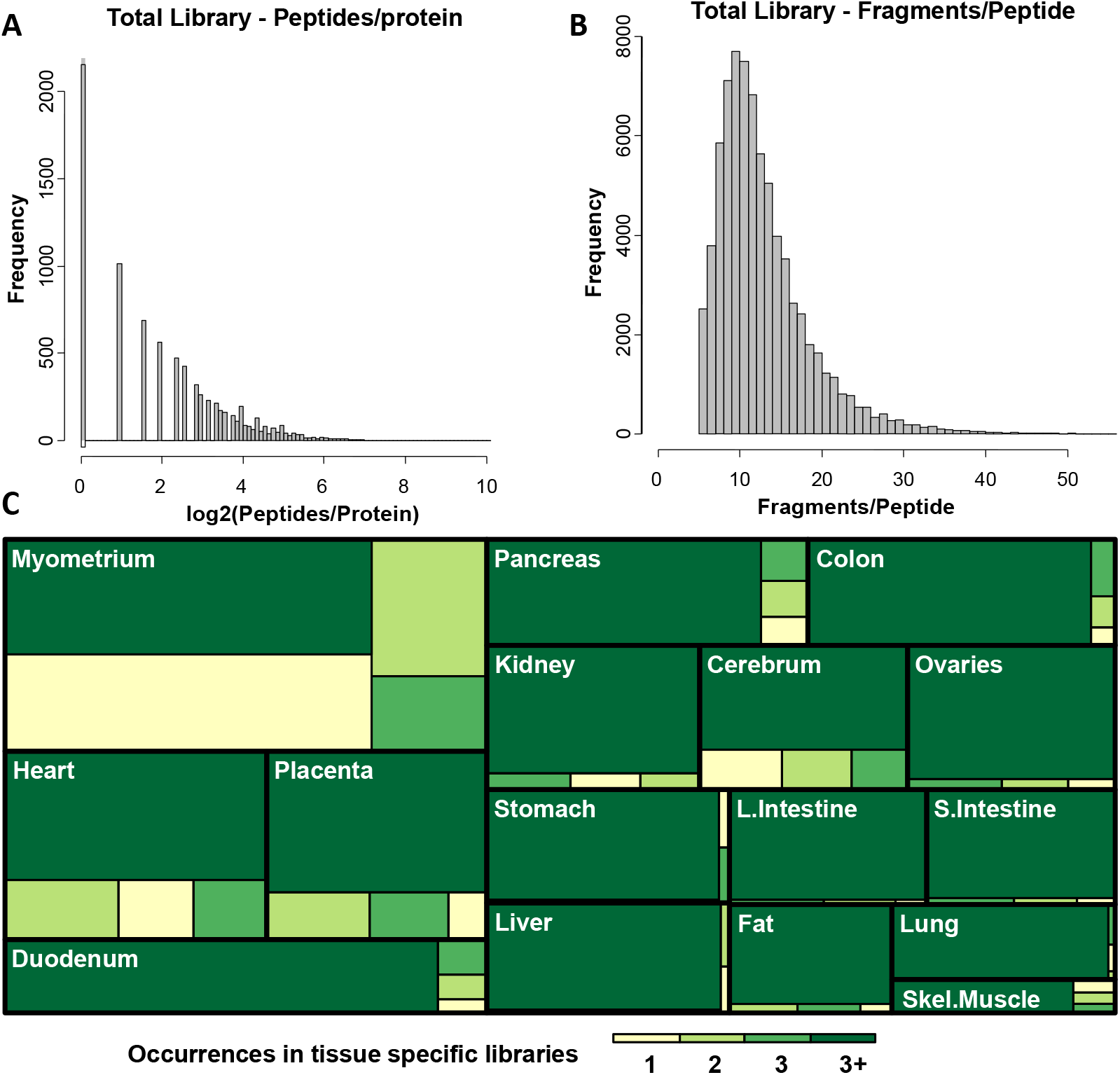
Summaries of library composition. Histograms indicating the number of peptides per protein (panel A) and the frequency distributions of fragment ions per peptide (panel B). Panel C, tissue-specific library contributions to the total library. Different colours indicate what proportion of the library is shared between multiple tissues (from 1 to 3+, 1 being unique to one tissue only and 3+ being found in more than 3 tissues).

The applicability of the total library for analysis of SWATH data was investigated by comparing the results of three biological replicates of guinea pig heart tissue digests (1μg protein per sample) analysed with: (i) Q-Exactive operating in DDA mode (UltiMate 3000 RSLCnano, 300μm x 5mm C18 PepMap C18 trap cartridge, 75μmx50cm PepMap C18 Easy-Spray column, 120min. gradient :8-30% acetonitrile, 300nl/min, Top10 HCD method) or (ii) SWATH on TripleTOF 6600 (Eksigent ekspert nanoLC425 HPLC, 20×0.18mm 2g-v/v 5μm C18 trap column, Waters NanoAcquity BSS T3 column, 1.8μm 250×0.075mm, 120min. gradient: 5-40% acetonitrile, 300nl/min., SWATH acquisition range of 400-1000m/z, with 27 SWATH bins).

The SWATH result files were processed in PeakView 2.1 (AB Sciex) with the SWATH MSMSall microapp, (top 1000 peptides/protein, top 5 transitions/peptide, confidence threshold: 90%, FDR <1%, modified peptides excluded, extraction window: 10min, XIC width: 10ppm).

The DDA output was analysed in MaxQuant 1.5.3.8 (top 5 unique peptides, max. missed cleavages: 2, non-modified or methionine oxidized peptides only, FDR <1%).

The comparison between SWATH and DDA (Supplementary Figure 3) show similar capability of each method in detecting proteins in a complex mixture such as heart that has a particularly high dynamic range. However, the potential of SWATH is revealed when used for quantification. Nearly all detected proteins could be quantified in each sample without missing peptide values, which in contrast is a significant problem for DDA-type experiments where a protein in only one sample (of the three) is enough for detection but would be insufficient for ant quantification between conditions.

In summary, we have generated the first spectral library of the guinea pig proteome for interrogation by SWATH-MS. This library contains peptide precursor and fragment ion information of 7667 guinea pig proteins. It greatly increases the validated guinea pig proteome information beyond that listed in Uniprot (September 2018). The freely accessible library will furnish the research community with a comprehensive proteomic resource to (i) explore future molecular-phenotypic studies using (re-engaging) the guinea pig as an experimental model and (ii) assess cross-species proteomic responses to consistent physiological and pathophysiological experimental challenges.

## Supporting information

R script 1

R script 2

R script 3

R script 4

## Acknowledgements

Supported by MRC (MR/L0009560/1) and the PRIME-XS project (Grant Agreement Number 262067, funded by the European Union Seventh Framework Program)

---

**Supplementary Figure 1. Tissue-specific library overlaps in protein and peptides.**

The overlap of identified proteins between individual tissue-specific libraries (above diagonal) contributing to the total library and the number of peptides per protein shared (below diagonal) are displayed. Tissues analysed in this study with corresponding number of LC-MS runs included in the spectral library are indicated on the diagonal.

**Supplementary Figure 2. Retention time correlation\ residual plot.**

The peptide retention time correlation between individual tissue specific libraries (above diagonal) and corresponding residual plots (below diagonal)

**Supplementary Figure 3. Comparison of DDA and SWATH**

A). Comparison between SWATH and DDA analysis of three guinea pig hearts. The clear advantage of SWATH is in a very low number of missing values for peptides/proteins between samples. The Number of identified proteins is similar with nearly 70% overlap (B). Quantitative results are very similar as shown on PCA plot (C) and with hierarchical clustering (D).

**Supplementary Table 1. Summary experimental procedures and data analysis.**

The details of individual sample processing procedures, LC-MS methods and instrumentation used, together with MS-MS searching information (searching batch composition)

**Supplementary data.**

Supplementary Figures 1 and 2

Supplementary table 1

R scripts (also available through MassIVE -MSV000083199):

heart library.R

individual_tissue_specific.R

myo_library.R

tissue specific combined.R

## References

[1] Domon B, Aebersold R. Nature Biotechnology. 2010; 28:710–721.

[2] Mallick P, Kuster B. Nature Biotechnology. 2010; 28:695–710.

[3] Wu JX, Song X, Pascovici D, Zaw T, Care N, Krisp C, Molloy MP. Molecular and Cellular Proteomics. 2016; 15:2501–2514.

[4] Schubert OT, Gillet LC, Collins BC, Navarro P, Rosenberger G, Wolski WE, Lam H, Amodei D, Mallick P, MacLean B, Aebersold R. Nature Protocols 2015; 10:426–441.

[5] G. Rosenberger, I. Bludau, U. Schmitt, M. Heusel, C. Hunter, Y. Liu, M.J. MacCoss, B.X. MacLean, A.I. Nesvizhskii, P.G.A. Pedriolo, L. Reiter, H.L. Rost, S. Tate, Y.S. Ting, B.C. Collins, R. Aebersold, Nat Methods 2017, 14, 921–927.

[6] B.C. Collins, C.L. Hunter, Y. Liu, B. Schilling, G. Rosenberger, S.L. Bader, D.W. Chan, B.W. Gibson, A. Gingras, J.M. Held, M. Hirayama-Kurogi, H. Hou, C. Krisp, B. Larsen, L. Lin, S. Liu, M.P. Molloy, R.L. Moritz, S. Ohtsuki, R. Schlapbach, N. Selevsek, S.N. Thomas, S. Tzeng, H. Zhang, R. Aebersold, Nat Commun 2017, 8, 291.

[7] G. Rosenberger, C.C. Koh, T. Guo, H.L. Rost, P. Kouvonen, B.C. Collins, M. Heusel, Y. Liu, E. Caron, A. Vichalkovski, M. Faini, O.T. Schubert, P. Faridi, H.A. Ebhardt, M. Matondo, H. Lam, S.L. Bader, D.S. Campbell, E.W. Deutsch, R.L. Moritz, S. Tate, R. Aebersold, Scientific Data. 2014, 1, 140031.

[8] E. Malstrom, O. Kilsgard, S. Hauri, E. Smeds, H. Herwals, L. Malmstrom, J, Malmstrom, Nat. Commun 2016, 7, 10261.

[9] C. Braccia, M. Pons Espinal, M. Pini, D. Pietri Tonelli, A. Amirotti A, Data in Brief 2018, 18, 1–8.

[10] O.T. Schubert, C. Ludwig, M. Kogadeeva, M. Zimmermann, G. Rosenberger, M. Gengenbacher, L.C. Gillet, B.C. Collins, H.L. Rost, S.H.E. Kaufmann, U. Sauer, R. Aebersold. Cell Host & Microbe 2015, 18, P96–P108.

[11] D.B. Muller, O.T. Schubert, H. Rost, R. Aebersold, J.A. Vorholt. Mol. Cell. Proteomics 2016, 15, 3256.

[12] B. Fabre, D. Korona, C.I. Mata, H.T. Parsons, M.J. Deery, M.L.A.T.M. Hertog, B.M. Nicolai, S. Russell, K.S. Lilley. Proteomics 2017, 17, 1700216.

[13] S.J. van der Plas-Duivesteijn, Y. Mohammed, H. Dalebout, A. Meijer, A. Botermans, J.L. Hoogendijk, A.A. Henneman, A.M. Deelder, H.P. Spaink, M. Palmblad. J Proteome Res 2014, 13, 1537–44.

[14] M.J. Taggart, B.F. Mitchell. Am J Physiol 2009, 297, R525–545.

[15] M.J. Taggart, Hume R, Lartey J, Johnson M, Cardiovasc Res. 2014.104: 226–227.

[16] N. Gomez-Lopez, W.C. Tong, M. Arenas-Hernandez, S. Tanaka, O. Hajar, D.M. Olson, M.J. Taggart, B.F. Mitchell. Am J Reprod Immunol 2015, 73, 341–52.

[17] J.L. Morrison, K.J. Botting, J.R. T. Darby, A.L. David, R.M. Dyson, K.L. Gatford, C. Gray, E.A. Herrera, J.J. Hirst, B. Kim8, K.L. Kind, B.J. Krause, S.G. Matthews, H.K. Palliser, T.R. H. Regnault, B.S. Richardson, A. Sasaki8, L.P. Thompson, M.J. Berry. 2018, J Physiol 1–35.

[18] D. Bers. Nature 2002, 415, 198.

[19] F.L.M. Ricciardolo, F. Nijkamp, V. De Rose, G. Folkerts, Curr Drug Targets 2008, 9, 452.

[20] A.J. Lemond, S. Keir, S. Terakosolphan, C.P. Page, B. Forbes. Drug Deliv Transl Res 2018, 8: 760–769.

[21] K.L. Marks, D.T Martel, C. Wu, G.J. Basura,. Sci Transl Med 2018, 10, eaal3175

[22] B.K. Podell, D.F. Ackart, M.A. Richardson, J.E. DiLisio, B. Pulford, R.J. Basaraba. Disease Models & Mechanisms 2017, 10, 151.

[23] D. Søgaard, M.M. Lindblad, M.D. Paidi, S. Hasselholt, J. Lykkesfeldt, P. Tveden-Nyborg. Nutr Res 34, 639.

[24] B. Yates, B. Braschi, K.A. Gray, R.L. Seal, S. Tweedie, E.A. Bruford. Nucleic Acids Res, 45, D619–25.

