## Supplementary figures and images for "The generation of a comprehensive spectral library for the analysis of the guinea pig proteome by SWATH-MS"

### Supplementary figure 1

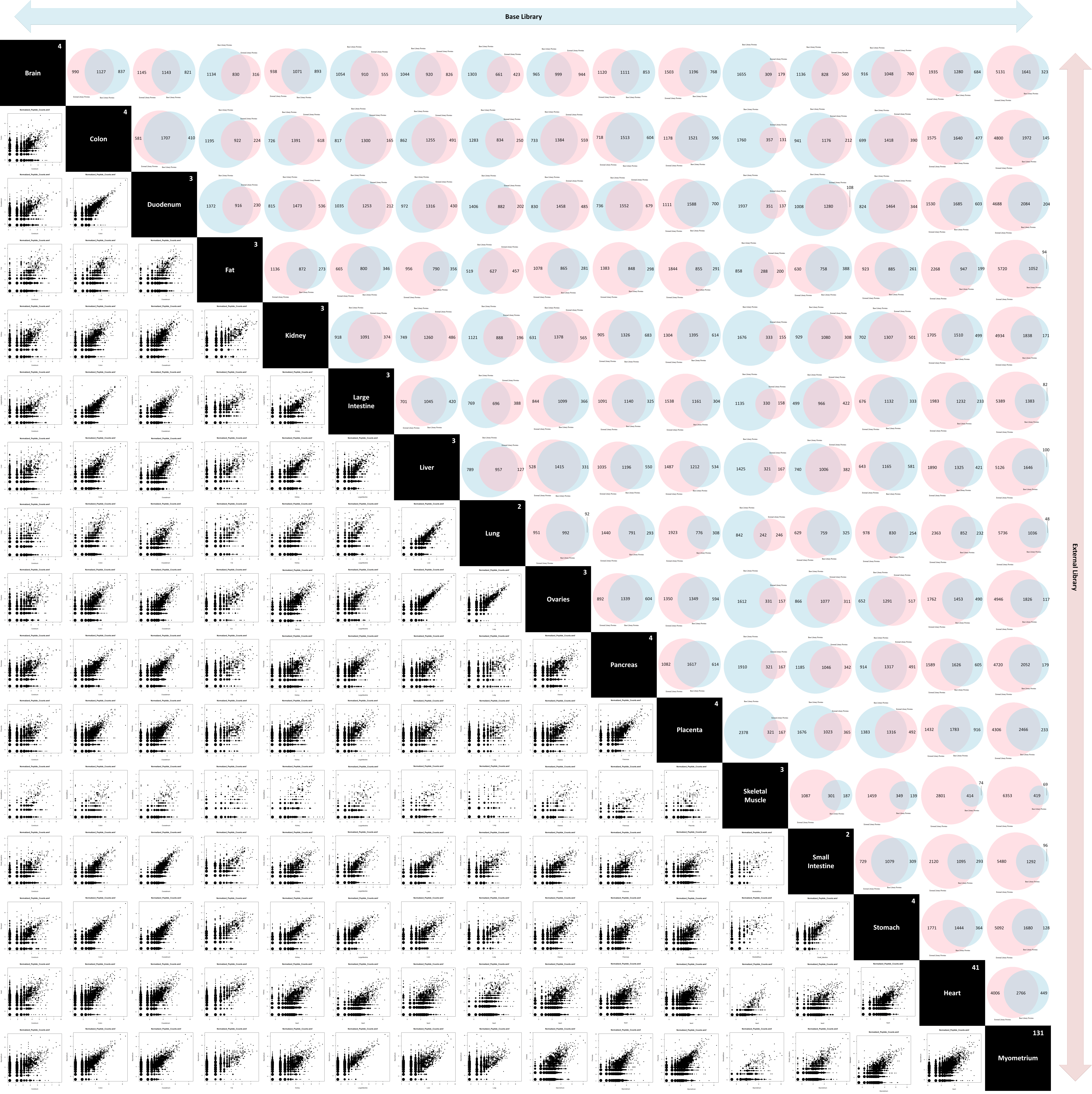

### Supplementary figure 2

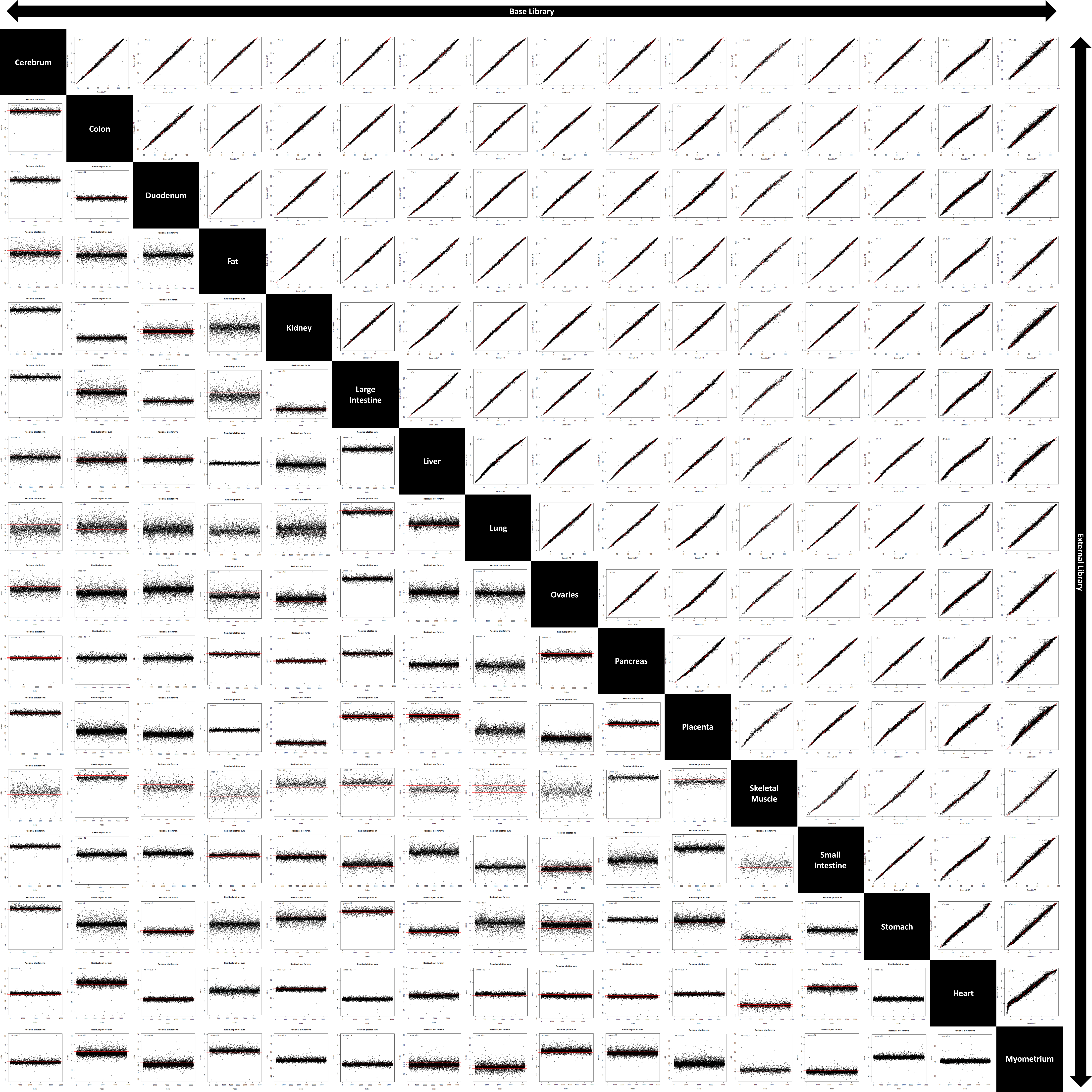

### Supplementary figure 3

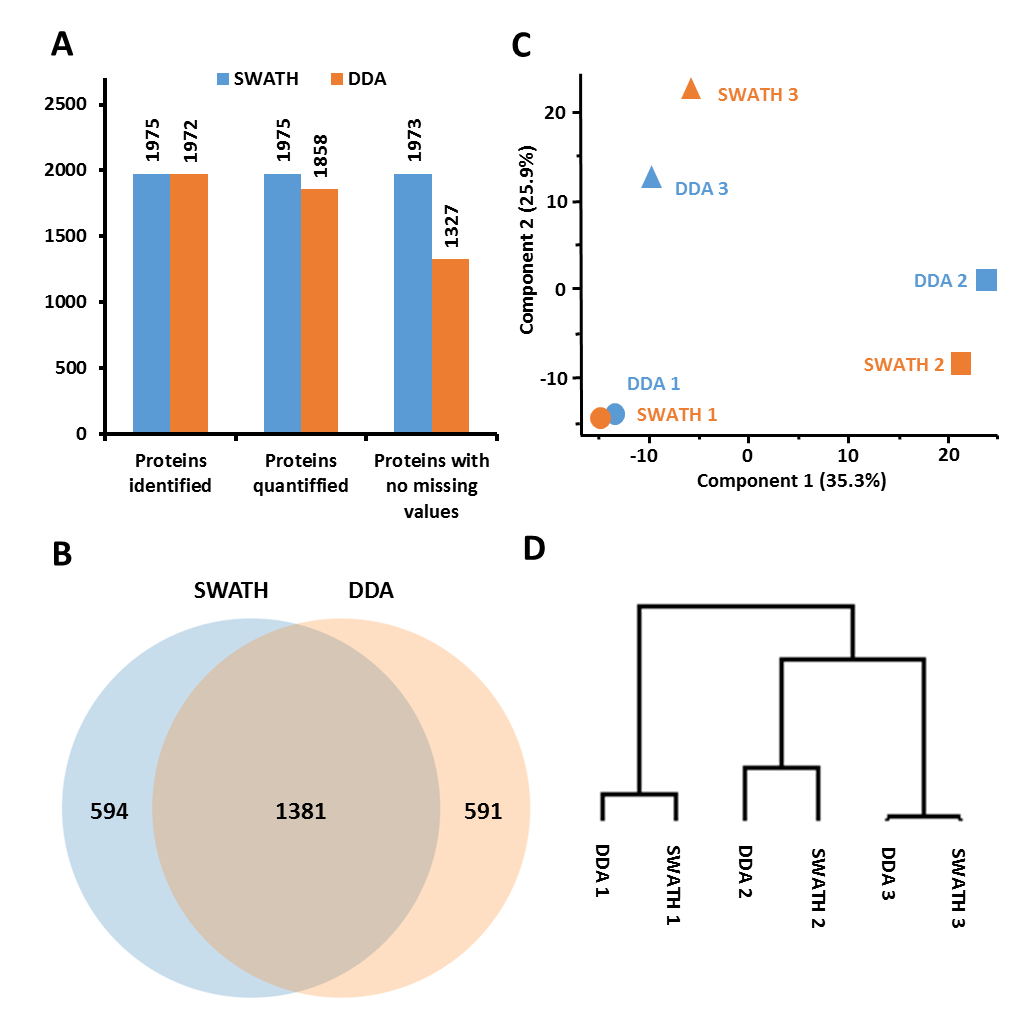
