## Supplementary material for "The generation of a comprehensive spectral library for the analysis of the guinea pig proteome by SWATH-MS": R script 1

```
#Individual search results (.group files) were converted into spectral library format Using PeakView software package with SWATH microApp and saved as .txt

#Tissue specific libraries are being assembled here using multiple idividual search results.

library(SwathXtend)
library(plyr)
library(openxlsx)

workDir<-getwd()

# Pancreas

#1
#Retrieving the conf.cutoff, corresponding to FDR<0.01, from the search summarry in .xlsx format

conf1<-read.xlsx("170303_02_PP_GP_Pancreas__FDR.xlsx", sheet = "Search Summary",
                 colNames = FALSE, rows=33)[1,4]
conf2<-read.xlsx("170310_02_PP_GP_Pancreas__FDR.xlsx", sheet = "Search Summary",
                 colNames = FALSE, rows=33)[1,4]
#Loading and cleaning the library files
lib1<-readLibFile("170303_02_PP_GP_Pancreas.txt",clean = TRUE,nomod=TRUE,nomc=FALSE, conf.cutoff=conf1)
lib2<-readLibFile("170310_02_PP_GP_Pancreas.txt",clean = TRUE,nomod=TRUE,nomc=FALSE, conf.cutoff=conf2)
#correcting the file format (removing the retension time calibrant and nonannotated peptides)
lib1<-canonicalFormat(lib1, format = "peakview")
lib2<-canonicalFormat(lib2, format = "peakview")
lib1<-lib1[lib1$uniprot_id!="[ RT-Cal protein ]",]
lib1<-lib1[lib1$uniprot_id!="1/[ RT-Cal protein ]",]
lib2<-lib2[lib2$uniprot_id!="[ RT-Cal protein ]",]
lib2<-lib2[lib2$uniprot_id!="1/[ RT-Cal protein ]",]
lib1<-lib1[!grepl("NA",rownames(lib1)),]
lib2<-lib2[!grepl("NA",rownames(lib2)),]


# Checking the linearity of RT correlation. If nonlinear, the gradient is divided to linear fragments. 

checkQuality(lib1,lib2)

#based on the visual inspection, the retension time correlation is non-linear
#The External library lib2 (the one that is being added to the "base library" lib1) is therefore manually divided into few, more linear fragments (lib2a-lib2c)

#Linear fragment one

lib2a<-lib2[lib2$RT_detected<60,]
RT1<-lib1[,c(1,3,8)]
RT2<-lib2a[,c(1,3,8)]
RT1<-RT1[RT1$modification_sequence%in%RT2$modification_sequence,]
RT2<-RT2[RT2$modification_sequence%in%RT1$modification_sequence,]
RT1<-RT1[!duplicated(RT1$modification_sequence),2:3]
RT2<-RT2[!duplicated(RT2$modification_sequence),2:3]
colnames(RT2)[1]<-"RT_detected_2"
RT<-join(RT1,RT2)

# Linear regression is calculated and the coeficients used to correct the external library RT to mach the base library

fit <- coef(lm(RT$RT_detected ~ RT$RT_detected_2))
lib2a$RT_detected<-(lib2a$RT_detected*fit[2])+fit[1]
lib2a$iRT<-lib2a$RT_detected

#Linear fragment 2

lib2b<-lib2[lib2$RT_detected>=60,]
lib2b<-lib2b[lib2b$RT_detected<96,]
RT1<-lib1[,c(1,3,8)]
RT2<-lib2b[,c(1,3,8)]
RT1<-RT1[RT1$modification_sequence%in%RT2$modification_sequence,]
RT2<-RT2[RT2$modification_sequence%in%RT1$modification_sequence,]
RT1<-RT1[!duplicated(RT1$modification_sequence),2:3]
RT2<-RT2[!duplicated(RT2$modification_sequence),2:3]
colnames(RT2)[1]<-"RT_detected_2"
RT<-join(RT1,RT2)
fit <- coef(lm(RT$RT_detected ~ RT$RT_detected_2))
lib2b$RT_detected<-(lib2b$RT_detected*fit[2])+fit[1]
lib2b$iRT<-lib2b$RT_detected

#Linear fragment 3

lib2c<-lib2[lib2$RT_detected>=95,]
RT1<-lib1[,c(1,3,8)]
RT2<-lib2c[,c(1,3,8)]
RT1<-RT1[RT1$modification_sequence%in%RT2$modification_sequence,]
RT2<-RT2[RT2$modification_sequence%in%RT1$modification_sequence,]
RT1<-RT1[!duplicated(RT1$modification_sequence),2:3]
RT2<-RT2[!duplicated(RT2$modification_sequence),2:3]
colnames(RT2)[1]<-"RT_detected_2"
RT<-join(RT1,RT2)
fit <- coef(lm(RT$RT_detected ~ RT$RT_detected_2))
lib2c$RT_detected<-(lib2c$RT_detected*fit[2])+fit[1]
lib2c$iRT<-lib2c$RT_detected

#Linear fragments are merged together 

lib2<-rbind(lib2a,lib2b,lib2c)

#Linear fragments are then reassembled  

buildSpectraLibPair(lib1, lib2, outputFile = "ExtendedLib_Pancreas.txt", plot = TRUE,
                    clean = FALSE, merge = TRUE, parseAcc = TRUE, consolidateAccession = TRUE,
                    nomod=TRUE, conf.cutoff=0.9)


#2
#The process described above is then repeated untill all the partial libraries are joined.
#In the followinf steps, only the external library needs to be cleaned.

conf2<-read.xlsx("170324_07_PP_GPlibrary_Pancreas__FDR.xlsx", sheet = "Search Summary",
                 colNames = FALSE, rows=33)[1,4]
lib1<-readLibFile("ExtendedLib_Pancreas.txt", clean=FALSE)
lib2<-readLibFile("170324_07_PP_GPlibrary_Pancreas.txt",clean=TRUE,nomod=TRUE,nomc=FALSE, conf.cutoff=conf2)
lib2<-canonicalFormat(lib2, format = "peakview")
lib2<-lib2[lib2$uniprot_id!="[ RT-Cal protein ]",]
lib2<-lib2[lib2$uniprot_id!="1/[ RT-Cal protein ]",]
lib1<-lib1[!grepl("NA",rownames(lib1)),]
lib2<-lib2[!grepl("NA",rownames(lib2)),]
lib1$iRT<-lib1$RT_detected

# checkQuality(lib1,lib2)

lib2a<-lib2[lib2$RT_detected<87,]
RT1<-lib1[,c(1,3,8)]
RT2<-lib2a[,c(1,3,8)]
RT1<-RT1[RT1$modification_sequence%in%RT2$modification_sequence,]
RT2<-RT2[RT2$modification_sequence%in%RT1$modification_sequence,]
RT1<-RT1[!duplicated(RT1$modification_sequence),2:3]
RT2<-RT2[!duplicated(RT2$modification_sequence),2:3]
colnames(RT2)[1]<-"RT_detected_2"
RT<-join(RT1,RT2)
fit <- coef(lm(RT$RT_detected ~ RT$RT_detected_2))
lib2a$RT_detected<-(lib2a$RT_detected*fit[2])+fit[1]
lib2a$iRT<-lib2a$RT_detected

lib2b<-lib2[lib2$RT_detected>=86,]
RT1<-lib1[,c(1,3,8)]
RT2<-lib2b[,c(1,3,8)]
RT1<-RT1[RT1$modification_sequence%in%RT2$modification_sequence,]
RT2<-RT2[RT2$modification_sequence%in%RT1$modification_sequence,]
RT1<-RT1[!duplicated(RT1$modification_sequence),2:3]
RT2<-RT2[!duplicated(RT2$modification_sequence),2:3]
colnames(RT2)[1]<-"RT_detected_2"
RT<-join(RT1,RT2)
fit <- coef(lm(RT$RT_detected ~ RT$RT_detected_2))
lib2b$RT_detected<-(lib2b$RT_detected*fit[2])+fit[1]
lib2b$iRT<-lib2b$RT_detected

lib2<-rbind(lib2a,lib2b)

buildSpectraLibPair(lib1, lib2, outputFile = "ExtendedLib_Pancreas.txt", plot = TRUE,
                    clean = FALSE, merge = TRUE, parseAcc = TRUE, consolidateAccession = TRUE,
                    nomod=TRUE, conf.cutoff=0.9)

#3

conf2<-read.xlsx("170324_07_PP_GPlibrary_Pancreas1__FDR.xlsx", sheet = "Search Summary",
                 colNames = FALSE, rows=33)[1,4]
lib1<-readLibFile("ExtendedLib_Pancreas.txt",clean=FALSE)
lib2<-readLibFile("170324_07_PP_GPlibrary_Pancreas1.txt",clean=TRUE,nomod=TRUE,nomc=FALSE,conf.cutoff=conf2)
lib2<-canonicalFormat(lib2, format = "peakview")
lib2<-lib2[lib2$uniprot_id!="[ RT-Cal protein ]",]
lib2<-lib2[lib2$uniprot_id!="1/[ RT-Cal protein ]",]
lib1<-lib1[!grepl("NA",rownames(lib1)),]
lib2<-lib2[!grepl("NA",rownames(lib2)),]
lib1$iRT<-lib1$RT_detected

# checkQuality(lib1,lib2)

lib2a<-lib2[lib2$RT_detected<87,]
RT1<-lib1[,c(1,3,8)]
RT2<-lib2a[,c(1,3,8)]
RT1<-RT1[RT1$modification_sequence%in%RT2$modification_sequence,]
RT2<-RT2[RT2$modification_sequence%in%RT1$modification_sequence,]
RT1<-RT1[!duplicated(RT1$modification_sequence),2:3]
RT2<-RT2[!duplicated(RT2$modification_sequence),2:3]
colnames(RT2)[1]<-"RT_detected_2"
RT<-join(RT1,RT2)
fit <- coef(lm(RT$RT_detected ~ RT$RT_detected_2))
lib2a$RT_detected<-(lib2a$RT_detected*fit[2])+fit[1]
lib2a$iRT<-lib2a$RT_detected

lib2b<-lib2[lib2$RT_detected>=86,]
RT1<-lib1[,c(1,3,8)]
RT2<-lib2b[,c(1,3,8)]
RT1<-RT1[RT1$modification_sequence%in%RT2$modification_sequence,]
RT2<-RT2[RT2$modification_sequence%in%RT1$modification_sequence,]
RT1<-RT1[!duplicated(RT1$modification_sequence),2:3]
RT2<-RT2[!duplicated(RT2$modification_sequence),2:3]
colnames(RT2)[1]<-"RT_detected_2"
RT<-join(RT1,RT2)
fit <- coef(lm(RT$RT_detected ~ RT$RT_detected_2))
lib2b$RT_detected<-(lib2b$RT_detected*fit[2])+fit[1]
lib2b$iRT<-lib2b$RT_detected

lib2<-rbind(lib2a,lib2b)

buildSpectraLibPair(lib1, lib2, outputFile = "ExtendedLib_Pancreas.txt", plot = TRUE,
                    clean = FALSE, merge = TRUE, parseAcc = TRUE, consolidateAccession = TRUE,
                    nomod=TRUE, conf.cutoff=0.9)

#The final library is now being loaded to remove peptide entries with less then 5 fragment ions
x<-read.delim("ExtendedLib_Pancreas.txt")
y<-as.data.frame(table(x$stripped_sequence))
y<-y[y$Freq>=5,]
x<-x[x$stripped_sequence%in%y$Var1,]
write.table(x,"ExtendedLib_Pancreas.txt",sep="\t")


#The whole process is now repeated for other tissues

# Duodenum

#1

conf1<-read.xlsx("170303_03_PP_GP_Duoadenum__FDR.xlsx", sheet = "Search Summary",
                 colNames = FALSE, rows=33)[1,4]
conf2<-read.xlsx("170310_03_PP_GP_Duoadenum__FDR.xlsx", sheet = "Search Summary",
                 colNames = FALSE, rows=33)[1,4]
lib1<-readLibFile("170303_03_PP_GP_Duoadenum.txt",clean = TRUE,nomod=TRUE,nomc=FALSE, conf.cutoff=conf1)
lib2<-readLibFile("170310_03_PP_GP_Duoadenum.txt",clean = TRUE,nomod=TRUE,nomc=FALSE, conf.cutoff=conf2)
lib1<-canonicalFormat(lib1, format = "peakview")
lib2<-canonicalFormat(lib2, format = "peakview")
lib1<-lib1[lib1$uniprot_id!="[ RT-Cal protein ]",]
lib1<-lib1[lib1$uniprot_id!="1/[ RT-Cal protein ]",]
lib2<-lib2[lib2$uniprot_id!="[ RT-Cal protein ]",]
lib2<-lib2[lib2$uniprot_id!="1/[ RT-Cal protein ]",]
lib1<-lib1[!grepl("NA",rownames(lib1)),]
lib2<-lib2[!grepl("NA",rownames(lib2)),]

# checkQuality(lib1,lib2)

lib2a<-lib2[lib2$RT_detected<60,]
RT1<-lib1[,c(1,3,8)]
RT2<-lib2a[,c(1,3,8)]
RT1<-RT1[RT1$modification_sequence%in%RT2$modification_sequence,]
RT2<-RT2[RT2$modification_sequence%in%RT1$modification_sequence,]
RT1<-RT1[!duplicated(RT1$modification_sequence),2:3]
RT2<-RT2[!duplicated(RT2$modification_sequence),2:3]
colnames(RT2)[1]<-"RT_detected_2"
RT<-join(RT1,RT2)
fit <- coef(lm(RT$RT_detected ~ RT$RT_detected_2))
lib2a$RT_detected<-(lib2a$RT_detected*fit[2])+fit[1]
lib2a$iRT<-lib2a$RT_detected

lib2b<-lib2[lib2$RT_detected>=60,]
lib2b<-lib2b[lib2b$RT_detected<96,]
RT1<-lib1[,c(1,3,8)]
RT2<-lib2b[,c(1,3,8)]
RT1<-RT1[RT1$modification_sequence%in%RT2$modification_sequence,]
RT2<-RT2[RT2$modification_sequence%in%RT1$modification_sequence,]
RT1<-RT1[!duplicated(RT1$modification_sequence),2:3]
RT2<-RT2[!duplicated(RT2$modification_sequence),2:3]
colnames(RT2)[1]<-"RT_detected_2"
RT<-join(RT1,RT2)
fit <- coef(lm(RT$RT_detected ~ RT$RT_detected_2))
lib2b$RT_detected<-(lib2b$RT_detected*fit[2])+fit[1]
lib2b$iRT<-lib2b$RT_detected

lib2c<-lib2[lib2$RT_detected>=95,]
RT1<-lib1[,c(1,3,8)]
RT2<-lib2c[,c(1,3,8)]
RT1<-RT1[RT1$modification_sequence%in%RT2$modification_sequence,]
RT2<-RT2[RT2$modification_sequence%in%RT1$modification_sequence,]
RT1<-RT1[!duplicated(RT1$modification_sequence),2:3]
RT2<-RT2[!duplicated(RT2$modification_sequence),2:3]
colnames(RT2)[1]<-"RT_detected_2"
RT<-join(RT1,RT2)
fit <- coef(lm(RT$RT_detected ~ RT$RT_detected_2))
lib2c$RT_detected<-(lib2c$RT_detected*fit[2])+fit[1]
lib2c$iRT<-lib2c$RT_detected

lib2<-rbind(lib2a,lib2b,lib2c)

buildSpectraLibPair(lib1, lib2, outputFile = "ExtendedLib_Duoadenum.txt", plot = TRUE,
                    clean = FALSE, merge = TRUE, parseAcc = TRUE, consolidateAccession = TRUE,
                    nomod=TRUE, conf.cutoff=0.9)
#2

conf2<-read.xlsx("170324_08_PP_GPlibrary_Duodenum__FDR.xlsx", sheet = "Search Summary",
                 colNames = FALSE, rows=33)[1,4]
lib1<-readLibFile("ExtendedLib_Duoadenum.txt",clean=FALSE)
lib2<-readLibFile("170324_08_PP_GPlibrary_Duodenum.txt",clean=TRUE,nomod=TRUE,nomc=FALSE, conf.cutoff=conf2)
lib2<-canonicalFormat(lib2, format = "peakview")
lib2<-lib2[lib2$uniprot_id!="[ RT-Cal protein ]",]
lib2<-lib2[lib2$uniprot_id!="1/[ RT-Cal protein ]",]
lib1<-lib1[!grepl("NA",rownames(lib1)),]
lib2<-lib2[!grepl("NA",rownames(lib2)),]
lib1$iRT<-lib1$RT_detected

# checkQuality(lib1,lib2)

lib2a<-lib2[lib2$RT_detected<87,]
RT1<-lib1[,c(1,3,8)]
RT2<-lib2a[,c(1,3,8)]
RT1<-RT1[RT1$modification_sequence%in%RT2$modification_sequence,]
RT2<-RT2[RT2$modification_sequence%in%RT1$modification_sequence,]
RT1<-RT1[!duplicated(RT1$modification_sequence),2:3]
RT2<-RT2[!duplicated(RT2$modification_sequence),2:3]
colnames(RT2)[1]<-"RT_detected_2"
RT<-join(RT1,RT2)
fit <- coef(lm(RT$RT_detected ~ RT$RT_detected_2))
lib2a$RT_detected<-(lib2a$RT_detected*fit[2])+fit[1]
lib2a$iRT<-lib2a$RT_detected

lib2b<-lib2[lib2$RT_detected>=86,]
RT1<-lib1[,c(1,3,8)]
RT2<-lib2b[,c(1,3,8)]
RT1<-RT1[RT1$modification_sequence%in%RT2$modification_sequence,]
RT2<-RT2[RT2$modification_sequence%in%RT1$modification_sequence,]
RT1<-RT1[!duplicated(RT1$modification_sequence),2:3]
RT2<-RT2[!duplicated(RT2$modification_sequence),2:3]
colnames(RT2)[1]<-"RT_detected_2"
RT<-join(RT1,RT2)
fit <- coef(lm(RT$RT_detected ~ RT$RT_detected_2))
lib2b$RT_detected<-(lib2b$RT_detected*fit[2])+fit[1]
lib2b$iRT<-lib2b$RT_detected

lib2<-rbind(lib2a,lib2b)

buildSpectraLibPair(lib1, lib2, outputFile = "ExtendedLib_Duoadenum.txt", plot = TRUE,
                    clean = FALSE, merge = TRUE, parseAcc = TRUE, consolidateAccession = TRUE,
                    nomod=TRUE, conf.cutoff=0.9)

x<-read.delim("ExtendedLib_Duoadenum.txt")
y<-as.data.frame(table(x$stripped_sequence))
y<-y[y$Freq>=5,]
x<-x[x$stripped_sequence%in%y$Var1,]
write.table(x,"ExtendedLib_Duoadenum.txt",sep="\t")


# Colon

#1
conf1<-read.xlsx("170303_04_PP_GP_Colon__FDR.xlsx", sheet = "Search Summary",
                 colNames = FALSE, rows=33)[1,4]
conf2<-read.xlsx("170310_04_PP_GP_Colon__FDR.xlsx", sheet = "Search Summary",
                 colNames = FALSE, rows=33)[1,4]
lib1<-readLibFile("170303_04_PP_GP_Colon.txt",clean = TRUE,nomod=TRUE,nomc=FALSE, conf.cutoff=conf1)
lib2<-readLibFile("170310_04_PP_GP_Colon.txt",clean = TRUE,nomod=TRUE,nomc=FALSE, conf.cutoff=conf2)
lib1<-canonicalFormat(lib1, format = "peakview")
lib2<-canonicalFormat(lib2, format = "peakview")
lib1<-lib1[lib1$uniprot_id!="[ RT-Cal protein ]",]
lib1<-lib1[lib1$uniprot_id!="1/[ RT-Cal protein ]",]
lib2<-lib2[lib2$uniprot_id!="[ RT-Cal protein ]",]
lib2<-lib2[lib2$uniprot_id!="1/[ RT-Cal protein ]",]
lib1<-lib1[!grepl("NA",rownames(lib1)),]
lib2<-lib2[!grepl("NA",rownames(lib2)),]

# checkQuality(lib1,lib2)

lib2a<-lib2[lib2$RT_detected<60,]
RT1<-lib1[,c(1,3,8)]
RT2<-lib2a[,c(1,3,8)]
RT1<-RT1[RT1$modification_sequence%in%RT2$modification_sequence,]
RT2<-RT2[RT2$modification_sequence%in%RT1$modification_sequence,]
RT1<-RT1[!duplicated(RT1$modification_sequence),2:3]
RT2<-RT2[!duplicated(RT2$modification_sequence),2:3]
colnames(RT2)[1]<-"RT_detected_2"
RT<-join(RT1,RT2)
fit <- coef(lm(RT$RT_detected ~ RT$RT_detected_2))
lib2a$RT_detected<-(lib2a$RT_detected*fit[2])+fit[1]
lib2a$iRT<-lib2a$RT_detected

lib2b<-lib2[lib2$RT_detected>=60,]
lib2b<-lib2b[lib2b$RT_detected<96,]
RT1<-lib1[,c(1,3,8)]
RT2<-lib2b[,c(1,3,8)]
RT1<-RT1[RT1$modification_sequence%in%RT2$modification_sequence,]
RT2<-RT2[RT2$modification_sequence%in%RT1$modification_sequence,]
RT1<-RT1[!duplicated(RT1$modification_sequence),2:3]
RT2<-RT2[!duplicated(RT2$modification_sequence),2:3]
colnames(RT2)[1]<-"RT_detected_2"
RT<-join(RT1,RT2)
fit <- coef(lm(RT$RT_detected ~ RT$RT_detected_2))
lib2b$RT_detected<-(lib2b$RT_detected*fit[2])+fit[1]
lib2b$iRT<-lib2b$RT_detected

lib2c<-lib2[lib2$RT_detected>=95,]
RT1<-lib1[,c(1,3,8)]
RT2<-lib2c[,c(1,3,8)]
RT1<-RT1[RT1$modification_sequence%in%RT2$modification_sequence,]
RT2<-RT2[RT2$modification_sequence%in%RT1$modification_sequence,]
RT1<-RT1[!duplicated(RT1$modification_sequence),2:3]
RT2<-RT2[!duplicated(RT2$modification_sequence),2:3]
colnames(RT2)[1]<-"RT_detected_2"
RT<-join(RT1,RT2)
fit <- coef(lm(RT$RT_detected ~ RT$RT_detected_2))
lib2c$RT_detected<-(lib2c$RT_detected*fit[2])+fit[1]
lib2c$iRT<-lib2c$RT_detected

lib2<-rbind(lib2a,lib2b,lib2c)

buildSpectraLibPair(lib1, lib2, outputFile = "ExtendedLib_Colon.txt", plot = TRUE,
                    clean = FALSE, merge = TRUE, parseAcc = TRUE, consolidateAccession = TRUE,
                    nomod=TRUE, conf.cutoff=0.9)
#2


conf2<-read.xlsx("170324_09_PP_GPlibrary_Colon__FDR.xlsx", sheet = "Search Summary",
                 colNames = FALSE, rows=33)[1,4]
lib1<-readLibFile("ExtendedLib_Colon.txt",clean=FALSE)
lib2<-readLibFile("170324_09_PP_GPlibrary_Colon.txt",clean = TRUE,nomod=TRUE,nomc=FALSE, conf.cutoff=conf2)
lib2<-canonicalFormat(lib2, format = "peakview")
lib2<-lib2[lib2$uniprot_id!="[ RT-Cal protein ]",]
lib2<-lib2[lib2$uniprot_id!="1/[ RT-Cal protein ]",]
lib1<-lib1[!grepl("NA",rownames(lib1)),]
lib2<-lib2[!grepl("NA",rownames(lib2)),]
lib1$iRT<-lib1$RT_detected

# checkQuality(lib1,lib2)

lib2a<-lib2[lib2$RT_detected<87,]
RT1<-lib1[,c(1,3,8)]
RT2<-lib2a[,c(1,3,8)]
RT1<-RT1[RT1$modification_sequence%in%RT2$modification_sequence,]
RT2<-RT2[RT2$modification_sequence%in%RT1$modification_sequence,]
RT1<-RT1[!duplicated(RT1$modification_sequence),2:3]
RT2<-RT2[!duplicated(RT2$modification_sequence),2:3]
colnames(RT2)[1]<-"RT_detected_2"
RT<-join(RT1,RT2)
fit <- coef(lm(RT$RT_detected ~ RT$RT_detected_2))
lib2a$RT_detected<-(lib2a$RT_detected*fit[2])+fit[1]
lib2a$iRT<-lib2a$RT_detected

lib2b<-lib2[lib2$RT_detected>=86,]
RT1<-lib1[,c(1,3,8)]
RT2<-lib2b[,c(1,3,8)]
RT1<-RT1[RT1$modification_sequence%in%RT2$modification_sequence,]
RT2<-RT2[RT2$modification_sequence%in%RT1$modification_sequence,]
RT1<-RT1[!duplicated(RT1$modification_sequence),2:3]
RT2<-RT2[!duplicated(RT2$modification_sequence),2:3]
colnames(RT2)[1]<-"RT_detected_2"
RT<-join(RT1,RT2)
fit <- coef(lm(RT$RT_detected ~ RT$RT_detected_2))
lib2b$RT_detected<-(lib2b$RT_detected*fit[2])+fit[1]
lib2b$iRT<-lib2b$RT_detected

lib2<-rbind(lib2a,lib2b)

buildSpectraLibPair(lib1, lib2, outputFile = "ExtendedLib_Colon.txt", plot = TRUE,
                    clean = FALSE, merge = TRUE, parseAcc = TRUE, consolidateAccession = TRUE,
                    nomod=TRUE, conf.cutoff=0.9)

#3

conf2<-read.xlsx("170324_09_PP_GPlibrary_Colon1__FDR.xlsx", sheet = "Search Summary",
                 colNames = FALSE, rows=33)[1,4]
lib1<-readLibFile("ExtendedLib_Colon.txt",clean=FALSE)
lib2<-readLibFile("170324_09_PP_GPlibrary_Colon1.txt",clean = TRUE,nomod=TRUE,nomc=FALSE, conf.cutoff=conf2)
lib2<-canonicalFormat(lib2, format = "peakview")
lib2<-lib2[lib2$uniprot_id!="[ RT-Cal protein ]",]
lib2<-lib2[lib2$uniprot_id!="1/[ RT-Cal protein ]",]
lib1<-lib1[!grepl("NA",rownames(lib1)),]
lib2<-lib2[!grepl("NA",rownames(lib2)),]
lib1$iRT<-lib1$RT_detected

# checkQuality(lib1,lib2)

lib2a<-lib2[lib2$RT_detected<87,]
RT1<-lib1[,c(1,3,8)]
RT2<-lib2a[,c(1,3,8)]
RT1<-RT1[RT1$modification_sequence%in%RT2$modification_sequence,]
RT2<-RT2[RT2$modification_sequence%in%RT1$modification_sequence,]
RT1<-RT1[!duplicated(RT1$modification_sequence),2:3]
RT2<-RT2[!duplicated(RT2$modification_sequence),2:3]
colnames(RT2)[1]<-"RT_detected_2"
RT<-join(RT1,RT2)
fit <- coef(lm(RT$RT_detected ~ RT$RT_detected_2))
lib2a$RT_detected<-(lib2a$RT_detected*fit[2])+fit[1]
lib2a$iRT<-lib2a$RT_detected

lib2b<-lib2[lib2$RT_detected>=86,]
RT1<-lib1[,c(1,3,8)]
RT2<-lib2b[,c(1,3,8)]
RT1<-RT1[RT1$modification_sequence%in%RT2$modification_sequence,]
RT2<-RT2[RT2$modification_sequence%in%RT1$modification_sequence,]
RT1<-RT1[!duplicated(RT1$modification_sequence),2:3]
RT2<-RT2[!duplicated(RT2$modification_sequence),2:3]
colnames(RT2)[1]<-"RT_detected_2"
RT<-join(RT1,RT2)
fit <- coef(lm(RT$RT_detected ~ RT$RT_detected_2))
lib2b$RT_detected<-(lib2b$RT_detected*fit[2])+fit[1]
lib2b$iRT<-lib2b$RT_detected

lib2<-rbind(lib2a,lib2b)

buildSpectraLibPair(lib1, lib2, outputFile = "ExtendedLib_Colon.txt", plot = TRUE,
                    clean = FALSE, merge = TRUE, parseAcc = TRUE, consolidateAccession = TRUE,
                    nomod=TRUE, conf.cutoff=0.9)

x<-read.delim("ExtendedLib_Colon.txt")
y<-as.data.frame(table(x$stripped_sequence))
y<-y[y$Freq>=5,]
x<-x[x$stripped_sequence%in%y$Var1,]
write.table(x,"ExtendedLib_Colon.txt",sep="\t")

# Cerebrum

#1

conf1<-read.xlsx("170303_05_PP_GP_Cerebrum__FDR.xlsx", sheet = "Search Summary",
                 colNames = FALSE, rows=33)[1,4]
conf2<-read.xlsx("170310_05_PP_GP_Cerebrum__FDR.xlsx", sheet = "Search Summary",
                 colNames = FALSE, rows=33)[1,4]
lib1<-readLibFile("170303_05_PP_GP_Cerebrum.txt",clean = TRUE,nomod=TRUE,nomc=FALSE, conf.cutoff=conf1)
lib2<-readLibFile("170310_05_PP_GP_Cerebrum.txt",clean = TRUE,nomod=TRUE,nomc=FALSE, conf.cutoff=conf2)
lib1<-canonicalFormat(lib1, format = "peakview")
lib2<-canonicalFormat(lib2, format = "peakview")
lib1<-lib1[lib1$uniprot_id!="[ RT-Cal protein ]",]
lib1<-lib1[lib1$uniprot_id!="1/[ RT-Cal protein ]",]
lib2<-lib2[lib2$uniprot_id!="[ RT-Cal protein ]",]
lib2<-lib2[lib2$uniprot_id!="1/[ RT-Cal protein ]",]
lib1<-lib1[!grepl("NA",rownames(lib1)),]
lib2<-lib2[!grepl("NA",rownames(lib2)),]

# checkQuality(lib1,lib2)

lib2a<-lib2[lib2$RT_detected<60,]
RT1<-lib1[,c(1,3,8)]
RT2<-lib2a[,c(1,3,8)]
RT1<-RT1[RT1$modification_sequence%in%RT2$modification_sequence,]
RT2<-RT2[RT2$modification_sequence%in%RT1$modification_sequence,]
RT1<-RT1[!duplicated(RT1$modification_sequence),2:3]
RT2<-RT2[!duplicated(RT2$modification_sequence),2:3]
colnames(RT2)[1]<-"RT_detected_2"
RT<-join(RT1,RT2)
fit <- coef(lm(RT$RT_detected ~ RT$RT_detected_2))
lib2a$RT_detected<-(lib2a$RT_detected*fit[2])+fit[1]
lib2a$iRT<-lib2a$RT_detected

lib2b<-lib2[lib2$RT_detected>=60,]
lib2b<-lib2b[lib2b$RT_detected<96,]
RT1<-lib1[,c(1,3,8)]
RT2<-lib2b[,c(1,3,8)]
RT1<-RT1[RT1$modification_sequence%in%RT2$modification_sequence,]
RT2<-RT2[RT2$modification_sequence%in%RT1$modification_sequence,]
RT1<-RT1[!duplicated(RT1$modification_sequence),2:3]
RT2<-RT2[!duplicated(RT2$modification_sequence),2:3]
colnames(RT2)[1]<-"RT_detected_2"
RT<-join(RT1,RT2)
fit <- coef(lm(RT$RT_detected ~ RT$RT_detected_2))
lib2b$RT_detected<-(lib2b$RT_detected*fit[2])+fit[1]
lib2b$iRT<-lib2b$RT_detected

lib2c<-lib2[lib2$RT_detected>=95,]
RT1<-lib1[,c(1,3,8)]
RT2<-lib2c[,c(1,3,8)]
RT1<-RT1[RT1$modification_sequence%in%RT2$modification_sequence,]
RT2<-RT2[RT2$modification_sequence%in%RT1$modification_sequence,]
RT1<-RT1[!duplicated(RT1$modification_sequence),2:3]
RT2<-RT2[!duplicated(RT2$modification_sequence),2:3]
colnames(RT2)[1]<-"RT_detected_2"
RT<-join(RT1,RT2)
fit <- coef(lm(RT$RT_detected ~ RT$RT_detected_2))
lib2c$RT_detected<-(lib2c$RT_detected*fit[2])+fit[1]
lib2c$iRT<-lib2c$RT_detected

lib2<-rbind(lib2a,lib2b,lib2c)

buildSpectraLibPair(lib1, lib2, outputFile = "ExtendedLib_Cerebrum.txt", plot = TRUE,
                    clean = FALSE, merge = TRUE, parseAcc = TRUE, consolidateAccession = TRUE,
                    nomod=TRUE, conf.cutoff=0.9)
#2

conf2<-read.xlsx("170324_10_PP_GPlibrary_Cerebrum__FDR.xlsx", sheet = "Search Summary",
                 colNames = FALSE, rows=33)[1,4]
lib1<-readLibFile("ExtendedLib_Cerebrum.txt",clean=FALSE)
lib2<-readLibFile("170324_10_PP_GPlibrary_Cerebrum.txt",clean =TRUE,nomod=TRUE,nomc=FALSE,conf.cutoff=conf2)
lib2<-canonicalFormat(lib2, format = "peakview")
lib2<-lib2[lib2$uniprot_id!="[ RT-Cal protein ]",]
lib2<-lib2[lib2$uniprot_id!="1/[ RT-Cal protein ]",]
lib1<-lib1[!grepl("NA",rownames(lib1)),]
lib2<-lib2[!grepl("NA",rownames(lib2)),]
lib1$iRT<-lib1$RT_detected

# checkQuality(lib1,lib2)

lib2a<-lib2[lib2$RT_detected<87,]
RT1<-lib1[,c(1,3,8)]
RT2<-lib2a[,c(1,3,8)]
RT1<-RT1[RT1$modification_sequence%in%RT2$modification_sequence,]
RT2<-RT2[RT2$modification_sequence%in%RT1$modification_sequence,]
RT1<-RT1[!duplicated(RT1$modification_sequence),2:3]
RT2<-RT2[!duplicated(RT2$modification_sequence),2:3]
colnames(RT2)[1]<-"RT_detected_2"
RT<-join(RT1,RT2)
fit <- coef(lm(RT$RT_detected ~ RT$RT_detected_2))
lib2a$RT_detected<-(lib2a$RT_detected*fit[2])+fit[1]
lib2a$iRT<-lib2a$RT_detected

lib2b<-lib2[lib2$RT_detected>=86,]
RT1<-lib1[,c(1,3,8)]
RT2<-lib2b[,c(1,3,8)]
RT1<-RT1[RT1$modification_sequence%in%RT2$modification_sequence,]
RT2<-RT2[RT2$modification_sequence%in%RT1$modification_sequence,]
RT1<-RT1[!duplicated(RT1$modification_sequence),2:3]
RT2<-RT2[!duplicated(RT2$modification_sequence),2:3]
colnames(RT2)[1]<-"RT_detected_2"
RT<-join(RT1,RT2)
fit <- coef(lm(RT$RT_detected ~ RT$RT_detected_2))
lib2b$RT_detected<-(lib2b$RT_detected*fit[2])+fit[1]
lib2b$iRT<-lib2b$RT_detected

lib2<-rbind(lib2a,lib2b)

buildSpectraLibPair(lib1, lib2, outputFile = "ExtendedLib_Cerebrum.txt", plot = TRUE,
                    clean = FALSE, merge = TRUE, parseAcc = TRUE, consolidateAccession = TRUE,
                    nomod=TRUE, conf.cutoff=0.9)

#3

conf2<-read.xlsx("170324_10_PP_GPlibrary_Cerebrum1__FDR.xlsx", sheet = "Search Summary",
                 colNames = FALSE, rows=33)[1,4]
lib1<-readLibFile("ExtendedLib_Cerebrum.txt",clean=FALSE)
lib2<-readLibFile("170324_10_PP_GPlibrary_Cerebrum1.txt",clean=TRUE,nomod=TRUE,nomc=FALSE,conf.cutoff=conf2)
lib2<-canonicalFormat(lib2, format = "peakview")
lib2<-lib2[lib2$uniprot_id!="[ RT-Cal protein ]",]
lib2<-lib2[lib2$uniprot_id!="1/[ RT-Cal protein ]",]
lib1<-lib1[!grepl("NA",rownames(lib1)),]
lib2<-lib2[!grepl("NA",rownames(lib2)),]
lib1$iRT<-lib1$RT_detected

# checkQuality(lib1,lib2)

lib2a<-lib2[lib2$RT_detected<87,]
RT1<-lib1[,c(1,3,8)]
RT2<-lib2a[,c(1,3,8)]
RT1<-RT1[RT1$modification_sequence%in%RT2$modification_sequence,]
RT2<-RT2[RT2$modification_sequence%in%RT1$modification_sequence,]
RT1<-RT1[!duplicated(RT1$modification_sequence),2:3]
RT2<-RT2[!duplicated(RT2$modification_sequence),2:3]
colnames(RT2)[1]<-"RT_detected_2"
RT<-join(RT1,RT2)
fit <- coef(lm(RT$RT_detected ~ RT$RT_detected_2))
lib2a$RT_detected<-(lib2a$RT_detected*fit[2])+fit[1]
lib2a$iRT<-lib2a$RT_detected

lib2b<-lib2[lib2$RT_detected>=86,]
RT1<-lib1[,c(1,3,8)]
RT2<-lib2b[,c(1,3,8)]
RT1<-RT1[RT1$modification_sequence%in%RT2$modification_sequence,]
RT2<-RT2[RT2$modification_sequence%in%RT1$modification_sequence,]
RT1<-RT1[!duplicated(RT1$modification_sequence),2:3]
RT2<-RT2[!duplicated(RT2$modification_sequence),2:3]
colnames(RT2)[1]<-"RT_detected_2"
RT<-join(RT1,RT2)
fit <- coef(lm(RT$RT_detected ~ RT$RT_detected_2))
lib2b$RT_detected<-(lib2b$RT_detected*fit[2])+fit[1]
lib2b$iRT<-lib2b$RT_detected

lib2<-rbind(lib2a,lib2b)

buildSpectraLibPair(lib1, lib2, outputFile = "ExtendedLib_Cerebrum.txt", plot = TRUE,
                    clean = FALSE, merge = TRUE, parseAcc = TRUE, consolidateAccession = TRUE,
                    nomod=TRUE, conf.cutoff=0.9)

x<-read.delim("ExtendedLib_Cerebrum.txt")
y<-as.data.frame(table(x$stripped_sequence))
y<-y[y$Freq>=5,]
x<-x[x$stripped_sequence%in%y$Var1,]
write.table(x,"ExtendedLib_Cerebrum.txt",sep="\t")

# Stomach

#1

conf1<-read.xlsx("170303_06_PP_GP_Stomach__FDR.xlsx", sheet = "Search Summary",
                 colNames = FALSE, rows=33)[1,4]
conf2<-read.xlsx("170310_06_PP_GP_Stomach__FDR.xlsx", sheet = "Search Summary",
                 colNames = FALSE, rows=33)[1,4]
lib1<-readLibFile("170303_06_PP_GP_Stomach.txt",clean = TRUE,nomod=TRUE,nomc=FALSE, conf.cutoff=conf1)
lib2<-readLibFile("170310_06_PP_GP_Stomach.txt",clean = TRUE,nomod=TRUE,nomc=FALSE, conf.cutoff=conf2)
lib1<-canonicalFormat(lib1, format = "peakview")
lib2<-canonicalFormat(lib2, format = "peakview")
lib1<-lib1[lib1$uniprot_id!="[ RT-Cal protein ]",]
lib1<-lib1[lib1$uniprot_id!="1/[ RT-Cal protein ]",]
lib2<-lib2[lib2$uniprot_id!="[ RT-Cal protein ]",]
lib2<-lib2[lib2$uniprot_id!="1/[ RT-Cal protein ]",]
lib1<-lib1[!grepl("NA",rownames(lib1)),]
lib2<-lib2[!grepl("NA",rownames(lib2)),]

# checkQuality(lib1,lib2)

lib2a<-lib2[lib2$RT_detected<60,]
RT1<-lib1[,c(1,3,8)]
RT2<-lib2a[,c(1,3,8)]
RT1<-RT1[RT1$modification_sequence%in%RT2$modification_sequence,]
RT2<-RT2[RT2$modification_sequence%in%RT1$modification_sequence,]
RT1<-RT1[!duplicated(RT1$modification_sequence),2:3]
RT2<-RT2[!duplicated(RT2$modification_sequence),2:3]
colnames(RT2)[1]<-"RT_detected_2"
RT<-join(RT1,RT2)
fit <- coef(lm(RT$RT_detected ~ RT$RT_detected_2))
lib2a$RT_detected<-(lib2a$RT_detected*fit[2])+fit[1]
lib2a$iRT<-lib2a$RT_detected

lib2b<-lib2[lib2$RT_detected>=60,]
lib2b<-lib2b[lib2b$RT_detected<96,]
RT1<-lib1[,c(1,3,8)]
RT2<-lib2b[,c(1,3,8)]
RT1<-RT1[RT1$modification_sequence%in%RT2$modification_sequence,]
RT2<-RT2[RT2$modification_sequence%in%RT1$modification_sequence,]
RT1<-RT1[!duplicated(RT1$modification_sequence),2:3]
RT2<-RT2[!duplicated(RT2$modification_sequence),2:3]
colnames(RT2)[1]<-"RT_detected_2"
RT<-join(RT1,RT2)
fit <- coef(lm(RT$RT_detected ~ RT$RT_detected_2))
lib2b$RT_detected<-(lib2b$RT_detected*fit[2])+fit[1]
lib2b$iRT<-lib2b$RT_detected

lib2c<-lib2[lib2$RT_detected>=95,]
RT1<-lib1[,c(1,3,8)]
RT2<-lib2c[,c(1,3,8)]
RT1<-RT1[RT1$modification_sequence%in%RT2$modification_sequence,]
RT2<-RT2[RT2$modification_sequence%in%RT1$modification_sequence,]
RT1<-RT1[!duplicated(RT1$modification_sequence),2:3]
RT2<-RT2[!duplicated(RT2$modification_sequence),2:3]
colnames(RT2)[1]<-"RT_detected_2"
RT<-join(RT1,RT2)
fit <- coef(lm(RT$RT_detected ~ RT$RT_detected_2))
lib2c$RT_detected<-(lib2c$RT_detected*fit[2])+fit[1]
lib2c$iRT<-lib2c$RT_detected

lib2<-rbind(lib2a,lib2b,lib2c)

buildSpectraLibPair(lib1, lib2, outputFile = "ExtendedLib_Stomach.txt", plot = TRUE,
                    clean = FALSE, merge = TRUE, parseAcc = TRUE, consolidateAccession = TRUE,
                    nomod=TRUE, conf.cutoff=0.9)
#2

conf2<-read.xlsx("170324_11_PP_GPlibrary_Stomach__FDR.xlsx", sheet = "Search Summary",
                 colNames = FALSE, rows=33)[1,4]
lib1<-readLibFile("ExtendedLib_Stomach.txt",clean=FALSE)
lib2<-readLibFile("170324_11_PP_GPlibrary_Stomach.txt",clean = TRUE,nomod=TRUE,nomc=FALSE,conf.cutoff=conf2)
lib2<-canonicalFormat(lib2, format = "peakview")
lib2<-lib2[lib2$uniprot_id!="[ RT-Cal protein ]",]
lib2<-lib2[lib2$uniprot_id!="1/[ RT-Cal protein ]",]
lib1<-lib1[!grepl("NA",rownames(lib1)),]
lib2<-lib2[!grepl("NA",rownames(lib2)),]
lib1$iRT<-lib1$RT_detected

# checkQuality(lib1,lib2)

lib2a<-lib2[lib2$RT_detected<87,]
RT1<-lib1[,c(1,3,8)]
RT2<-lib2a[,c(1,3,8)]
RT1<-RT1[RT1$modification_sequence%in%RT2$modification_sequence,]
RT2<-RT2[RT2$modification_sequence%in%RT1$modification_sequence,]
RT1<-RT1[!duplicated(RT1$modification_sequence),2:3]
RT2<-RT2[!duplicated(RT2$modification_sequence),2:3]
colnames(RT2)[1]<-"RT_detected_2"
RT<-join(RT1,RT2)
fit <- coef(lm(RT$RT_detected ~ RT$RT_detected_2))
lib2a$RT_detected<-(lib2a$RT_detected*fit[2])+fit[1]
lib2a$iRT<-lib2a$RT_detected

lib2b<-lib2[lib2$RT_detected>=86,]
RT1<-lib1[,c(1,3,8)]
RT2<-lib2b[,c(1,3,8)]
RT1<-RT1[RT1$modification_sequence%in%RT2$modification_sequence,]
RT2<-RT2[RT2$modification_sequence%in%RT1$modification_sequence,]
RT1<-RT1[!duplicated(RT1$modification_sequence),2:3]
RT2<-RT2[!duplicated(RT2$modification_sequence),2:3]
colnames(RT2)[1]<-"RT_detected_2"
RT<-join(RT1,RT2)
fit <- coef(lm(RT$RT_detected ~ RT$RT_detected_2))
lib2b$RT_detected<-(lib2b$RT_detected*fit[2])+fit[1]
lib2b$iRT<-lib2b$RT_detected

lib2<-rbind(lib2a,lib2b)

buildSpectraLibPair(lib1, lib2, outputFile = "ExtendedLib_Stomach.txt", plot = TRUE,
                    clean = FALSE, merge = TRUE, parseAcc = TRUE, consolidateAccession = TRUE,
                    nomod=TRUE, conf.cutoff=0.9)

#3

conf2<-read.xlsx("170324_11_PP_GPlibrary_Stomach1__FDR.xlsx", sheet = "Search Summary",
                 colNames = FALSE, rows=33)[1,4]
lib1<-readLibFile("ExtendedLib_Stomach.txt",clean=FALSE)
lib2<-readLibFile("170324_11_PP_GPlibrary_Stomach1.txt",clean = TRUE,nomod=TRUE,nomc=FALSE,
                  conf.cutoff=conf2)
lib2<-canonicalFormat(lib2, format = "peakview")
lib2<-lib2[lib2$uniprot_id!="[ RT-Cal protein ]",]
lib2<-lib2[lib2$uniprot_id!="1/[ RT-Cal protein ]",]
lib1<-lib1[!grepl("NA",rownames(lib1)),]
lib2<-lib2[!grepl("NA",rownames(lib2)),]
lib1$iRT<-lib1$RT_detected

# checkQuality(lib1,lib2)

lib2a<-lib2[lib2$RT_detected<87,]
RT1<-lib1[,c(1,3,8)]
RT2<-lib2a[,c(1,3,8)]
RT1<-RT1[RT1$modification_sequence%in%RT2$modification_sequence,]
RT2<-RT2[RT2$modification_sequence%in%RT1$modification_sequence,]
RT1<-RT1[!duplicated(RT1$modification_sequence),2:3]
RT2<-RT2[!duplicated(RT2$modification_sequence),2:3]
colnames(RT2)[1]<-"RT_detected_2"
RT<-join(RT1,RT2)
fit <- coef(lm(RT$RT_detected ~ RT$RT_detected_2))
lib2a$RT_detected<-(lib2a$RT_detected*fit[2])+fit[1]
lib2a$iRT<-lib2a$RT_detected

lib2b<-lib2[lib2$RT_detected>=86,]
RT1<-lib1[,c(1,3,8)]
RT2<-lib2b[,c(1,3,8)]
RT1<-RT1[RT1$modification_sequence%in%RT2$modification_sequence,]
RT2<-RT2[RT2$modification_sequence%in%RT1$modification_sequence,]
RT1<-RT1[!duplicated(RT1$modification_sequence),2:3]
RT2<-RT2[!duplicated(RT2$modification_sequence),2:3]
colnames(RT2)[1]<-"RT_detected_2"
RT<-join(RT1,RT2)
fit <- coef(lm(RT$RT_detected ~ RT$RT_detected_2))
lib2b$RT_detected<-(lib2b$RT_detected*fit[2])+fit[1]
lib2b$iRT<-lib2b$RT_detected

lib2<-rbind(lib2a,lib2b)

buildSpectraLibPair(lib1, lib2, outputFile = "ExtendedLib_Stomach.txt", plot = TRUE,
                    clean = FALSE, merge = TRUE, parseAcc = TRUE, consolidateAccession = TRUE,
                    nomod=TRUE, conf.cutoff=0.9)

x<-read.delim("ExtendedLib_Stomach.txt")
y<-as.data.frame(table(x$stripped_sequence))
y<-y[y$Freq>=5,]
x<-x[x$stripped_sequence%in%y$Var1,]
write.table(x,"ExtendedLib_Stomach.txt",sep="\t")

# Placenta

#1

conf1<-read.xlsx("170303_07_PP_GP_Placenta__FDR.xlsx", sheet = "Search Summary",
                 colNames = FALSE, rows=33)[1,4]
conf2<-read.xlsx("170310_07_PP_GP_Placenta__FDR.xlsx", sheet = "Search Summary",
                 colNames = FALSE, rows=33)[1,4]
lib1<-readLibFile("170303_07_PP_GP_Placenta.txt",clean = TRUE,nomod=TRUE,nomc=FALSE, conf.cutoff=conf1)
lib2<-readLibFile("170310_07_PP_GP_Placenta.txt",clean = TRUE,nomod=TRUE,nomc=FALSE, conf.cutoff=conf2)
lib1<-canonicalFormat(lib1, format = "peakview")
lib2<-canonicalFormat(lib2, format = "peakview")
lib1<-lib1[lib1$uniprot_id!="[ RT-Cal protein ]",]
lib1<-lib1[lib1$uniprot_id!="1/[ RT-Cal protein ]",]
lib2<-lib2[lib2$uniprot_id!="[ RT-Cal protein ]",]
lib2<-lib2[lib2$uniprot_id!="1/[ RT-Cal protein ]",]
lib1<-lib1[!grepl("NA",rownames(lib1)),]
lib2<-lib2[!grepl("NA",rownames(lib2)),]

# checkQuality(lib1,lib2)

lib2a<-lib2[lib2$RT_detected<60,]
RT1<-lib1[,c(1,3,8)]
RT2<-lib2a[,c(1,3,8)]
RT1<-RT1[RT1$modification_sequence%in%RT2$modification_sequence,]
RT2<-RT2[RT2$modification_sequence%in%RT1$modification_sequence,]
RT1<-RT1[!duplicated(RT1$modification_sequence),2:3]
RT2<-RT2[!duplicated(RT2$modification_sequence),2:3]
colnames(RT2)[1]<-"RT_detected_2"
RT<-join(RT1,RT2)
fit <- coef(lm(RT$RT_detected ~ RT$RT_detected_2))
lib2a$RT_detected<-(lib2a$RT_detected*fit[2])+fit[1]
lib2a$iRT<-lib2a$RT_detected

lib2b<-lib2[lib2$RT_detected>=60,]
lib2b<-lib2b[lib2b$RT_detected<96,]
RT1<-lib1[,c(1,3,8)]
RT2<-lib2b[,c(1,3,8)]
RT1<-RT1[RT1$modification_sequence%in%RT2$modification_sequence,]
RT2<-RT2[RT2$modification_sequence%in%RT1$modification_sequence,]
RT1<-RT1[!duplicated(RT1$modification_sequence),2:3]
RT2<-RT2[!duplicated(RT2$modification_sequence),2:3]
colnames(RT2)[1]<-"RT_detected_2"
RT<-join(RT1,RT2)
fit <- coef(lm(RT$RT_detected ~ RT$RT_detected_2))
lib2b$RT_detected<-(lib2b$RT_detected*fit[2])+fit[1]
lib2b$iRT<-lib2b$RT_detected

lib2c<-lib2[lib2$RT_detected>=95,]
RT1<-lib1[,c(1,3,8)]
RT2<-lib2c[,c(1,3,8)]
RT1<-RT1[RT1$modification_sequence%in%RT2$modification_sequence,]
RT2<-RT2[RT2$modification_sequence%in%RT1$modification_sequence,]
RT1<-RT1[!duplicated(RT1$modification_sequence),2:3]
RT2<-RT2[!duplicated(RT2$modification_sequence),2:3]
colnames(RT2)[1]<-"RT_detected_2"
RT<-join(RT1,RT2)
fit <- coef(lm(RT$RT_detected ~ RT$RT_detected_2))
lib2c$RT_detected<-(lib2c$RT_detected*fit[2])+fit[1]
lib2c$iRT<-lib2c$RT_detected

lib2<-rbind(lib2a,lib2b,lib2c)

buildSpectraLibPair(lib1, lib2, outputFile = "ExtendedLib_Placenta.txt", plot = TRUE,
                    clean = FALSE, merge = TRUE, parseAcc = TRUE, consolidateAccession = TRUE,
                    nomod=TRUE, conf.cutoff=0.9)
#2

conf2<-read.xlsx("170324_12_PP_GPlibrary_Placenta__FDR.xlsx", sheet = "Search Summary",
                 colNames = FALSE, rows=33)[1,4]
lib1<-readLibFile("ExtendedLib_Placenta.txt",clean=FALSE)
lib2<-readLibFile("170324_12_PP_GPlibrary_Placenta.txt",clean =TRUE,nomod=TRUE,nomc=FALSE,conf.cutoff=conf2)
lib2<-canonicalFormat(lib2, format = "peakview")
lib2<-lib2[lib2$uniprot_id!="[ RT-Cal protein ]",]
lib2<-lib2[lib2$uniprot_id!="1/[ RT-Cal protein ]",]
lib1<-lib1[!grepl("NA",rownames(lib1)),]
lib2<-lib2[!grepl("NA",rownames(lib2)),]
lib1$iRT<-lib1$RT_detected

# checkQuality(lib1,lib2)

lib2a<-lib2[lib2$RT_detected<87,]
RT1<-lib1[,c(1,3,8)]
RT2<-lib2a[,c(1,3,8)]
RT1<-RT1[RT1$modification_sequence%in%RT2$modification_sequence,]
RT2<-RT2[RT2$modification_sequence%in%RT1$modification_sequence,]
RT1<-RT1[!duplicated(RT1$modification_sequence),2:3]
RT2<-RT2[!duplicated(RT2$modification_sequence),2:3]
colnames(RT2)[1]<-"RT_detected_2"
RT<-join(RT1,RT2)
fit <- coef(lm(RT$RT_detected ~ RT$RT_detected_2))
lib2a$RT_detected<-(lib2a$RT_detected*fit[2])+fit[1]
lib2a$iRT<-lib2a$RT_detected

lib2b<-lib2[lib2$RT_detected>=86,]
RT1<-lib1[,c(1,3,8)]
RT2<-lib2b[,c(1,3,8)]
RT1<-RT1[RT1$modification_sequence%in%RT2$modification_sequence,]
RT2<-RT2[RT2$modification_sequence%in%RT1$modification_sequence,]
RT1<-RT1[!duplicated(RT1$modification_sequence),2:3]
RT2<-RT2[!duplicated(RT2$modification_sequence),2:3]
colnames(RT2)[1]<-"RT_detected_2"
RT<-join(RT1,RT2)
fit <- coef(lm(RT$RT_detected ~ RT$RT_detected_2))
lib2b$RT_detected<-(lib2b$RT_detected*fit[2])+fit[1]
lib2b$iRT<-lib2b$RT_detected

lib2<-rbind(lib2a,lib2b)

buildSpectraLibPair(lib1, lib2, outputFile = "ExtendedLib_Placenta.txt", plot = TRUE,
                    clean = FALSE, merge = TRUE, parseAcc = TRUE, consolidateAccession = TRUE,
                    nomod=TRUE, conf.cutoff=0.9)

#3

conf2<-read.xlsx("170324_12_PP_GPlibrary_Placenta1__FDR.xlsx", sheet = "Search Summary",
                 colNames = FALSE, rows=33)[1,4]
lib1<-readLibFile("ExtendedLib_Placenta.txt",clean=FALSE)
lib2<-readLibFile("170324_12_PP_GPlibrary_Placenta1.txt",clean = TRUE,nomod=TRUE,nomc=FALSE,
                  conf.cutoff=conf2)
lib2<-canonicalFormat(lib2, format = "peakview")
lib2<-lib2[lib2$uniprot_id!="[ RT-Cal protein ]",]
lib2<-lib2[lib2$uniprot_id!="1/[ RT-Cal protein ]",]
lib1<-lib1[!grepl("NA",rownames(lib1)),]
lib2<-lib2[!grepl("NA",rownames(lib2)),]
lib1$iRT<-lib1$RT_detected

# checkQuality(lib1,lib2)

lib2a<-lib2[lib2$RT_detected<87,]
RT1<-lib1[,c(1,3,8)]
RT2<-lib2a[,c(1,3,8)]
RT1<-RT1[RT1$modification_sequence%in%RT2$modification_sequence,]
RT2<-RT2[RT2$modification_sequence%in%RT1$modification_sequence,]
RT1<-RT1[!duplicated(RT1$modification_sequence),2:3]
RT2<-RT2[!duplicated(RT2$modification_sequence),2:3]
colnames(RT2)[1]<-"RT_detected_2"
RT<-join(RT1,RT2)
fit <- coef(lm(RT$RT_detected ~ RT$RT_detected_2))
lib2a$RT_detected<-(lib2a$RT_detected*fit[2])+fit[1]
lib2a$iRT<-lib2a$RT_detected

lib2b<-lib2[lib2$RT_detected>=86,]
RT1<-lib1[,c(1,3,8)]
RT2<-lib2b[,c(1,3,8)]
RT1<-RT1[RT1$modification_sequence%in%RT2$modification_sequence,]
RT2<-RT2[RT2$modification_sequence%in%RT1$modification_sequence,]
RT1<-RT1[!duplicated(RT1$modification_sequence),2:3]
RT2<-RT2[!duplicated(RT2$modification_sequence),2:3]
colnames(RT2)[1]<-"RT_detected_2"
RT<-join(RT1,RT2)
fit <- coef(lm(RT$RT_detected ~ RT$RT_detected_2))
lib2b$RT_detected<-(lib2b$RT_detected*fit[2])+fit[1]
lib2b$iRT<-lib2b$RT_detected

lib2<-rbind(lib2a,lib2b)

buildSpectraLibPair(lib1, lib2, outputFile = "ExtendedLib_Placenta.txt", plot = TRUE,
                    clean = FALSE, merge = TRUE, parseAcc = TRUE, consolidateAccession = TRUE,
                    nomod=TRUE, conf.cutoff=0.9)


x<-read.delim("ExtendedLib_Placenta.txt")
y<-as.data.frame(table(x$stripped_sequence))
y<-y[y$Freq>=5,]
x<-x[x$stripped_sequence%in%y$Var1,]
write.table(x,"ExtendedLib_Placenta.txt",sep="\t")

# LargeIntestine

#1

conf1<-read.xlsx("170303_08_PP_GP_LargeIntestine__FDR.xlsx", sheet = "Search Summary",
                 colNames = FALSE, rows=33)[1,4]
conf2<-read.xlsx("170310_08_PP_GP_LargeIntestine__FDR.xlsx", sheet = "Search Summary",
                 colNames = FALSE, rows=33)[1,4]
lib1<-readLibFile("170303_08_PP_GP_LargeIntestine.txt",clean = TRUE,nomod=TRUE,nomc=FALSE,conf.cutoff=conf1)
lib2<-readLibFile("170310_08_PP_GP_LargeIntestine.txt",clean = TRUE,nomod=TRUE,nomc=FALSE,conf.cutoff=conf2)
lib1<-canonicalFormat(lib1, format = "peakview")
lib2<-canonicalFormat(lib2, format = "peakview")
lib1<-lib1[lib1$uniprot_id!="[ RT-Cal protein ]",]
lib1<-lib1[lib1$uniprot_id!="1/[ RT-Cal protein ]",]
lib2<-lib2[lib2$uniprot_id!="[ RT-Cal protein ]",]
lib2<-lib2[lib2$uniprot_id!="1/[ RT-Cal protein ]",]
lib1<-lib1[!grepl("NA",rownames(lib1)),]
lib2<-lib2[!grepl("NA",rownames(lib2)),]

# checkQuality(lib1,lib2)

lib2a<-lib2[lib2$RT_detected<60,]
RT1<-lib1[,c(1,3,8)]
RT2<-lib2a[,c(1,3,8)]
RT1<-RT1[RT1$modification_sequence%in%RT2$modification_sequence,]
RT2<-RT2[RT2$modification_sequence%in%RT1$modification_sequence,]
RT1<-RT1[!duplicated(RT1$modification_sequence),2:3]
RT2<-RT2[!duplicated(RT2$modification_sequence),2:3]
colnames(RT2)[1]<-"RT_detected_2"
RT<-join(RT1,RT2)
fit <- coef(lm(RT$RT_detected ~ RT$RT_detected_2))
lib2a$RT_detected<-(lib2a$RT_detected*fit[2])+fit[1]
lib2a$iRT<-lib2a$RT_detected

lib2b<-lib2[lib2$RT_detected>=60,]
lib2b<-lib2b[lib2b$RT_detected<96,]
RT1<-lib1[,c(1,3,8)]
RT2<-lib2b[,c(1,3,8)]
RT1<-RT1[RT1$modification_sequence%in%RT2$modification_sequence,]
RT2<-RT2[RT2$modification_sequence%in%RT1$modification_sequence,]
RT1<-RT1[!duplicated(RT1$modification_sequence),2:3]
RT2<-RT2[!duplicated(RT2$modification_sequence),2:3]
colnames(RT2)[1]<-"RT_detected_2"
RT<-join(RT1,RT2)
fit <- coef(lm(RT$RT_detected ~ RT$RT_detected_2))
lib2b$RT_detected<-(lib2b$RT_detected*fit[2])+fit[1]
lib2b$iRT<-lib2b$RT_detected

lib2c<-lib2[lib2$RT_detected>=95,]
RT1<-lib1[,c(1,3,8)]
RT2<-lib2c[,c(1,3,8)]
RT1<-RT1[RT1$modification_sequence%in%RT2$modification_sequence,]
RT2<-RT2[RT2$modification_sequence%in%RT1$modification_sequence,]
RT1<-RT1[!duplicated(RT1$modification_sequence),2:3]
RT2<-RT2[!duplicated(RT2$modification_sequence),2:3]
colnames(RT2)[1]<-"RT_detected_2"
RT<-join(RT1,RT2)
fit <- coef(lm(RT$RT_detected ~ RT$RT_detected_2))
lib2c$RT_detected<-(lib2c$RT_detected*fit[2])+fit[1]
lib2c$iRT<-lib2c$RT_detected

lib2<-rbind(lib2a,lib2b,lib2c)

buildSpectraLibPair(lib1, lib2, outputFile = "ExtendedLib_LargeIntestine.txt", plot = TRUE,
                    clean = FALSE, merge = TRUE, parseAcc = TRUE, consolidateAccession = TRUE,
                    nomod=TRUE, conf.cutoff=0.9)
#2

conf2<-read.xlsx("170324_13_PP_GPlibrary_LargeIntestine__FDR.xlsx", sheet = "Search Summary",
                 colNames = FALSE, rows=33)[1,4]
lib1<-readLibFile("ExtendedLib_LargeIntestine.txt",clean=FALSE)
lib2<-readLibFile("170324_13_PP_GPlibrary_LargeIntestine.txt",clean = TRUE,nomod=TRUE,
                  nomc=FALSE, conf.cutoff=conf2)
lib2<-canonicalFormat(lib2, format = "peakview")
lib2<-lib2[lib2$uniprot_id!="[ RT-Cal protein ]",]
lib2<-lib2[lib2$uniprot_id!="1/[ RT-Cal protein ]",]
lib1<-lib1[!grepl("NA",rownames(lib1)),]
lib2<-lib2[!grepl("NA",rownames(lib2)),]
lib1$iRT<-lib1$RT_detected

# checkQuality(lib1,lib2)

lib2a<-lib2[lib2$RT_detected<87,]
RT1<-lib1[,c(1,3,8)]
RT2<-lib2a[,c(1,3,8)]
RT1<-RT1[RT1$modification_sequence%in%RT2$modification_sequence,]
RT2<-RT2[RT2$modification_sequence%in%RT1$modification_sequence,]
RT1<-RT1[!duplicated(RT1$modification_sequence),2:3]
RT2<-RT2[!duplicated(RT2$modification_sequence),2:3]
colnames(RT2)[1]<-"RT_detected_2"
RT<-join(RT1,RT2)
fit <- coef(lm(RT$RT_detected ~ RT$RT_detected_2))
lib2a$RT_detected<-(lib2a$RT_detected*fit[2])+fit[1]
lib2a$iRT<-lib2a$RT_detected

lib2b<-lib2[lib2$RT_detected>=86,]
RT1<-lib1[,c(1,3,8)]
RT2<-lib2b[,c(1,3,8)]
RT1<-RT1[RT1$modification_sequence%in%RT2$modification_sequence,]
RT2<-RT2[RT2$modification_sequence%in%RT1$modification_sequence,]
RT1<-RT1[!duplicated(RT1$modification_sequence),2:3]
RT2<-RT2[!duplicated(RT2$modification_sequence),2:3]
colnames(RT2)[1]<-"RT_detected_2"
RT<-join(RT1,RT2)
fit <- coef(lm(RT$RT_detected ~ RT$RT_detected_2))
lib2b$RT_detected<-(lib2b$RT_detected*fit[2])+fit[1]
lib2b$iRT<-lib2b$RT_detected

lib2<-rbind(lib2a,lib2b)

buildSpectraLibPair(lib1, lib2, outputFile = "ExtendedLib_LargeIntestine.txt", plot = TRUE,
                    clean = FALSE, merge = TRUE, parseAcc = TRUE, consolidateAccession = TRUE,
                    nomod=TRUE, conf.cutoff=0.9)


x<-read.delim("ExtendedLib_LargeIntestine.txt")
y<-as.data.frame(table(x$stripped_sequence))
y<-y[y$Freq>=5,]
x<-x[x$stripped_sequence%in%y$Var1,]
write.table(x,"ExtendedLib_LargeIntestine.txt",sep="\t")

# Kidney

#1

conf1<-read.xlsx("170303_09_PP_GP_Kidney__FDR.xlsx", sheet = "Search Summary",
                 colNames = FALSE, rows=33)[1,4]
conf2<-read.xlsx("170310_09_PP_GP_Kidney__FDR.xlsx", sheet = "Search Summary",
                 colNames = FALSE, rows=33)[1,4]
lib1<-readLibFile("170303_09_PP_GP_Kidney.txt",clean = TRUE,nomod=TRUE,nomc=FALSE, conf.cutoff=conf1)
lib2<-readLibFile("170310_09_PP_GP_Kidney.txt",clean = TRUE,nomod=TRUE,nomc=FALSE, conf.cutoff=conf2)
lib1<-canonicalFormat(lib1, format = "peakview")
lib2<-canonicalFormat(lib2, format = "peakview")
lib1<-lib1[lib1$uniprot_id!="[ RT-Cal protein ]",]
lib1<-lib1[lib1$uniprot_id!="1/[ RT-Cal protein ]",]
lib2<-lib2[lib2$uniprot_id!="[ RT-Cal protein ]",]
lib2<-lib2[lib2$uniprot_id!="1/[ RT-Cal protein ]",]
lib1<-lib1[!grepl("NA",rownames(lib1)),]
lib2<-lib2[!grepl("NA",rownames(lib2)),]

# checkQuality(lib1,lib2)

lib2a<-lib2[lib2$RT_detected<60,]
RT1<-lib1[,c(1,3,8)]
RT2<-lib2a[,c(1,3,8)]
RT1<-RT1[RT1$modification_sequence%in%RT2$modification_sequence,]
RT2<-RT2[RT2$modification_sequence%in%RT1$modification_sequence,]
RT1<-RT1[!duplicated(RT1$modification_sequence),2:3]
RT2<-RT2[!duplicated(RT2$modification_sequence),2:3]
colnames(RT2)[1]<-"RT_detected_2"
RT<-join(RT1,RT2)
fit <- coef(lm(RT$RT_detected ~ RT$RT_detected_2))
lib2a$RT_detected<-(lib2a$RT_detected*fit[2])+fit[1]
lib2a$iRT<-lib2a$RT_detected

lib2b<-lib2[lib2$RT_detected>=60,]
lib2b<-lib2b[lib2b$RT_detected<96,]
RT1<-lib1[,c(1,3,8)]
RT2<-lib2b[,c(1,3,8)]
RT1<-RT1[RT1$modification_sequence%in%RT2$modification_sequence,]
RT2<-RT2[RT2$modification_sequence%in%RT1$modification_sequence,]
RT1<-RT1[!duplicated(RT1$modification_sequence),2:3]
RT2<-RT2[!duplicated(RT2$modification_sequence),2:3]
colnames(RT2)[1]<-"RT_detected_2"
RT<-join(RT1,RT2)
fit <- coef(lm(RT$RT_detected ~ RT$RT_detected_2))
lib2b$RT_detected<-(lib2b$RT_detected*fit[2])+fit[1]
lib2b$iRT<-lib2b$RT_detected

lib2c<-lib2[lib2$RT_detected>=95,]
RT1<-lib1[,c(1,3,8)]
RT2<-lib2c[,c(1,3,8)]
RT1<-RT1[RT1$modification_sequence%in%RT2$modification_sequence,]
RT2<-RT2[RT2$modification_sequence%in%RT1$modification_sequence,]
RT1<-RT1[!duplicated(RT1$modification_sequence),2:3]
RT2<-RT2[!duplicated(RT2$modification_sequence),2:3]
colnames(RT2)[1]<-"RT_detected_2"
RT<-join(RT1,RT2)
fit <- coef(lm(RT$RT_detected ~ RT$RT_detected_2))
lib2c$RT_detected<-(lib2c$RT_detected*fit[2])+fit[1]
lib2c$iRT<-lib2c$RT_detected

lib2<-rbind(lib2a,lib2b,lib2c)

buildSpectraLibPair(lib1, lib2, outputFile = "ExtendedLib_Kidney.txt", plot = TRUE,
                    clean = FALSE, merge = TRUE, parseAcc = TRUE, consolidateAccession = TRUE,
                    nomod=TRUE, conf.cutoff=0.9)
#2

conf2<-read.xlsx("170324_14_PP_GPlibrary_Kidney__FDR.xlsx", sheet = "Search Summary",
                 colNames = FALSE, rows=33)[1,4]
lib1<-readLibFile("ExtendedLib_Kidney.txt",clean=FALSE)
lib2<-readLibFile("170324_14_PP_GPlibrary_Kidney.txt",clean = TRUE,nomod=TRUE,nomc=FALSE,conf.cutoff=conf2)
lib2<-canonicalFormat(lib2, format = "peakview")
lib2<-lib2[lib2$uniprot_id!="[ RT-Cal protein ]",]
lib2<-lib2[lib2$uniprot_id!="1/[ RT-Cal protein ]",]
lib1<-lib1[!grepl("NA",rownames(lib1)),]
lib2<-lib2[!grepl("NA",rownames(lib2)),]
lib1$iRT<-lib1$RT_detected

# checkQuality(lib1,lib2)

lib2a<-lib2[lib2$RT_detected<87,]
RT1<-lib1[,c(1,3,8)]
RT2<-lib2a[,c(1,3,8)]
RT1<-RT1[RT1$modification_sequence%in%RT2$modification_sequence,]
RT2<-RT2[RT2$modification_sequence%in%RT1$modification_sequence,]
RT1<-RT1[!duplicated(RT1$modification_sequence),2:3]
RT2<-RT2[!duplicated(RT2$modification_sequence),2:3]
colnames(RT2)[1]<-"RT_detected_2"
RT<-join(RT1,RT2)
fit <- coef(lm(RT$RT_detected ~ RT$RT_detected_2))
lib2a$RT_detected<-(lib2a$RT_detected*fit[2])+fit[1]
lib2a$iRT<-lib2a$RT_detected

lib2b<-lib2[lib2$RT_detected>=86,]
RT1<-lib1[,c(1,3,8)]
RT2<-lib2b[,c(1,3,8)]
RT1<-RT1[RT1$modification_sequence%in%RT2$modification_sequence,]
RT2<-RT2[RT2$modification_sequence%in%RT1$modification_sequence,]
RT1<-RT1[!duplicated(RT1$modification_sequence),2:3]
RT2<-RT2[!duplicated(RT2$modification_sequence),2:3]
colnames(RT2)[1]<-"RT_detected_2"
RT<-join(RT1,RT2)
fit <- coef(lm(RT$RT_detected ~ RT$RT_detected_2))
lib2b$RT_detected<-(lib2b$RT_detected*fit[2])+fit[1]
lib2b$iRT<-lib2b$RT_detected

lib2<-rbind(lib2a,lib2b)

buildSpectraLibPair(lib1, lib2, outputFile = "ExtendedLib_Kidney.txt", plot = TRUE,
                    clean = FALSE, merge = TRUE, parseAcc = TRUE, consolidateAccession = TRUE,
                    nomod=TRUE, conf.cutoff=0.9)

x<-read.delim("ExtendedLib_Kidney.txt")
y<-as.data.frame(table(x$stripped_sequence))
y<-y[y$Freq>=5,]
x<-x[x$stripped_sequence%in%y$Var1,]
write.table(x,"ExtendedLib_Kidney.txt",sep="\t")


#Fat
#1

conf1<-read.xlsx("170303_11_PP_GP_Fat__FDR.xlsx", sheet = "Search Summary",
                 colNames = FALSE, rows=33)[1,4]
conf2<-read.xlsx("170310_11_PP_GP_Fat__FDR.xlsx", sheet = "Search Summary",
                 colNames = FALSE, rows=33)[1,4]
lib1<-readLibFile("170303_11_PP_GP_Fat.txt",clean = TRUE,nomod=TRUE,nomc=FALSE, conf.cutoff=conf1)
lib2<-readLibFile("170310_11_PP_GP_Fat.txt",clean = TRUE,nomod=TRUE,nomc=FALSE, conf.cutoff=conf2)
lib1<-canonicalFormat(lib1, format = "peakview")
lib2<-canonicalFormat(lib2, format = "peakview")
lib1<-lib1[lib1$uniprot_id!="[ RT-Cal protein ]",]
lib1<-lib1[lib1$uniprot_id!="1/[ RT-Cal protein ]",]
lib2<-lib2[lib2$uniprot_id!="[ RT-Cal protein ]",]
lib2<-lib2[lib2$uniprot_id!="1/[ RT-Cal protein ]",]
lib1<-lib1[!grepl("NA",rownames(lib1)),]
lib2<-lib2[!grepl("NA",rownames(lib2)),]

# checkQuality(lib1,lib2)

lib2a<-lib2[lib2$RT_detected<60,]
RT1<-lib1[,c(1,3,8)]
RT2<-lib2a[,c(1,3,8)]
RT1<-RT1[RT1$modification_sequence%in%RT2$modification_sequence,]
RT2<-RT2[RT2$modification_sequence%in%RT1$modification_sequence,]
RT1<-RT1[!duplicated(RT1$modification_sequence),2:3]
RT2<-RT2[!duplicated(RT2$modification_sequence),2:3]
colnames(RT2)[1]<-"RT_detected_2"
RT<-join(RT1,RT2)
fit <- coef(lm(RT$RT_detected ~ RT$RT_detected_2))
lib2a$RT_detected<-(lib2a$RT_detected*fit[2])+fit[1]
lib2a$iRT<-lib2a$RT_detected

lib2b<-lib2[lib2$RT_detected>=60,]
lib2b<-lib2b[lib2b$RT_detected<96,]
RT1<-lib1[,c(1,3,8)]
RT2<-lib2b[,c(1,3,8)]
RT1<-RT1[RT1$modification_sequence%in%RT2$modification_sequence,]
RT2<-RT2[RT2$modification_sequence%in%RT1$modification_sequence,]
RT1<-RT1[!duplicated(RT1$modification_sequence),2:3]
RT2<-RT2[!duplicated(RT2$modification_sequence),2:3]
colnames(RT2)[1]<-"RT_detected_2"
RT<-join(RT1,RT2)
fit <- coef(lm(RT$RT_detected ~ RT$RT_detected_2))
lib2b$RT_detected<-(lib2b$RT_detected*fit[2])+fit[1]
lib2b$iRT<-lib2b$RT_detected

lib2c<-lib2[lib2$RT_detected>=95,]
RT1<-lib1[,c(1,3,8)]
RT2<-lib2c[,c(1,3,8)]
RT1<-RT1[RT1$modification_sequence%in%RT2$modification_sequence,]
RT2<-RT2[RT2$modification_sequence%in%RT1$modification_sequence,]
RT1<-RT1[!duplicated(RT1$modification_sequence),2:3]
RT2<-RT2[!duplicated(RT2$modification_sequence),2:3]
colnames(RT2)[1]<-"RT_detected_2"
RT<-join(RT1,RT2)
fit <- coef(lm(RT$RT_detected ~ RT$RT_detected_2))
lib2c$RT_detected<-(lib2c$RT_detected*fit[2])+fit[1]
lib2c$iRT<-lib2c$RT_detected

lib2<-rbind(lib2a,lib2b,lib2c)

buildSpectraLibPair(lib1, lib2, outputFile = "ExtendedLib_Fat.txt", plot = TRUE,
                    clean = FALSE, merge = TRUE, parseAcc = TRUE, consolidateAccession = TRUE,
                    nomod=TRUE, conf.cutoff=0.9)
#2

conf2<-read.xlsx("170324_17_PP_GPlibrary_Fat__FDR.xlsx", sheet = "Search Summary",
                 colNames = FALSE, rows=33)[1,4]
lib1<-readLibFile("ExtendedLib_Fat.txt",clean=FALSE)
lib2<-readLibFile("170324_17_PP_GPlibrary_Fat.txt",clean = TRUE,nomod=TRUE,nomc=FALSE, conf.cutoff=conf2)
lib2<-canonicalFormat(lib2, format = "peakview")
lib2<-lib2[lib2$uniprot_id!="[ RT-Cal protein ]",]
lib2<-lib2[lib2$uniprot_id!="1/[ RT-Cal protein ]",]
lib1<-lib1[!grepl("NA",rownames(lib1)),]
lib2<-lib2[!grepl("NA",rownames(lib2)),]
lib1$iRT<-lib1$RT_detected

# checkQuality(lib1,lib2)

lib2a<-lib2[lib2$RT_detected<87,]
RT1<-lib1[,c(1,3,8)]
RT2<-lib2a[,c(1,3,8)]
RT1<-RT1[RT1$modification_sequence%in%RT2$modification_sequence,]
RT2<-RT2[RT2$modification_sequence%in%RT1$modification_sequence,]
RT1<-RT1[!duplicated(RT1$modification_sequence),2:3]
RT2<-RT2[!duplicated(RT2$modification_sequence),2:3]
colnames(RT2)[1]<-"RT_detected_2"
RT<-join(RT1,RT2)
fit <- coef(lm(RT$RT_detected ~ RT$RT_detected_2))
lib2a$RT_detected<-(lib2a$RT_detected*fit[2])+fit[1]
lib2a$iRT<-lib2a$RT_detected

lib2b<-lib2[lib2$RT_detected>=86,]
RT1<-lib1[,c(1,3,8)]
RT2<-lib2b[,c(1,3,8)]
RT1<-RT1[RT1$modification_sequence%in%RT2$modification_sequence,]
RT2<-RT2[RT2$modification_sequence%in%RT1$modification_sequence,]
RT1<-RT1[!duplicated(RT1$modification_sequence),2:3]
RT2<-RT2[!duplicated(RT2$modification_sequence),2:3]
colnames(RT2)[1]<-"RT_detected_2"
RT<-join(RT1,RT2)
fit <- coef(lm(RT$RT_detected ~ RT$RT_detected_2))
lib2b$RT_detected<-(lib2b$RT_detected*fit[2])+fit[1]
lib2b$iRT<-lib2b$RT_detected

lib2<-rbind(lib2a,lib2b)

buildSpectraLibPair(lib1, lib2, outputFile = "ExtendedLib_Fat.txt", plot = TRUE,
                    clean = FALSE, merge = TRUE, parseAcc = TRUE, consolidateAccession = TRUE,
                    nomod=TRUE, conf.cutoff=0.9)

x<-read.delim("ExtendedLib_Fat.txt")
y<-as.data.frame(table(x$stripped_sequence))
y<-y[y$Freq>=5,]
x<-x[x$stripped_sequence%in%y$Var1,]
write.table(x,"ExtendedLib_Fat.txt",sep="\t")


#Small_Intestine
#1

conf1<-read.xlsx("170306_12_PP_GP_Small_Intestine__FDR.xlsx", sheet = "Search Summary",
                 colNames = FALSE, rows=33)[1,4]
conf2<-read.xlsx("170310_12_PP_GP_Small_Intestine__FDR.xlsx", sheet = "Search Summary",
                 colNames = FALSE, rows=33)[1,4]
lib1<-readLibFile("170306_12_PP_GP_Small_Intestine.txt",clean =TRUE,nomod=TRUE,nomc=FALSE,conf.cutoff=conf1)
lib2<-readLibFile("170310_12_PP_GP_Small_Intestine.txt",clean =TRUE,nomod=TRUE,nomc=FALSE,conf.cutoff=conf2)
lib1<-canonicalFormat(lib1, format = "peakview")
lib2<-canonicalFormat(lib2, format = "peakview")
lib1<-lib1[lib1$uniprot_id!="[ RT-Cal protein ]",]
lib1<-lib1[lib1$uniprot_id!="1/[ RT-Cal protein ]",]
lib2<-lib2[lib2$uniprot_id!="[ RT-Cal protein ]",]
lib2<-lib2[lib2$uniprot_id!="1/[ RT-Cal protein ]",]
lib1<-lib1[!grepl("NA",rownames(lib1)),]
lib2<-lib2[!grepl("NA",rownames(lib2)),]

# checkQuality(lib1,lib2)

lib2a<-lib2[lib2$RT_detected<60,]
RT1<-lib1[,c(1,3,8)]
RT2<-lib2a[,c(1,3,8)]
RT1<-RT1[RT1$modification_sequence%in%RT2$modification_sequence,]
RT2<-RT2[RT2$modification_sequence%in%RT1$modification_sequence,]
RT1<-RT1[!duplicated(RT1$modification_sequence),2:3]
RT2<-RT2[!duplicated(RT2$modification_sequence),2:3]
colnames(RT2)[1]<-"RT_detected_2"
RT<-join(RT1,RT2)
fit <- coef(lm(RT$RT_detected ~ RT$RT_detected_2))
lib2a$RT_detected<-(lib2a$RT_detected*fit[2])+fit[1]
lib2a$iRT<-lib2a$RT_detected

lib2b<-lib2[lib2$RT_detected>=60,]
lib2b<-lib2b[lib2b$RT_detected<96,]
RT1<-lib1[,c(1,3,8)]
RT2<-lib2b[,c(1,3,8)]
RT1<-RT1[RT1$modification_sequence%in%RT2$modification_sequence,]
RT2<-RT2[RT2$modification_sequence%in%RT1$modification_sequence,]
RT1<-RT1[!duplicated(RT1$modification_sequence),2:3]
RT2<-RT2[!duplicated(RT2$modification_sequence),2:3]
colnames(RT2)[1]<-"RT_detected_2"
RT<-join(RT1,RT2)
fit <- coef(lm(RT$RT_detected ~ RT$RT_detected_2))
lib2b$RT_detected<-(lib2b$RT_detected*fit[2])+fit[1]
lib2b$iRT<-lib2b$RT_detected

lib2c<-lib2[lib2$RT_detected>=95,]
RT1<-lib1[,c(1,3,8)]
RT2<-lib2c[,c(1,3,8)]
RT1<-RT1[RT1$modification_sequence%in%RT2$modification_sequence,]
RT2<-RT2[RT2$modification_sequence%in%RT1$modification_sequence,]
RT1<-RT1[!duplicated(RT1$modification_sequence),2:3]
RT2<-RT2[!duplicated(RT2$modification_sequence),2:3]
colnames(RT2)[1]<-"RT_detected_2"
RT<-join(RT1,RT2)
fit <- coef(lm(RT$RT_detected ~ RT$RT_detected_2))
lib2c$RT_detected<-(lib2c$RT_detected*fit[2])+fit[1]
lib2c$iRT<-lib2c$RT_detected

lib2<-rbind(lib2a,lib2b,lib2c)

buildSpectraLibPair(lib1, lib2, outputFile = "ExtendedLib_Small_Intestine.txt", plot = TRUE,
                    clean = FALSE, merge = TRUE, parseAcc = TRUE, consolidateAccession = TRUE,
                    nomod=TRUE, conf.cutoff=0.9)

x<-read.delim("ExtendedLib_Small_Intestine.txt")
y<-as.data.frame(table(x$stripped_sequence))
y<-y[y$Freq>=5,]
x<-x[x$stripped_sequence%in%y$Var1,]
write.table(x,"ExtendedLib_Small_Intestine.txt",sep="\t")


#Lung
#1

conf1<-read.xlsx("170306_13_PP_GP_Lung__FDR.xlsx", sheet = "Search Summary",
                 colNames = FALSE, rows=33)[1,4]
conf2<-read.xlsx("170310_13_PP_GP_Lung__FDR.xlsx", sheet = "Search Summary",
                 colNames = FALSE, rows=33)[1,4]
lib1<-readLibFile("170306_13_PP_GP_Lung.txt",clean = TRUE,nomod=TRUE,nomc=FALSE, conf.cutoff=conf1)
lib2<-readLibFile("170310_13_PP_GP_Lung.txt",clean = TRUE,nomod=TRUE,nomc=FALSE, conf.cutoff=conf2)
lib1<-canonicalFormat(lib1, format = "peakview")
lib2<-canonicalFormat(lib2, format = "peakview")
lib1<-lib1[lib1$uniprot_id!="[ RT-Cal protein ]",]
lib1<-lib1[lib1$uniprot_id!="1/[ RT-Cal protein ]",]
lib2<-lib2[lib2$uniprot_id!="[ RT-Cal protein ]",]
lib2<-lib2[lib2$uniprot_id!="1/[ RT-Cal protein ]",]
lib1<-lib1[!grepl("NA",rownames(lib1)),]
lib2<-lib2[!grepl("NA",rownames(lib2)),]

# checkQuality(lib1,lib2)

lib2a<-lib2[lib2$RT_detected<60,]
RT1<-lib1[,c(1,3,8)]
RT2<-lib2a[,c(1,3,8)]
RT1<-RT1[RT1$modification_sequence%in%RT2$modification_sequence,]
RT2<-RT2[RT2$modification_sequence%in%RT1$modification_sequence,]
RT1<-RT1[!duplicated(RT1$modification_sequence),2:3]
RT2<-RT2[!duplicated(RT2$modification_sequence),2:3]
colnames(RT2)[1]<-"RT_detected_2"
RT<-join(RT1,RT2)
fit <- coef(lm(RT$RT_detected ~ RT$RT_detected_2))
lib2a$RT_detected<-(lib2a$RT_detected*fit[2])+fit[1]
lib2a$iRT<-lib2a$RT_detected

lib2b<-lib2[lib2$RT_detected>=60,]
lib2b<-lib2b[lib2b$RT_detected<96,]
RT1<-lib1[,c(1,3,8)]
RT2<-lib2b[,c(1,3,8)]
RT1<-RT1[RT1$modification_sequence%in%RT2$modification_sequence,]
RT2<-RT2[RT2$modification_sequence%in%RT1$modification_sequence,]
RT1<-RT1[!duplicated(RT1$modification_sequence),2:3]
RT2<-RT2[!duplicated(RT2$modification_sequence),2:3]
colnames(RT2)[1]<-"RT_detected_2"
RT<-join(RT1,RT2)
fit <- coef(lm(RT$RT_detected ~ RT$RT_detected_2))
lib2b$RT_detected<-(lib2b$RT_detected*fit[2])+fit[1]
lib2b$iRT<-lib2b$RT_detected

lib2c<-lib2[lib2$RT_detected>=95,]
RT1<-lib1[,c(1,3,8)]
RT2<-lib2c[,c(1,3,8)]
RT1<-RT1[RT1$modification_sequence%in%RT2$modification_sequence,]
RT2<-RT2[RT2$modification_sequence%in%RT1$modification_sequence,]
RT1<-RT1[!duplicated(RT1$modification_sequence),2:3]
RT2<-RT2[!duplicated(RT2$modification_sequence),2:3]
colnames(RT2)[1]<-"RT_detected_2"
RT<-join(RT1,RT2)
fit <- coef(lm(RT$RT_detected ~ RT$RT_detected_2))
lib2c$RT_detected<-(lib2c$RT_detected*fit[2])+fit[1]
lib2c$iRT<-lib2c$RT_detected

lib2<-rbind(lib2a,lib2b,lib2c)

buildSpectraLibPair(lib1, lib2, outputFile = "ExtendedLib_Lung.txt", plot = TRUE,
                    clean = FALSE, merge = TRUE, parseAcc = TRUE, consolidateAccession = TRUE,
                    nomod=TRUE, conf.cutoff=0.9)


x<-read.delim("ExtendedLib_Lung.txt")
y<-as.data.frame(table(x$stripped_sequence))
y<-y[y$Freq>=5,]
x<-x[x$stripped_sequence%in%y$Var1,]
write.table(x,"ExtendedLib_Lung.txt",sep="\t")


#Liver
#1

conf1<-read.xlsx("170307_15_PP_GP_Liver__FDR.xlsx", sheet = "Search Summary",
                 colNames = FALSE, rows=33)[1,4]
conf2<-read.xlsx("170310_15_PP_GP_Liver__FDR.xlsx", sheet = "Search Summary",
                 colNames = FALSE, rows=33)[1,4]
lib1<-readLibFile("170307_15_PP_GP_Liver.txt",clean = TRUE,nomod=TRUE,nomc=FALSE, conf.cutoff=conf1)
lib2<-readLibFile("170310_15_PP_GP_Liver.txt",clean = TRUE,nomod=TRUE,nomc=FALSE, conf.cutoff=conf2)
lib1<-canonicalFormat(lib1, format = "peakview")
lib2<-canonicalFormat(lib2, format = "peakview")
lib1<-lib1[lib1$uniprot_id!="[ RT-Cal protein ]",]
lib1<-lib1[lib1$uniprot_id!="1/[ RT-Cal protein ]",]
lib2<-lib2[lib2$uniprot_id!="[ RT-Cal protein ]",]
lib2<-lib2[lib2$uniprot_id!="1/[ RT-Cal protein ]",]
lib1<-lib1[!grepl("NA",rownames(lib1)),]
lib2<-lib2[!grepl("NA",rownames(lib2)),]


# checkQuality(lib1,lib2)

lib2a<-lib2[lib2$RT_detected<60,]
RT1<-lib1[,c(1,3,8)]
RT2<-lib2a[,c(1,3,8)]
RT1<-RT1[RT1$modification_sequence%in%RT2$modification_sequence,]
RT2<-RT2[RT2$modification_sequence%in%RT1$modification_sequence,]
RT1<-RT1[!duplicated(RT1$modification_sequence),2:3]
RT2<-RT2[!duplicated(RT2$modification_sequence),2:3]
colnames(RT2)[1]<-"RT_detected_2"
RT<-join(RT1,RT2)
fit <- coef(lm(RT$RT_detected ~ RT$RT_detected_2))
lib2a$RT_detected<-(lib2a$RT_detected*fit[2])+fit[1]
lib2a$iRT<-lib2a$RT_detected

lib2b<-lib2[lib2$RT_detected>=60,]
lib2b<-lib2b[lib2b$RT_detected<96,]
RT1<-lib1[,c(1,3,8)]
RT2<-lib2b[,c(1,3,8)]
RT1<-RT1[RT1$modification_sequence%in%RT2$modification_sequence,]
RT2<-RT2[RT2$modification_sequence%in%RT1$modification_sequence,]
RT1<-RT1[!duplicated(RT1$modification_sequence),2:3]
RT2<-RT2[!duplicated(RT2$modification_sequence),2:3]
colnames(RT2)[1]<-"RT_detected_2"
RT<-join(RT1,RT2)
fit <- coef(lm(RT$RT_detected ~ RT$RT_detected_2))
lib2b$RT_detected<-(lib2b$RT_detected*fit[2])+fit[1]
lib2b$iRT<-lib2b$RT_detected

lib2c<-lib2[lib2$RT_detected>=95,]
RT1<-lib1[,c(1,3,8)]
RT2<-lib2c[,c(1,3,8)]
RT1<-RT1[RT1$modification_sequence%in%RT2$modification_sequence,]
RT2<-RT2[RT2$modification_sequence%in%RT1$modification_sequence,]
RT1<-RT1[!duplicated(RT1$modification_sequence),2:3]
RT2<-RT2[!duplicated(RT2$modification_sequence),2:3]
colnames(RT2)[1]<-"RT_detected_2"
RT<-join(RT1,RT2)
fit <- coef(lm(RT$RT_detected ~ RT$RT_detected_2))
lib2c$RT_detected<-(lib2c$RT_detected*fit[2])+fit[1]
lib2c$iRT<-lib2c$RT_detected

lib2<-rbind(lib2a,lib2b,lib2c)

buildSpectraLibPair(lib1, lib2, outputFile = "ExtendedLib_Liver.txt", plot = TRUE,
                    clean = FALSE, merge = TRUE, parseAcc = TRUE, consolidateAccession = TRUE,
                    nomod=TRUE, conf.cutoff=0.9)
#2

conf2<-read.xlsx("170324_20_PP_GPlibrary_Liver__FDR.xlsx", sheet = "Search Summary",
                 colNames = FALSE, rows=33)[1,4]
lib1<-readLibFile("ExtendedLib_Liver.txt",clean=FALSE)
lib2<-readLibFile("170324_20_PP_GPlibrary_Liver.txt",clean = TRUE,nomod=TRUE,nomc=FALSE, conf.cutoff=conf2)
lib2<-canonicalFormat(lib2, format = "peakview")
lib2<-lib2[lib2$uniprot_id!="[ RT-Cal protein ]",]
lib2<-lib2[lib2$uniprot_id!="1/[ RT-Cal protein ]",]
lib1<-lib1[!grepl("NA",rownames(lib1)),]
lib2<-lib2[!grepl("NA",rownames(lib2)),]
lib1$iRT<-lib1$RT_detected

# checkQuality(lib1,lib2)

lib2a<-lib2[lib2$RT_detected<87,]
RT1<-lib1[,c(1,3,8)]
RT2<-lib2a[,c(1,3,8)]
RT1<-RT1[RT1$modification_sequence%in%RT2$modification_sequence,]
RT2<-RT2[RT2$modification_sequence%in%RT1$modification_sequence,]
RT1<-RT1[!duplicated(RT1$modification_sequence),2:3]
RT2<-RT2[!duplicated(RT2$modification_sequence),2:3]
colnames(RT2)[1]<-"RT_detected_2"
RT<-join(RT1,RT2)
fit <- coef(lm(RT$RT_detected ~ RT$RT_detected_2))
lib2a$RT_detected<-(lib2a$RT_detected*fit[2])+fit[1]
lib2a$iRT<-lib2a$RT_detected

lib2b<-lib2[lib2$RT_detected>=86,]
RT1<-lib1[,c(1,3,8)]
RT2<-lib2b[,c(1,3,8)]
RT1<-RT1[RT1$modification_sequence%in%RT2$modification_sequence,]
RT2<-RT2[RT2$modification_sequence%in%RT1$modification_sequence,]
RT1<-RT1[!duplicated(RT1$modification_sequence),2:3]
RT2<-RT2[!duplicated(RT2$modification_sequence),2:3]
colnames(RT2)[1]<-"RT_detected_2"
RT<-join(RT1,RT2)
fit <- coef(lm(RT$RT_detected ~ RT$RT_detected_2))
lib2b$RT_detected<-(lib2b$RT_detected*fit[2])+fit[1]
lib2b$iRT<-lib2b$RT_detected

lib2<-rbind(lib2a,lib2b)

buildSpectraLibPair(lib1, lib2, outputFile = "ExtendedLib_Liver.txt", plot = TRUE,
                    clean = FALSE, merge = TRUE, parseAcc = TRUE, consolidateAccession = TRUE,
                    nomod=TRUE, conf.cutoff=0.9)


x<-read.delim("ExtendedLib_Liver.txt")
y<-as.data.frame(table(x$stripped_sequence))
y<-y[y$Freq>=5,]
x<-x[x$stripped_sequence%in%y$Var1,]
write.table(x,"ExtendedLib_Liver.txt",sep="\t")

#Overies

#1

conf1<-read.xlsx("170307_16_PP_GP_Overies__FDR.xlsx", sheet = "Search Summary",
                 colNames = FALSE, rows=33)[1,4]
conf2<-read.xlsx("170310_16_PP_GP_Overies__FDR.xlsx", sheet = "Search Summary",
                 colNames = FALSE, rows=33)[1,4]
lib1<-readLibFile("170307_16_PP_GP_Overies.txt",clean = TRUE,nomod=TRUE,nomc=FALSE, conf.cutoff=conf1)
lib2<-readLibFile("170310_16_PP_GP_Overies.txt",clean = TRUE,nomod=TRUE,nomc=FALSE, conf.cutoff=conf2)
lib1<-canonicalFormat(lib1, format = "peakview")
lib2<-canonicalFormat(lib2, format = "peakview")
lib1<-lib1[lib1$uniprot_id!="[ RT-Cal protein ]",]
lib1<-lib1[lib1$uniprot_id!="1/[ RT-Cal protein ]",]
lib2<-lib2[lib2$uniprot_id!="[ RT-Cal protein ]",]
lib2<-lib2[lib2$uniprot_id!="1/[ RT-Cal protein ]",]
lib1<-lib1[!grepl("NA",rownames(lib1)),]
lib2<-lib2[!grepl("NA",rownames(lib2)),]

# checkQuality(lib1,lib2)

lib2a<-lib2[lib2$RT_detected<60,]
RT1<-lib1[,c(1,3,8)]
RT2<-lib2a[,c(1,3,8)]
RT1<-RT1[RT1$modification_sequence%in%RT2$modification_sequence,]
RT2<-RT2[RT2$modification_sequence%in%RT1$modification_sequence,]
RT1<-RT1[!duplicated(RT1$modification_sequence),2:3]
RT2<-RT2[!duplicated(RT2$modification_sequence),2:3]
colnames(RT2)[1]<-"RT_detected_2"
RT<-join(RT1,RT2)
fit <- coef(lm(RT$RT_detected ~ RT$RT_detected_2))
lib2a$RT_detected<-(lib2a$RT_detected*fit[2])+fit[1]
lib2a$iRT<-lib2a$RT_detected

lib2b<-lib2[lib2$RT_detected>=60,]
lib2b<-lib2b[lib2b$RT_detected<96,]
RT1<-lib1[,c(1,3,8)]
RT2<-lib2b[,c(1,3,8)]
RT1<-RT1[RT1$modification_sequence%in%RT2$modification_sequence,]
RT2<-RT2[RT2$modification_sequence%in%RT1$modification_sequence,]
RT1<-RT1[!duplicated(RT1$modification_sequence),2:3]
RT2<-RT2[!duplicated(RT2$modification_sequence),2:3]
colnames(RT2)[1]<-"RT_detected_2"
RT<-join(RT1,RT2)
fit <- coef(lm(RT$RT_detected ~ RT$RT_detected_2))
lib2b$RT_detected<-(lib2b$RT_detected*fit[2])+fit[1]
lib2b$iRT<-lib2b$RT_detected

lib2c<-lib2[lib2$RT_detected>=95,]
RT1<-lib1[,c(1,3,8)]
RT2<-lib2c[,c(1,3,8)]
RT1<-RT1[RT1$modification_sequence%in%RT2$modification_sequence,]
RT2<-RT2[RT2$modification_sequence%in%RT1$modification_sequence,]
RT1<-RT1[!duplicated(RT1$modification_sequence),2:3]
RT2<-RT2[!duplicated(RT2$modification_sequence),2:3]
colnames(RT2)[1]<-"RT_detected_2"
RT<-join(RT1,RT2)
fit <- coef(lm(RT$RT_detected ~ RT$RT_detected_2))
lib2c$RT_detected<-(lib2c$RT_detected*fit[2])+fit[1]
lib2c$iRT<-lib2c$RT_detected

lib2<-rbind(lib2a,lib2b,lib2c)

buildSpectraLibPair(lib1, lib2, outputFile = "ExtendedLib_Ovaries.txt", plot = TRUE,
                    clean = FALSE, merge = TRUE, parseAcc = TRUE, consolidateAccession = TRUE,
                    nomod=TRUE, conf.cutoff=0.9)
#2

conf2<-read.xlsx("170324_21_PP_GPlibrary_Ovaries__FDR.xlsx", sheet = "Search Summary",
                 colNames = FALSE, rows=33)[1,4]
lib1<-readLibFile("ExtendedLib_Ovaries.txt",clean=FALSE)
lib2<-readLibFile("170324_21_PP_GPlibrary_Ovaries.txt",clean = TRUE,nomod=TRUE,nomc=FALSE,conf.cutoff=conf2)
lib2<-canonicalFormat(lib2, format = "peakview")
lib2<-lib2[lib2$uniprot_id!="[ RT-Cal protein ]",]
lib2<-lib2[lib2$uniprot_id!="1/[ RT-Cal protein ]",]
lib1<-lib1[!grepl("NA",rownames(lib1)),]
lib2<-lib2[!grepl("NA",rownames(lib2)),]
lib1$iRT<-lib1$RT_detected

# checkQuality(lib1,lib2)

lib2a<-lib2[lib2$RT_detected<87,]
RT1<-lib1[,c(1,3,8)]
RT2<-lib2a[,c(1,3,8)]
RT1<-RT1[RT1$modification_sequence%in%RT2$modification_sequence,]
RT2<-RT2[RT2$modification_sequence%in%RT1$modification_sequence,]
RT1<-RT1[!duplicated(RT1$modification_sequence),2:3]
RT2<-RT2[!duplicated(RT2$modification_sequence),2:3]
colnames(RT2)[1]<-"RT_detected_2"
RT<-join(RT1,RT2)
fit <- coef(lm(RT$RT_detected ~ RT$RT_detected_2))
lib2a$RT_detected<-(lib2a$RT_detected*fit[2])+fit[1]
lib2a$iRT<-lib2a$RT_detected

lib2b<-lib2[lib2$RT_detected>=86,]
RT1<-lib1[,c(1,3,8)]
RT2<-lib2b[,c(1,3,8)]
RT1<-RT1[RT1$modification_sequence%in%RT2$modification_sequence,]
RT2<-RT2[RT2$modification_sequence%in%RT1$modification_sequence,]
RT1<-RT1[!duplicated(RT1$modification_sequence),2:3]
RT2<-RT2[!duplicated(RT2$modification_sequence),2:3]
colnames(RT2)[1]<-"RT_detected_2"
RT<-join(RT1,RT2)
fit <- coef(lm(RT$RT_detected ~ RT$RT_detected_2))
lib2b$RT_detected<-(lib2b$RT_detected*fit[2])+fit[1]
lib2b$iRT<-lib2b$RT_detected

lib2<-rbind(lib2a,lib2b)

buildSpectraLibPair(lib1, lib2, outputFile = "ExtendedLib_Ovaries.txt", plot = TRUE,
                    clean = FALSE, merge = TRUE, parseAcc = TRUE, consolidateAccession = TRUE,
                    nomod=TRUE, conf.cutoff=0.9)

x<-read.delim("ExtendedLib_Ovaries.txt")
y<-as.data.frame(table(x$stripped_sequence))
y<-y[y$Freq>=5,]
x<-x[x$stripped_sequence%in%y$Var1,]
write.table(x,"ExtendedLib_Ovaries.txt",sep="\t")

#SkeletalMusc

#1

conf1<-read.xlsx("170320_08_PP_SkeletalMusc_FASP__FDR.xlsx", sheet = "Search Summary",
                 colNames = FALSE, rows=33)[1,4]
conf2<-read.xlsx("170323_02_GPMuscle_NEW__FDR.xlsx", sheet = "Search Summary",
                 colNames = FALSE, rows=33)[1,4]
lib1<-readLibFile("170320_08_PP_SkeletalMusc_FASP.txt",clean =TRUE,nomod=TRUE,nomc=FALSE, conf.cutoff=conf1)
lib2<-readLibFile("170323_02_GPMuscle_NEW.txt",clean = TRUE,nomod=TRUE,nomc=FALSE, conf.cutoff=conf2)
lib1<-canonicalFormat(lib1, format = "peakview")
lib2<-canonicalFormat(lib2, format = "peakview")
lib1<-lib1[lib1$uniprot_id!="[ RT-Cal protein ]",]
lib1<-lib1[lib1$uniprot_id!="1/[ RT-Cal protein ]",]
lib2<-lib2[lib2$uniprot_id!="[ RT-Cal protein ]",]
lib2<-lib2[lib2$uniprot_id!="1/[ RT-Cal protein ]",]
lib1<-lib1[!grepl("NA",rownames(lib1)),]
lib2<-lib2[!grepl("NA",rownames(lib2)),]

checkQuality(lib1,lib2)

lib2a<-lib2[lib2$RT_detected<95,]
RT1<-lib1[,c(1,3,8)]
RT2<-lib2a[,c(1,3,8)]
RT1<-RT1[RT1$modification_sequence%in%RT2$modification_sequence,]
RT2<-RT2[RT2$modification_sequence%in%RT1$modification_sequence,]
RT1<-RT1[!duplicated(RT1$modification_sequence),2:3]
RT2<-RT2[!duplicated(RT2$modification_sequence),2:3]
colnames(RT2)[1]<-"RT_detected_2"
RT<-join(RT1,RT2)
fit <- coef(lm(RT$RT_detected ~ RT$RT_detected_2))
lib2a$RT_detected<-(lib2a$RT_detected*fit[2])+fit[1]
lib2a$iRT<-lib2a$RT_detected

lib2b<-lib2[lib2$RT_detected>=95,]
RT1<-lib1[,c(1,3,8)]
RT2<-lib2b[,c(1,3,8)]
RT1<-RT1[RT1$modification_sequence%in%RT2$modification_sequence,]
RT2<-RT2[RT2$modification_sequence%in%RT1$modification_sequence,]
RT1<-RT1[!duplicated(RT1$modification_sequence),2:3]
RT2<-RT2[!duplicated(RT2$modification_sequence),2:3]
colnames(RT2)[1]<-"RT_detected_2"
RT<-join(RT1,RT2)
fit <- coef(lm(RT$RT_detected ~ RT$RT_detected_2))
lib2b$RT_detected<-(lib2b$RT_detected*fit[2])+fit[1]
lib2b$iRT<-lib2b$RT_detected

lib2<-rbind(lib2a,lib2b)

buildSpectraLibPair(lib1, lib2, outputFile = "ExtendedLib_SkeletalMusc.txt", plot = TRUE,
                    clean = FALSE, merge = TRUE, parseAcc = TRUE, consolidateAccession = TRUE,
                    nomod=TRUE, conf.cutoff=0.9)
#2

conf2<-read.xlsx("170323_03_GPMuscle_OLD__FDR.xlsx", sheet = "Search Summary",
                 colNames = FALSE, rows=33)[1,4]
lib1<-readLibFile("ExtendedLib_SkeletalMusc.txt",clean=FALSE)
lib2<-readLibFile("170323_03_GPMuscle_OLD.txt",clean = TRUE,nomod=TRUE,nomc=FALSE, conf.cutoff=conf2)
lib2<-canonicalFormat(lib2, format = "peakview")
lib2<-lib2[lib2$uniprot_id!="[ RT-Cal protein ]",]
lib2<-lib2[lib2$uniprot_id!="1/[ RT-Cal protein ]",]
lib1<-lib1[!grepl("NA",rownames(lib1)),]
lib2<-lib2[!grepl("NA",rownames(lib2)),]
lib1$iRT<-lib1$RT_detected

checkQuality(lib1,lib2)
lib2a<-lib2[lib2$RT_detected<95,]
RT1<-lib1[,c(1,3,8)]
RT2<-lib2a[,c(1,3,8)]
RT1<-RT1[RT1$modification_sequence%in%RT2$modification_sequence,]
RT2<-RT2[RT2$modification_sequence%in%RT1$modification_sequence,]
RT1<-RT1[!duplicated(RT1$modification_sequence),2:3]
RT2<-RT2[!duplicated(RT2$modification_sequence),2:3]
colnames(RT2)[1]<-"RT_detected_2"
RT<-join(RT1,RT2)
fit <- coef(lm(RT$RT_detected ~ RT$RT_detected_2))
lib2a$RT_detected<-(lib2a$RT_detected*fit[2])+fit[1]
lib2a$iRT<-lib2a$RT_detected

lib2b<-lib2[lib2$RT_detected>=95,]
RT1<-lib1[,c(1,3,8)]
RT2<-lib2b[,c(1,3,8)]
RT1<-RT1[RT1$modification_sequence%in%RT2$modification_sequence,]
RT2<-RT2[RT2$modification_sequence%in%RT1$modification_sequence,]
RT1<-RT1[!duplicated(RT1$modification_sequence),2:3]
RT2<-RT2[!duplicated(RT2$modification_sequence),2:3]
colnames(RT2)[1]<-"RT_detected_2"
RT<-join(RT1,RT2)
fit <- coef(lm(RT$RT_detected ~ RT$RT_detected_2))
lib2b$RT_detected<-(lib2b$RT_detected*fit[2])+fit[1]
lib2b$iRT<-lib2b$RT_detected

lib2<-rbind(lib2a,lib2b)

buildSpectraLibPair(lib1, lib2, outputFile = "ExtendedLib_SkeletalMusc.txt", plot = TRUE,
                    clean = FALSE, merge = TRUE, parseAcc = TRUE, consolidateAccession = TRUE,
                    nomod=TRUE, conf.cutoff=0.9)


x<-read.delim("ExtendedLib_SkeletalMusc.txt")
y<-as.data.frame(table(x$stripped_sequence))
y<-y[y$Freq>=5,]
x<-x[x$stripped_sequence%in%y$Var1,]
write.table(x,"ExtendedLib_SkeletalMusc.txt",sep="\t")
```
