## Supplementary material for "The generation of a comprehensive spectral library for the analysis of the guinea pig proteome by SWATH-MS": R script 2

```
#Individual search results (.group files) were converted into spectral library format Using PeakView software package with SWATH microApp and saved as .txt

#Guinea pig heart spectral library is being assembled here using multiple idividual search results.

library(SwathXtend)
library(plyr)
library(openxlsx)

workDir<-getwd()

#1

#Retrieving the conf.cutoff, corresponding to FDR<0.01, from the search summarry in .xlsx format
conf1<-read.xlsx("Heart__FDR.xlsx", sheet = "Search Summary", colNames = FALSE, rows=33)[1,4]
conf2<-read.xlsx("Newcastle_1__FDR.xlsx", sheet = "Search Summary", colNames = FALSE, rows=33)[1,4]

#Loading and cleaning the library files
lib1<-readLibFile("Heart.txt", clean = TRUE, intensity.cutoff = 5, conf.cutoff = conf1,
                  nomod = TRUE, nomc = FALSE, enz = "trypsin")
lib2<-readLibFile("Newcastle_1.txt", clean = TRUE, intensity.cutoff = 5, conf.cutoff = conf2,
                  nomod = TRUE, nomc = FALSE, enz = "trypsin")
#correcting the file format (removing the retension time calibrant and nonannotated peptides)
lib1<-canonicalFormat(lib1, format = "peakview")
lib2<-canonicalFormat(lib2, format = "peakview")
lib1<-lib1[lib1$uniprot_id!="[ RT-Cal protein ]",]
lib1<-lib1[lib1$uniprot_id!="1/[ RT-Cal protein ]",]
lib2<-lib2[lib2$uniprot_id!="[ RT-Cal protein ]",]
lib2<-lib2[lib2$uniprot_id!="1/[ RT-Cal protein ]",]
lib1<-lib1[!grepl("NA",rownames(lib1)),]
lib2<-lib2[!grepl("NA",rownames(lib2)),]

#creating a directory for the output files (quality measures for the first step of the library assembly)
dir.create("1")
setwd("1")

# Checking the linearity of RT correlation. If nonlinear, the gradient is divided to linear fragments. 
# QC measures get saved in dedicated folder

checkQuality(lib1,lib2)

#based on the visual inspection, the retension time correlation is non-linear
#The External library lib2 (the one that is being added to the "base library" lib1) is therefore manually divided into few, more linear fragments (lib2a-lib2c)

#1corr

#Linear fragment one

lib2a<-lib2[lib2$RT_detected<30,]

RT1<-lib1[,c(1,3,8)]
RT2<-lib2a[,c(1,3,8)]
RT1<-RT1[RT1$modification_sequence%in%RT2$modification_sequence,]
RT2<-RT2[RT2$modification_sequence%in%RT1$modification_sequence,]
RT1<-RT1[!duplicated(RT1$modification_sequence),2:3]
RT2<-RT2[!duplicated(RT2$modification_sequence),2:3]
colnames(RT2)[1]<-"RT_detected_2"
RT<-join(RT1,RT2)

# Linear regression is calculated and the coeficients used to correct the external library RT to mach the base library

fit <- coef(lm(RT$RT_detected ~ RT$RT_detected_2))
lib2a$RT_detected<-(lib2a$RT_detected*fit[2])+fit[1]
lib2a$iRT<-lib2a$RT_detected

#Linear fragment 2

lib2b<-lib2[lib2$RT_detected>=30,]
lib2b<-lib2b[lib2b$RT_detected<115,]

RT1<-lib1[,c(1,3,8)]
RT2<-lib2b[,c(1,3,8)]
RT1<-RT1[RT1$modification_sequence%in%RT2$modification_sequence,]
RT2<-RT2[RT2$modification_sequence%in%RT1$modification_sequence,]
RT1<-RT1[!duplicated(RT1$modification_sequence),2:3]
RT2<-RT2[!duplicated(RT2$modification_sequence),2:3]
colnames(RT2)[1]<-"RT_detected_2"
RT<-join(RT1,RT2)
fit <- coef(lm(RT$RT_detected ~ RT$RT_detected_2))
lib2b$RT_detected<-(lib2b$RT_detected*fit[2])+fit[1]
lib2b$iRT<-lib2b$RT_detected

#Linear fragment 3

lib2c<-lib2[lib2$RT_detected>115,]

RT1<-lib1[,c(1,3,8)]
RT2<-lib2c[,c(1,3,8)]
RT1<-RT1[RT1$modification_sequence%in%RT2$modification_sequence,]
RT2<-RT2[RT2$modification_sequence%in%RT1$modification_sequence,]
RT1<-RT1[!duplicated(RT1$modification_sequence),2:3]
RT2<-RT2[!duplicated(RT2$modification_sequence),2:3]
colnames(RT2)[1]<-"RT_detected_2"
RT<-join(RT1,RT2)
fit <- coef(lm(RT$RT_detected ~ RT$RT_detected_2))
lib2c$RT_detected<-(lib2c$RT_detected*fit[2])+fit[1]
lib2c$iRT<-lib2c$RT_detected

#Linear fragments are then reassembled  

lib2<-rbind(lib2a,lib2b,lib2c)

#A subdirectory is created to save the output of merging process of the linearized external library with the base library

dir.create("1corr")
setwd("1corr")

#Libraries are merged. QC measures get saved in dedicated subfolder. No cleaning is required at this step since both libraries were cleaned already.

buildSpectraLibPair(lib1, lib2, outputFile = "ExtendedLib_heart.txt", plot = TRUE,
                    clean = FALSE, merge = TRUE, parseAcc = TRUE, consolidateAccession = TRUE, 
                    intensity.cutoff = 5, conf.cutoff = 0.5, nomod = TRUE, nomc = FALSE, enz = "trypsin")

# The merged library is moved to the working directory and will be used as the base library in the following steps.

file.rename(from=paste(getwd(),"/ExtendedLib_heart.txt",sep=""), to=paste(workDir,"/ExtendedLib_heart.txt", sep=""))
setwd(workDir)

#2
#The process described above is then repeated untill all the partial libraries are joined.
#In the followinf steps, only the external library needs to be cleaned.

conf2<-read.xlsx("Newcastle_3__FDR.xlsx", sheet = "Search Summary", colNames = FALSE, rows=33)[1,4]
lib1<-readLibFile("ExtendedLib_heart.txt", clean = FALSE)
lib2<-readLibFile("Newcastle_3.txt", clean = TRUE, intensity.cutoff = 5, conf.cutoff = conf2,
                  nomod = TRUE, nomc = FALSE, enz = "trypsin")
lib2<-canonicalFormat(lib2, format = "peakview")
lib2<-lib2[lib2$uniprot_id!="[ RT-Cal protein ]",]
lib2<-lib2[lib2$uniprot_id!="1/[ RT-Cal protein ]",]
lib2<-lib2[!grepl("NA",rownames(lib2)),]
lib1$iRT<-lib1$RT_detected

dir.create("2")
setwd("2")
checkQuality(lib1,lib2)


#2corr

lib2a<-lib2[lib2$RT_detected<30,]

RT1<-lib1[,c(1,3,8)]
RT2<-lib2a[,c(1,3,8)]
RT1<-RT1[RT1$modification_sequence%in%RT2$modification_sequence,]
RT2<-RT2[RT2$modification_sequence%in%RT1$modification_sequence,]
RT1<-RT1[!duplicated(RT1$modification_sequence),2:3]
RT2<-RT2[!duplicated(RT2$modification_sequence),2:3]
colnames(RT2)[1]<-"RT_detected_2"
RT<-join(RT1,RT2)
fit <- coef(lm(RT$RT_detected ~ RT$RT_detected_2))
lib2a$RT_detected<-(lib2a$RT_detected*fit[2])+fit[1]
lib2a$iRT<-lib2a$RT_detected

lib2b<-lib2[lib2$RT_detected>=30,]
lib2b<-lib2b[lib2b$RT_detected<40,]

RT1<-lib1[,c(1,3,8)]
RT2<-lib2b[,c(1,3,8)]
RT1<-RT1[RT1$modification_sequence%in%RT2$modification_sequence,]
RT2<-RT2[RT2$modification_sequence%in%RT1$modification_sequence,]
RT1<-RT1[!duplicated(RT1$modification_sequence),2:3]
RT2<-RT2[!duplicated(RT2$modification_sequence),2:3]
colnames(RT2)[1]<-"RT_detected_2"
RT<-join(RT1,RT2)
fit <- coef(lm(RT$RT_detected ~ RT$RT_detected_2))
lib2b$RT_detected<-(lib2b$RT_detected*fit[2])+fit[1]
lib2b$iRT<-lib2b$RT_detected

lib2c<-lib2[lib2$RT_detected>=40,]
lib2c<-lib2c[lib2c$RT_detected<110,]

RT1<-lib1[,c(1,3,8)]
RT2<-lib2c[,c(1,3,8)]
RT1<-RT1[RT1$modification_sequence%in%RT2$modification_sequence,]
RT2<-RT2[RT2$modification_sequence%in%RT1$modification_sequence,]
RT1<-RT1[!duplicated(RT1$modification_sequence),2:3]
RT2<-RT2[!duplicated(RT2$modification_sequence),2:3]
colnames(RT2)[1]<-"RT_detected_2"
RT<-join(RT1,RT2)
fit <- coef(lm(RT$RT_detected ~ RT$RT_detected_2))
lib2c$RT_detected<-(lib2c$RT_detected*fit[2])+fit[1]
lib2c$iRT<-lib2c$RT_detected

lib2<-rbind(lib2a,lib2b,lib2c)

dir.create("2corr")
setwd("2corr")

buildSpectraLibPair(lib1, lib2, outputFile = "ExtendedLib_heart.txt", plot = TRUE,
                    clean = FALSE, merge = TRUE, parseAcc = TRUE, consolidateAccession = TRUE, 
                    intensity.cutoff = 5, conf.cutoff = 0.5, nomod = TRUE, nomc = FALSE, enz = "trypsin")

file.rename(from=paste(getwd(),"/ExtendedLib_heart.txt",sep=""), to=paste(workDir,"/ExtendedLib_heart.txt", sep=""))
setwd(workDir)

#3
conf2<-read.xlsx("Newcastle_4__FDR.xlsx", sheet = "Search Summary", colNames = FALSE, rows=33)[1,4]
lib1<-readLibFile("ExtendedLib_heart.txt", clean = FALSE)
lib2<-readLibFile("Newcastle_4.txt", clean = TRUE, intensity.cutoff = 5, conf.cutoff = conf2,
                  nomod = TRUE, nomc = FALSE, enz = "trypsin")
lib2<-canonicalFormat(lib2, format = "peakview")
lib2<-lib2[lib2$uniprot_id!="[ RT-Cal protein ]",]
lib2<-lib2[lib2$uniprot_id!="1/[ RT-Cal protein ]",]
lib2<-lib2[!grepl("NA",rownames(lib2)),]
lib1$iRT<-lib1$RT_detected

dir.create("3")
setwd("3")
checkQuality(lib1,lib2)

#3corr

lib2a<-lib2[lib2$RT_detected<30,]

RT1<-lib1[,c(1,3,8)]
RT2<-lib2a[,c(1,3,8)]
RT1<-RT1[RT1$modification_sequence%in%RT2$modification_sequence,]
RT2<-RT2[RT2$modification_sequence%in%RT1$modification_sequence,]
RT1<-RT1[!duplicated(RT1$modification_sequence),2:3]
RT2<-RT2[!duplicated(RT2$modification_sequence),2:3]
colnames(RT2)[1]<-"RT_detected_2"
RT<-join(RT1,RT2)
fit <- coef(lm(RT$RT_detected ~ RT$RT_detected_2))
lib2a$RT_detected<-(lib2a$RT_detected*fit[2])+fit[1]
lib2a$iRT<-lib2a$RT_detected


lib2b<-lib2[lib2$RT_detected>=30,]
lib2b<-lib2b[lib2b$RT_detected<47,]

RT1<-lib1[,c(1,3,8)]
RT2<-lib2b[,c(1,3,8)]
RT1<-RT1[RT1$modification_sequence%in%RT2$modification_sequence,]
RT2<-RT2[RT2$modification_sequence%in%RT1$modification_sequence,]
RT1<-RT1[!duplicated(RT1$modification_sequence),2:3]
RT2<-RT2[!duplicated(RT2$modification_sequence),2:3]
colnames(RT2)[1]<-"RT_detected_2"
RT<-join(RT1,RT2)
fit <- coef(lm(RT$RT_detected ~ RT$RT_detected_2))
lib2b$RT_detected<-(lib2b$RT_detected*fit[2])+fit[1]
lib2b$iRT<-lib2b$RT_detected

lib2c<-lib2[lib2$RT_detected>=45,]
lib2c<-lib2c[lib2c$RT_detected<80,]

RT1<-lib1[,c(1,3,8)]
RT2<-lib2c[,c(1,3,8)]
RT1<-RT1[RT1$modification_sequence%in%RT2$modification_sequence,]
RT2<-RT2[RT2$modification_sequence%in%RT1$modification_sequence,]
RT1<-RT1[!duplicated(RT1$modification_sequence),2:3]
RT2<-RT2[!duplicated(RT2$modification_sequence),2:3]
colnames(RT2)[1]<-"RT_detected_2"
RT<-join(RT1,RT2)
fit <- coef(lm(RT$RT_detected ~ RT$RT_detected_2))
lib2c$RT_detected<-(lib2c$RT_detected*fit[2])+fit[1]
lib2c$iRT<-lib2c$RT_detected

lib2<-rbind(lib2a,lib2b,lib2c)

dir.create("3corr")
setwd("3corr")

buildSpectraLibPair(lib1, lib2, outputFile = "ExtendedLib_heart.txt", plot = TRUE,
                    clean = FALSE, merge = TRUE, parseAcc = TRUE, consolidateAccession = TRUE, 
                    intensity.cutoff = 5, conf.cutoff = 0.5, nomod = TRUE, nomc = FALSE, enz = "trypsin")

file.rename(from=paste(getwd(),"/ExtendedLib_heart.txt",sep=""), to=paste(workDir,"/ExtendedLib_heart.txt", sep=""))
setwd(workDir)

#4
conf2<-read.xlsx("170116_03_PP_DDA_LV_127__FDR.xlsx", sheet = "Search Summary", colNames = FALSE,
                 rows=33)[1,4]
lib1<-readLibFile("ExtendedLib_heart.txt", clean = FALSE)
lib2<-readLibFile("170116_03_PP_DDA_LV_127.txt", clean = TRUE, intensity.cutoff = 5, conf.cutoff = conf2,
                  nomod = TRUE, nomc = FALSE, enz = "trypsin")
lib2<-canonicalFormat(lib2, format = "peakview")
lib2<-lib2[lib2$uniprot_id!="[ RT-Cal protein ]",]
lib2<-lib2[lib2$uniprot_id!="1/[ RT-Cal protein ]",]
lib2<-lib2[!grepl("NA",rownames(lib2)),]
lib1$iRT<-lib1$RT_detected

dir.create("4")
setwd("4")
checkQuality(lib1,lib2)

#4corr

lib2a<-lib2[lib2$RT_detected<38,]

RT1<-lib1[,c(1,3,8)]
RT2<-lib2a[,c(1,3,8)]
RT1<-RT1[RT1$modification_sequence%in%RT2$modification_sequence,]
RT2<-RT2[RT2$modification_sequence%in%RT1$modification_sequence,]
RT1<-RT1[!duplicated(RT1$modification_sequence),2:3]
RT2<-RT2[!duplicated(RT2$modification_sequence),2:3]
colnames(RT2)[1]<-"RT_detected_2"
RT<-join(RT1,RT2)
fit <- coef(lm(RT$RT_detected ~ RT$RT_detected_2))
lib2a$RT_detected<-(lib2a$RT_detected*fit[2])+fit[1]
lib2a$iRT<-lib2a$RT_detected

lib2b<-lib2[lib2$RT_detected>=38,]
lib2b<-lib2b[lib2b$RT_detected<60,]

RT1<-lib1[,c(1,3,8)]
RT2<-lib2b[,c(1,3,8)]
RT1<-RT1[RT1$modification_sequence%in%RT2$modification_sequence,]
RT2<-RT2[RT2$modification_sequence%in%RT1$modification_sequence,]
RT1<-RT1[!duplicated(RT1$modification_sequence),2:3]
RT2<-RT2[!duplicated(RT2$modification_sequence),2:3]
colnames(RT2)[1]<-"RT_detected_2"
RT<-join(RT1,RT2)
fit <- coef(lm(RT$RT_detected ~ RT$RT_detected_2))
lib2b$RT_detected<-(lib2b$RT_detected*fit[2])+fit[1]
lib2b$iRT<-lib2b$RT_detected

lib2c<-lib2[lib2$RT_detected>=60,]
lib2c<-lib2c[lib2c$RT_detected<100,]

RT1<-lib1[,c(1,3,8)]
RT2<-lib2c[,c(1,3,8)]
RT1<-RT1[RT1$modification_sequence%in%RT2$modification_sequence,]
RT2<-RT2[RT2$modification_sequence%in%RT1$modification_sequence,]
RT1<-RT1[!duplicated(RT1$modification_sequence),2:3]
RT2<-RT2[!duplicated(RT2$modification_sequence),2:3]
colnames(RT2)[1]<-"RT_detected_2"
RT<-join(RT1,RT2)
fit <- coef(lm(RT$RT_detected ~ RT$RT_detected_2))
lib2c$RT_detected<-(lib2c$RT_detected*fit[2])+fit[1]
lib2c$iRT<-lib2c$RT_detected

lib2<-rbind(lib2a,lib2b,lib2c)

dir.create("4corr")
setwd("4corr")

buildSpectraLibPair(lib1, lib2, outputFile = "ExtendedLib_heart.txt", plot = TRUE,
                    clean = FALSE, merge = TRUE, parseAcc = TRUE, consolidateAccession = TRUE, 
                    intensity.cutoff = 5, conf.cutoff = 0.5, nomod = TRUE, nomc = FALSE, enz = "trypsin")

file.rename(from=paste(getwd(),"/ExtendedLib_heart.txt",sep=""), to=paste(workDir,"/ExtendedLib_heart.txt", sep=""))
setwd(workDir)

#5
conf2<-read.xlsx("Worrington research__FDR.xlsx", sheet = "Search Summary", colNames = FALSE,
                 rows=33)[1,4]
lib1<-readLibFile("ExtendedLib_heart.txt", clean = FALSE)
lib2<-readLibFile("Worrington research.txt", clean = TRUE, intensity.cutoff = 5, conf.cutoff = conf2,
                  nomod = TRUE, nomc = FALSE, enz = "trypsin")
lib1$iRT<-lib1$RT_detected
dir.create("5")
setwd("5")

checkQuality(lib1, lib2)

#5corr

lib2a<-lib2[lib2$RT_detected<21,]

RT1<-lib1[,c(1,3,8)]
RT2<-lib2a[,c(1,3,8)]
RT1<-RT1[RT1$modification_sequence%in%RT2$modification_sequence,]
RT2<-RT2[RT2$modification_sequence%in%RT1$modification_sequence,]
RT1<-RT1[!duplicated(RT1$modification_sequence),2:3]
RT2<-RT2[!duplicated(RT2$modification_sequence),2:3]
colnames(RT2)[1]<-"RT_detected_2"
RT<-join(RT1,RT2)
fit <- coef(lm(RT$RT_detected ~ RT$RT_detected_2))
lib2a$RT_detected<-(lib2a$RT_detected*fit[2])+fit[1]
lib2a$iRT<-lib2a$RT_detected


lib2b<-lib2[lib2$RT_detected>=21,]
lib2b<-lib2b[lib2b$RT_detected<38,]

RT1<-lib1[,c(1,3,8)]
RT2<-lib2b[,c(1,3,8)]
RT1<-RT1[RT1$modification_sequence%in%RT2$modification_sequence,]
RT2<-RT2[RT2$modification_sequence%in%RT1$modification_sequence,]
RT1<-RT1[!duplicated(RT1$modification_sequence),2:3]
RT2<-RT2[!duplicated(RT2$modification_sequence),2:3]
colnames(RT2)[1]<-"RT_detected_2"
RT<-join(RT1,RT2)
fit <- coef(lm(RT$RT_detected ~ RT$RT_detected_2))
lib2b$RT_detected<-(lib2b$RT_detected*fit[2])+fit[1]
lib2b$iRT<-lib2b$RT_detected

lib2c<-lib2[lib2$RT_detected>=38,]
lib2c<-lib2c[lib2c$RT_detected<78,]


RT1<-lib1[,c(1,3,8)]
RT2<-lib2c[,c(1,3,8)]
RT1<-RT1[RT1$modification_sequence%in%RT2$modification_sequence,]
RT2<-RT2[RT2$modification_sequence%in%RT1$modification_sequence,]
RT1<-RT1[!duplicated(RT1$modification_sequence),2:3]
RT2<-RT2[!duplicated(RT2$modification_sequence),2:3]
colnames(RT2)[1]<-"RT_detected_2"
RT<-join(RT1,RT2)
fit <- coef(lm(RT$RT_detected ~ RT$RT_detected_2))
lib2c$RT_detected<-(lib2c$RT_detected*fit[2])+fit[1]
lib2c$iRT<-lib2c$RT_detected

lib2d<-lib2[lib2$RT_detected>=78,]
lib2d<-lib2d[lib2d$RT_detected<92,]


RT1<-lib1[,c(1,3,8)]
RT2<-lib2d[,c(1,3,8)]
RT1<-RT1[RT1$modification_sequence%in%RT2$modification_sequence,]
RT2<-RT2[RT2$modification_sequence%in%RT1$modification_sequence,]
RT1<-RT1[!duplicated(RT1$modification_sequence),2:3]
RT2<-RT2[!duplicated(RT2$modification_sequence),2:3]
colnames(RT2)[1]<-"RT_detected_2"
RT<-join(RT1,RT2)
fit <- coef(lm(RT$RT_detected ~ RT$RT_detected_2))
lib2d$RT_detected<-(lib2d$RT_detected*fit[2])+fit[1]
lib2d$iRT<-lib2d$RT_detected

lib2e<-lib2[lib2$RT_detected>=92,]

RT1<-lib1[,c(1,3,8)]
RT2<-lib2e[,c(1,3,8)]
RT1<-RT1[RT1$modification_sequence%in%RT2$modification_sequence,]
RT2<-RT2[RT2$modification_sequence%in%RT1$modification_sequence,]
RT1<-RT1[!duplicated(RT1$modification_sequence),2:3]
RT2<-RT2[!duplicated(RT2$modification_sequence),2:3]
colnames(RT2)[1]<-"RT_detected_2"
RT<-join(RT1,RT2)
fit <- coef(lm(RT$RT_detected ~ RT$RT_detected_2))
lib2e$RT_detected<-(lib2e$RT_detected*fit[2])+fit[1]
lib2e$iRT<-lib2e$RT_detected

lib2<-rbind(lib2a,lib2b,lib2c,lib2d,lib2e)

dir.create("5corr")
setwd("5corr")

buildSpectraLibPair(lib1, lib2, outputFile = "ExtendedLib_heart.txt", plot = TRUE,
                    clean = FALSE, merge = TRUE, parseAcc = TRUE, consolidateAccession = TRUE, 
                    intensity.cutoff = 5, conf.cutoff = 0.5, nomod = TRUE, nomc = FALSE, enz = "trypsin")

file.rename(from=paste(getwd(),"/ExtendedLib_heart.txt",sep=""), 
            to=paste(workDir,"/ExtendedLib_heart.txt", sep=""))
setwd(workDir)

#The final library is now being loaded to remove peptide entries with less then 5 fragment ions
x<-read.delim("ExtendedLib_heart.txt")
y<-as.data.frame(table(x$stripped_sequence))
y<-y[y$Freq>=5,]
x<-x[x$stripped_sequence%in%y$Var1,]
write.table(x,"ExtendedLib_heart.txt",sep="\t")
```
