## Supplementary material for "The generation of a comprehensive spectral library for the analysis of the guinea pig proteome by SWATH-MS": R script 3

```
#Individual search results (.group files) were converted into spectral library format Using PeakView software package with SWATH microApp and saved as .txt

#Guinea pig myometrium spectral library is being assembled here using multiple idividual search results.

#Some searches are more alike than others (due to differences in gradients or column/LC performance). 3 cohorts were identified gruping result files most simmilar to each other. These were first joined into 3 intermediate sublibraries (ExtendedLib_Myometrium.txt, ExtendedLib_1.txt, ExtendedLib_2.txt) which were then merged together in the final to steps.


library(SwathXtend)
library(plyr)
library(openxlsx)

workDir<-getwd()

#1
#Retrieving the conf.cutoff, corresponding to FDR<0.01, from the search summarry in .xlsx format
conf1<-read.xlsx("910__FDR.xlsx", sheet = "Search Summary", colNames = FALSE, rows=33)[1,4]
conf2<-read.xlsx("506__FDR.xlsx", sheet = "Search Summary", colNames = FALSE, rows=33)[1,4]
#Loading and cleaning the library files
lib1<-readLibFile("910.txt", clean = TRUE, intensity.cutoff = 5, conf.cutoff = conf1,
                  nomod = TRUE, nomc = FALSE, enz = "trypsin")
lib2<-readLibFile("506.txt", clean = TRUE, intensity.cutoff = 5, conf.cutoff = conf2,
                  nomod = TRUE, nomc = FALSE, enz = "trypsin")
#correcting the file format (removing the retension time calibrant and nonannotated peptides)
lib1<-canonicalFormat(lib1, format = "peakview")
lib2<-canonicalFormat(lib2, format = "peakview")
lib1<-lib1[lib1$uniprot_id!="[ RT-Cal protein ]",]
lib1<-lib1[lib1$uniprot_id!="1/[ RT-Cal protein ]",]
lib2<-lib2[lib2$uniprot_id!="[ RT-Cal protein ]",]
lib2<-lib2[lib2$uniprot_id!="1/[ RT-Cal protein ]",]
lib1<-lib1[!grepl("NA",rownames(lib1)),]
lib2<-lib2[!grepl("NA",rownames(lib2)),]
#creating a directory for the output files (quality measures for the first step of the library assembly)

dir.create("1")
setwd("1")
# Checking the linearity of RT correlation. If nonlinear, the gradient is divided to linear fragments. 
# QC measures get saved in dedicated folder

# Checking the linearity of RT correlation. If nonlinear, the gradient is divided to linear fragments. 
# QC measures get saved in dedicated folder

checkQuality(lib1,lib2)

#1corr

#based on the visual inspection, the retension time correlation is non-linear
#The External library lib2 (the one that is being added to the "base library" lib1) is therefore manually divided into few, more linear fragments (lib2a-lib2d)

#Linear fragment one

lib2a<-lib2[lib2$RT_detected<19.5,]

RT1<-lib1[,c(1,3,8)]
RT2<-lib2a[,c(1,3,8)]
RT1<-RT1[RT1$modification_sequence%in%RT2$modification_sequence,]
RT2<-RT2[RT2$modification_sequence%in%RT1$modification_sequence,]
RT1<-RT1[!duplicated(RT1$modification_sequence),2:3]
RT2<-RT2[!duplicated(RT2$modification_sequence),2:3]
colnames(RT2)[1]<-"RT_detected_2"
RT<-join(RT1,RT2)

# Linear regression is calculated and the coeficients used to correct the external library RT to mach the base library

fit <- coef(lm(RT$RT_detected ~ RT$RT_detected_2))

lib2a$RT_detected<-(lib2a$RT_detected*fit[2])+fit[1]
lib2a$iRT<-lib2a$RT_detected

#Linear fragment 2

lib2b<-lib2[lib2$RT_detected<26,]
lib2b<-lib2b[lib2b$RT_detected>=19.5,]

RT1<-lib1[,c(1,3,8)]
RT2<-lib2b[,c(1,3,8)]
RT1<-RT1[RT1$modification_sequence%in%RT2$modification_sequence,]
RT2<-RT2[RT2$modification_sequence%in%RT1$modification_sequence,]
RT1<-RT1[!duplicated(RT1$modification_sequence),2:3]
RT2<-RT2[!duplicated(RT2$modification_sequence),2:3]
colnames(RT2)[1]<-"RT_detected_2"
RT<-join(RT1,RT2)
fit <- coef(lm(RT$RT_detected ~ RT$RT_detected_2))
lib2b$RT_detected<-(lib2b$RT_detected*fit[2])+fit[1]
lib2b$iRT<-lib2b$RT_detected

#Linear fragment 3

lib2c<-lib2[lib2$RT_detected>=26,]
lib2c<-lib2c[lib2c$RT_detected<40,]

RT1<-lib1[,c(1,3,8)]
RT2<-lib2c[,c(1,3,8)]
RT1<-RT1[RT1$modification_sequence%in%RT2$modification_sequence,]
RT2<-RT2[RT2$modification_sequence%in%RT1$modification_sequence,]
RT1<-RT1[!duplicated(RT1$modification_sequence),2:3]
RT2<-RT2[!duplicated(RT2$modification_sequence),2:3]
colnames(RT2)[1]<-"RT_detected_2"
RT<-join(RT1,RT2)
fit <- coef(lm(RT$RT_detected ~ RT$RT_detected_2))
lib2c$RT_detected<-(lib2c$RT_detected*fit[2])+fit[1]
lib2c$iRT<-lib2c$RT_detected

#Linear fragment 4

lib2d<-lib2[lib2$RT_detected>=40,]

RT1<-lib1[,c(1,3,8)]
RT2<-lib2d[,c(1,3,8)]
RT1<-RT1[RT1$modification_sequence%in%RT2$modification_sequence,]
RT2<-RT2[RT2$modification_sequence%in%RT1$modification_sequence,]
RT1<-RT1[!duplicated(RT1$modification_sequence),2:3]
RT2<-RT2[!duplicated(RT2$modification_sequence),2:3]
colnames(RT2)[1]<-"RT_detected_2"
RT<-join(RT1,RT2)
fit <- coef(lm(RT$RT_detected ~ RT$RT_detected_2))
lib2d$RT_detected<-(lib2d$RT_detected*fit[2])+fit[1]
lib2d$iRT<-lib2d$RT_detected

#Linear fragments are then reassembled  

lib2<-rbind(lib2a,lib2b,lib2c,lib2d)

#A subdirectory is created to save the output of merging process of the linearized external library with the base library

dir.create("1corr")
setwd("1corr")

#Libraries are merged. QC measures get saved in dedicated subfolder. No cleaning is required at this step since both libraries were cleaned already.


buildSpectraLibPair(lib1, lib2, outputFile = "ExtendedLib_1.txt", plot = TRUE,
                    clean = FALSE, merge = TRUE, parseAcc = TRUE, consolidateAccession = TRUE,
                    nomod=TRUE, conf.cutoff=0.9)

# The merged library is moved to the working directory and will be used as the base library in the following steps.

file.rename(from=paste(getwd(),"/ExtendedLib_1.txt",sep=""), to=paste(workDir,"/ExtendedLib_1.txt", sep=""))
setwd(workDir)

#2

#The process described above is then repeated untill all the partial libraries are joined.
#In the followinf steps, only the external library needs to be cleaned.

conf2<-read.xlsx("308__FDR.xlsx", sheet = "Search Summary", colNames = FALSE, rows=33)[1,4]
lib1<-readLibFile("ExtendedLib_1.txt", clean = FALSE)
lib2<-readLibFile("308.txt", clean = TRUE, intensity.cutoff = 5, conf.cutoff = conf2,
                  nomod = TRUE, nomc = FALSE, enz = "trypsin")
lib2<-canonicalFormat(lib2, format = "peakview")
lib2<-lib2[lib2$uniprot_id!="[ RT-Cal protein ]",]
lib2<-lib2[lib2$uniprot_id!="1/[ RT-Cal protein ]",]
lib2<-lib2[!grepl("NA",rownames(lib2)),]
lib1$RT_detected<-lib1$iRT

dir.create("2")
setwd("2")
checkQuality(lib1,lib2)

#2corr

lib2a<-lib2[lib2$RT_detected<22,]

RT1<-lib1[,c(1,3,8)]
RT2<-lib2a[,c(1,3,8)]
RT1<-RT1[RT1$modification_sequence%in%RT2$modification_sequence,]
RT2<-RT2[RT2$modification_sequence%in%RT1$modification_sequence,]
RT1<-RT1[!duplicated(RT1$modification_sequence),2:3]
RT2<-RT2[!duplicated(RT2$modification_sequence),2:3]
colnames(RT2)[1]<-"RT_detected_2"
RT<-join(RT1,RT2)
fit <- coef(lm(RT$RT_detected ~ RT$RT_detected_2))
lib2a$RT_detected<-(lib2a$RT_detected*fit[2])+fit[1]
lib2a$iRT<-lib2a$RT_detected

lib2b<-lib2[lib2$RT_detected>=22,]
lib2b<-lib2b[lib2b$RT_detected<35,]

RT1<-lib1[,c(1,3,8)]
RT2<-lib2b[,c(1,3,8)]
RT1<-RT1[RT1$modification_sequence%in%RT2$modification_sequence,]
RT2<-RT2[RT2$modification_sequence%in%RT1$modification_sequence,]
RT1<-RT1[!duplicated(RT1$modification_sequence),2:3]
RT2<-RT2[!duplicated(RT2$modification_sequence),2:3]
colnames(RT2)[1]<-"RT_detected_2"
RT<-join(RT1,RT2)
fit <- coef(lm(RT$RT_detected ~ RT$RT_detected_2))
lib2b$RT_detected<-(lib2b$RT_detected*fit[2])+fit[1]
lib2b$iRT<-lib2b$RT_detected

lib2c<-lib2[lib2$RT_detected>=35,]
lib2c<-lib2c[lib2c$RT_detected<55,]

RT1<-lib1[,c(1,3,8)]
RT2<-lib2c[,c(1,3,8)]
RT1<-RT1[RT1$modification_sequence%in%RT2$modification_sequence,]
RT2<-RT2[RT2$modification_sequence%in%RT1$modification_sequence,]
RT1<-RT1[!duplicated(RT1$modification_sequence),2:3]
RT2<-RT2[!duplicated(RT2$modification_sequence),2:3]
colnames(RT2)[1]<-"RT_detected_2"
RT<-join(RT1,RT2)
fit <- coef(lm(RT$RT_detected ~ RT$RT_detected_2))
lib2c$RT_detected<-(lib2c$RT_detected*fit[2])+fit[1]
lib2c$iRT<-lib2c$RT_detected

lib2d<-lib2[lib2$RT_detected>=55,]

RT1<-lib1[,c(1,3,8)]
RT2<-lib2d[,c(1,3,8)]
RT1<-RT1[RT1$modification_sequence%in%RT2$modification_sequence,]
RT2<-RT2[RT2$modification_sequence%in%RT1$modification_sequence,]
RT1<-RT1[!duplicated(RT1$modification_sequence),2:3]
RT2<-RT2[!duplicated(RT2$modification_sequence),2:3]
colnames(RT2)[1]<-"RT_detected_2"
RT<-join(RT1,RT2)
fit <- coef(lm(RT$RT_detected ~ RT$RT_detected_2))
lib2d$RT_detected<-(lib2d$RT_detected*fit[2])+fit[1]
lib2d$iRT<-lib2d$RT_detected

lib2<-rbind(lib2a,lib2b,lib2c,lib2d)

dir.create("2corr")
setwd("2corr")

buildSpectraLibPair(lib1, lib2, outputFile = "ExtendedLib_1.txt", plot = TRUE,
                    clean = FALSE, merge = TRUE, parseAcc = TRUE, consolidateAccession = TRUE,
                    nomod=TRUE, conf.cutoff=0.9)

file.rename(from=paste(getwd(),"/ExtendedLib_1.txt",sep=""), to=paste(workDir,"/ExtendedLib_1.txt", sep=""))
setwd(workDir)

#3

conf2<-read.xlsx("13-189-210__FDR.xlsx", sheet = "Search Summary", colNames = FALSE, rows=33)[1,4]
lib1<-readLibFile("ExtendedLib_1.txt", clean = FALSE)
lib2<-readLibFile("13-189-210.txt", clean = TRUE, intensity.cutoff = 5, conf.cutoff = conf2,
                  nomod = TRUE, nomc = FALSE, enz = "trypsin")
lib2<-canonicalFormat(lib2, format = "peakview")
lib2<-lib2[lib2$uniprot_id!="[ RT-Cal protein ]",]
lib2<-lib2[lib2$uniprot_id!="1/[ RT-Cal protein ]",]
lib2<-lib2[!grepl("NA",rownames(lib2)),]
lib1$RT_detected<-lib1$iRT

dir.create("3")
setwd("3")
checkQuality(lib1,lib2)


#3corr

RT1<-lib1[,c(1,3,8)]
RT2<-lib2[,c(1,3,8)]
RT1<-RT1[RT1$modification_sequence%in%RT2$modification_sequence,]
RT2<-RT2[RT2$modification_sequence%in%RT1$modification_sequence,]
RT1<-RT1[!duplicated(RT1$modification_sequence),2:3]
RT2<-RT2[!duplicated(RT2$modification_sequence),2:3]
colnames(RT2)[1]<-"RT_detected_2"
RT<-join(RT1,RT2)
fit <- coef(lm(RT$RT_detected ~ RT$RT_detected_2))
lib2$RT_detected<-(lib2$RT_detected*fit[2])+fit[1]
lib2$iRT<-lib2$RT_detected

dir.create("3corr")
setwd("3corr")

buildSpectraLibPair(lib1, lib2, outputFile = "ExtendedLib_1.txt", plot = TRUE,
                    clean = FALSE, merge = TRUE, parseAcc = TRUE, consolidateAccession = TRUE,
                    nomod=TRUE, conf.cutoff=0.9)

file.rename(from=paste(getwd(),"/ExtendedLib_1.txt",sep=""), to=paste(workDir,"/ExtendedLib_1.txt", sep=""))
setwd(workDir)

#4

conf2<-read.xlsx("408_150315120153__FDR.xlsx", sheet = "Search Summary", colNames = FALSE, rows=33)[1,4]
lib1<-readLibFile("ExtendedLib_1.txt",clean=FALSE)
lib2<-readLibFile("408_150315120153.txt", clean = TRUE, intensity.cutoff = 5, conf.cutoff = conf2,
                  nomod = TRUE, nomc = FALSE, enz = "trypsin")
lib2<-canonicalFormat(lib2, format = "peakview")
lib2<-lib2[lib2$uniprot_id!="[ RT-Cal protein ]",]
lib2<-lib2[lib2$uniprot_id!="1/[ RT-Cal protein ]",]
lib2<-lib2[!grepl("NA",rownames(lib2)),]
lib1$RT_detected<-lib1$iRT


dir.create("4")
setwd("4")
checkQuality(lib1,lib2)

#4corr

lib2a<-lib2[lib2$RT_detected<25,]

RT1<-lib1[,c(1,3,8)]
RT2<-lib2a[,c(1,3,8)]
RT1<-RT1[RT1$modification_sequence%in%RT2$modification_sequence,]
RT2<-RT2[RT2$modification_sequence%in%RT1$modification_sequence,]
RT1<-RT1[!duplicated(RT1$modification_sequence),2:3]
RT2<-RT2[!duplicated(RT2$modification_sequence),2:3]
colnames(RT2)[1]<-"RT_detected_2"
RT<-join(RT1,RT2)
fit <- coef(lm(RT$RT_detected ~ RT$RT_detected_2))
lib2a$RT_detected<-(lib2a$RT_detected*fit[2])+fit[1]
lib2a$iRT<-lib2a$RT_detected

lib2b<-lib2[lib2$RT_detected>=25,]
lib2b<-lib2b[lib2b$RT_detected<65,]

RT1<-lib1[,c(1,3,8)]
RT2<-lib2b[,c(1,3,8)]
RT1<-RT1[RT1$modification_sequence%in%RT2$modification_sequence,]
RT2<-RT2[RT2$modification_sequence%in%RT1$modification_sequence,]
RT1<-RT1[!duplicated(RT1$modification_sequence),2:3]
RT2<-RT2[!duplicated(RT2$modification_sequence),2:3]
colnames(RT2)[1]<-"RT_detected_2"
RT<-join(RT1,RT2)
fit <- coef(lm(RT$RT_detected ~ RT$RT_detected_2))
lib2b$RT_detected<-(lib2b$RT_detected*fit[2])+fit[1]
lib2b$iRT<-lib2b$RT_detected

lib2c<-lib2[lib2$RT_detected>=65,]

RT1<-lib1[,c(1,3,8)]
RT2<-lib2c[,c(1,3,8)]
RT1<-RT1[RT1$modification_sequence%in%RT2$modification_sequence,]
RT2<-RT2[RT2$modification_sequence%in%RT1$modification_sequence,]
RT1<-RT1[!duplicated(RT1$modification_sequence),2:3]
RT2<-RT2[!duplicated(RT2$modification_sequence),2:3]
colnames(RT2)[1]<-"RT_detected_2"
RT<-join(RT1,RT2)
fit <- coef(lm(RT$RT_detected ~ RT$RT_detected_2))
lib2c$RT_detected<-(lib2c$RT_detected*fit[2])+fit[1]
lib2c$iRT<-lib2c$RT_detected

lib2<-rbind(lib2a,lib2b,lib2c)

dir.create("4corr")
setwd("4corr")

buildSpectraLibPair(lib1, lib2, outputFile = "ExtendedLib_1.txt", plot = TRUE,
                    clean = FALSE, merge = TRUE, parseAcc = TRUE, consolidateAccession = TRUE,
                    nomod=TRUE, conf.cutoff=0.9)

file.rename(from=paste(getwd(),"/ExtendedLib_1.txt",sep=""), to=paste(workDir,"/ExtendedLib_1.txt", sep=""))
setwd(workDir)

#5

conf2<-read.xlsx("408_150314024101__FDR.xlsx", sheet = "Search Summary", colNames = FALSE, rows=33)[1,4]
lib1<-readLibFile("ExtendedLib_1.txt",clean=FALSE)
lib2<-readLibFile("408_150314024101.txt", clean = TRUE, intensity.cutoff = 5, conf.cutoff = conf2,
                  nomod = TRUE, nomc = FALSE, enz = "trypsin")
lib2<-canonicalFormat(lib2, format = "peakview")
lib2<-lib2[lib2$uniprot_id!="[ RT-Cal protein ]",]
lib2<-lib2[lib2$uniprot_id!="1/[ RT-Cal protein ]",]
lib2<-lib2[!grepl("NA",rownames(lib2)),]
lib1$RT_detected<-lib1$iRT

dir.create("5")
setwd("5")
checkQuality(lib1,lib2)

#5corr

lib2a<-lib2[lib2$RT_detected<25,]
lib2a<-lib2a[lib2a$RT_detected>=20,]

RT1<-lib1[,c(1,3,8)]
RT2<-lib2a[,c(1,3,8)]
RT1<-RT1[RT1$modification_sequence%in%RT2$modification_sequence,]
RT2<-RT2[RT2$modification_sequence%in%RT1$modification_sequence,]
RT1<-RT1[!duplicated(RT1$modification_sequence),2:3]
RT2<-RT2[!duplicated(RT2$modification_sequence),2:3]
colnames(RT2)[1]<-"RT_detected_2"
RT<-join(RT1,RT2)
fit <- coef(lm(RT$RT_detected ~ RT$RT_detected_2))
lib2a$RT_detected<-(lib2a$RT_detected*fit[2])+fit[1]
lib2a$iRT<-lib2a$RT_detected

lib2b<-lib2[lib2$RT_detected>=25,]
lib2b<-lib2b[lib2b$RT_detected<40,]

RT1<-lib1[,c(1,3,8)]
RT2<-lib2b[,c(1,3,8)]
RT1<-RT1[RT1$modification_sequence%in%RT2$modification_sequence,]
RT2<-RT2[RT2$modification_sequence%in%RT1$modification_sequence,]
RT1<-RT1[!duplicated(RT1$modification_sequence),2:3]
RT2<-RT2[!duplicated(RT2$modification_sequence),2:3]
colnames(RT2)[1]<-"RT_detected_2"
RT<-join(RT1,RT2)
fit <- coef(lm(RT$RT_detected ~ RT$RT_detected_2))
lib2b$RT_detected<-(lib2b$RT_detected*fit[2])+fit[1]
lib2b$iRT<-lib2b$RT_detected

lib2c<-lib2[lib2$RT_detected>=40,]

RT1<-lib1[,c(1,3,8)]
RT2<-lib2c[,c(1,3,8)]
RT1<-RT1[RT1$modification_sequence%in%RT2$modification_sequence,]
RT2<-RT2[RT2$modification_sequence%in%RT1$modification_sequence,]
RT1<-RT1[!duplicated(RT1$modification_sequence),2:3]
RT2<-RT2[!duplicated(RT2$modification_sequence),2:3]
colnames(RT2)[1]<-"RT_detected_2"
RT<-join(RT1,RT2)
fit <- coef(lm(RT$RT_detected ~ RT$RT_detected_2))
lib2c$RT_detected<-(lib2c$RT_detected*fit[2])+fit[1]
lib2c$iRT<-lib2c$RT_detected

lib2<-rbind(lib2a,lib2b,lib2c)

dir.create("5corr")
setwd("5corr")

buildSpectraLibPair(lib1, lib2, outputFile = "ExtendedLib_1.txt", plot = TRUE,
                    clean = FALSE, merge = TRUE, parseAcc = TRUE, consolidateAccession = TRUE,
                    nomod=TRUE, conf.cutoff=0.9)

file.rename(from=paste(getwd(),"/ExtendedLib_1.txt",sep=""), to=paste(workDir,"/ExtendedLib_1.txt", sep=""))
setwd(workDir)

#6

conf2<-read.xlsx("20151030_147NP_DDA_08__FDR.xlsx", sheet = "Search Summary", colNames = FALSE,
                 rows=33)[1,4]
lib1<-readLibFile("ExtendedLib_1.txt",clean=FALSE)
lib2<-readLibFile("20151030_147NP_DDA_08.txt", clean = TRUE, intensity.cutoff = 5, conf.cutoff = conf2,
                  nomod = TRUE, nomc = FALSE, enz = "trypsin")
lib2<-canonicalFormat(lib2, format = "peakview")
lib2<-lib2[lib2$uniprot_id!="[ RT-Cal protein ]",]
lib2<-lib2[lib2$uniprot_id!="1/[ RT-Cal protein ]",]
lib2<-lib2[!grepl("NA",rownames(lib2)),]
lib1$RT_detected<-lib1$iRT

dir.create("6")
setwd("6")
checkQuality(lib1,lib2)

#6corr

lib2a<-lib2[lib2$RT_detected<24,]

RT1<-lib1[,c(1,3,8)]
RT2<-lib2a[,c(1,3,8)]
RT1<-RT1[RT1$modification_sequence%in%RT2$modification_sequence,]
RT2<-RT2[RT2$modification_sequence%in%RT1$modification_sequence,]
RT1<-RT1[!duplicated(RT1$modification_sequence),2:3]
RT2<-RT2[!duplicated(RT2$modification_sequence),2:3]
colnames(RT2)[1]<-"RT_detected_2"
RT<-join(RT1,RT2)
fit <- coef(lm(RT$RT_detected ~ RT$RT_detected_2))
lib2a$RT_detected<-(lib2a$RT_detected*fit[2])+fit[1]
lib2a$iRT<-lib2a$RT_detected

lib2b<-lib2[lib2$RT_detected>=24,]
lib2b<-lib2b[lib2b$RT_detected<145,]

RT1<-lib1[,c(1,3,8)]
RT2<-lib2b[,c(1,3,8)]
RT1<-RT1[RT1$modification_sequence%in%RT2$modification_sequence,]
RT2<-RT2[RT2$modification_sequence%in%RT1$modification_sequence,]
RT1<-RT1[!duplicated(RT1$modification_sequence),2:3]
RT2<-RT2[!duplicated(RT2$modification_sequence),2:3]
colnames(RT2)[1]<-"RT_detected_2"
RT<-join(RT1,RT2)
fit <- coef(lm(RT$RT_detected ~ RT$RT_detected_2))
lib2b$RT_detected<-(lib2b$RT_detected*fit[2])+fit[1]
lib2b$iRT<-lib2b$RT_detected

lib2<-rbind(lib2a,lib2b)

dir.create("6corr")
setwd("6corr")

buildSpectraLibPair(lib1, lib2, outputFile = "ExtendedLib_1.txt", plot = TRUE,
                    clean = FALSE, merge = TRUE, parseAcc = TRUE, consolidateAccession = TRUE,
                    nomod=TRUE, conf.cutoff=0.9)

file.rename(from=paste(getwd(),"/ExtendedLib_1.txt",sep=""), to=paste(workDir,"/ExtendedLib_1.txt", sep=""))
setwd(workDir)

#7

conf2<-read.xlsx("20151030_147NP_DDA_08_151201053308__FDR.xlsx", sheet = "Search Summary", 
                 colNames = FALSE, rows=33)[1,4]
lib1<-readLibFile("ExtendedLib_1.txt",clean=FALSE)
lib2<-readLibFile("20151030_147NP_DDA_08_151201053308.txt",clean = TRUE, intensity.cutoff = 5, 
                  conf.cutoff = conf2, nomod = TRUE, nomc = FALSE, enz = "trypsin")
lib2<-canonicalFormat(lib2, format = "peakview")
lib2<-lib2[lib2$uniprot_id!="[ RT-Cal protein ]",]
lib2<-lib2[lib2$uniprot_id!="1/[ RT-Cal protein ]",]
lib2<-lib2[!grepl("NA",rownames(lib2)),]
lib1$RT_detected<-lib1$iRT

dir.create("7")
setwd("7")
checkQuality(lib1,lib2)

#7corr

lib2a<-lib2[lib2$RT_detected>=55,]
lib2a<-lib2a[lib2a$RT_detected<145,]

RT1<-lib1[,c(1,3,8)]
RT2<-lib2a[,c(1,3,8)]
RT1<-RT1[RT1$modification_sequence%in%RT2$modification_sequence,]
RT2<-RT2[RT2$modification_sequence%in%RT1$modification_sequence,]
RT1<-RT1[!duplicated(RT1$modification_sequence),2:3]
RT2<-RT2[!duplicated(RT2$modification_sequence),2:3]
colnames(RT2)[1]<-"RT_detected_2"
RT<-join(RT1,RT2)
fit <- coef(lm(RT$RT_detected ~ RT$RT_detected_2))
lib2a$RT_detected<-(lib2a$RT_detected*fit[2])+fit[1]
lib2a$iRT<-lib2a$RT_detected


lib2b<-lib2[lib2$RT_detected<55,]
lib2b<-lib2b[lib2b$RT_detected>=35,]

RT1<-lib1[,c(1,3,8)]
RT2<-lib2b[,c(1,3,8)]
RT1<-RT1[RT1$modification_sequence%in%RT2$modification_sequence,]
RT2<-RT2[RT2$modification_sequence%in%RT1$modification_sequence,]
RT1<-RT1[!duplicated(RT1$modification_sequence),2:3]
RT2<-RT2[!duplicated(RT2$modification_sequence),2:3]
colnames(RT2)[1]<-"RT_detected_2"
RT<-join(RT1,RT2)
fit <- coef(lm(RT$RT_detected ~ RT$RT_detected_2))
lib2b$RT_detected<-(lib2b$RT_detected*fit[2])+fit[1]
lib2b$iRT<-lib2b$RT_detected

lib2c<-lib2[lib2$RT_detected<35,]

RT1<-lib1[,c(1,3,8)]
RT2<-lib2c[,c(1,3,8)]
RT1<-RT1[RT1$modification_sequence%in%RT2$modification_sequence,]
RT2<-RT2[RT2$modification_sequence%in%RT1$modification_sequence,]
RT1<-RT1[!duplicated(RT1$modification_sequence),2:3]
RT2<-RT2[!duplicated(RT2$modification_sequence),2:3]
colnames(RT2)[1]<-"RT_detected_2"
RT<-join(RT1,RT2)
fit <- coef(lm(RT$RT_detected ~ RT$RT_detected_2))
lib2c$RT_detected<-(lib2c$RT_detected*fit[2])+fit[1]
lib2c$iRT<-lib2c$RT_detected

lib2<-rbind(lib2a,lib2b,lib2c)

dir.create("7corr")
setwd("7corr")

buildSpectraLibPair(lib1, lib2, outputFile = "ExtendedLib_1.txt", plot = TRUE,
                    clean = FALSE, merge = TRUE, parseAcc = TRUE, consolidateAccession = TRUE,
                    nomod=TRUE, conf.cutoff=0.9)

file.rename(from=paste(getwd(),"/ExtendedLib_1.txt",sep=""), to=paste(workDir,"/ExtendedLib_1.txt", sep=""))
setwd(workDir)

#8


conf1<-read.xlsx("170228_14_PP_GP_Myometrium__FDR.xlsx", sheet = "Search Summary", colNames = FALSE,
                 rows=33)[1,4]
conf2<-read.xlsx("170306_14_PP_GP_Myometrium__FDR.xlsx", sheet = "Search Summary", colNames = FALSE,
                 rows=33)[1,4]
lib1<-readLibFile("170228_14_PP_GP_Myometrium.txt", clean = TRUE, intensity.cutoff = 5, 
                  conf.cutoff = conf1, nomod = TRUE, nomc = FALSE, enz = "trypsin")
lib2<-readLibFile("170306_14_PP_GP_Myometrium.txt", clean = TRUE, intensity.cutoff = 5, conf.cutoff = conf2,
                  nomod = TRUE, nomc = FALSE, enz = "trypsin")
lib1<-canonicalFormat(lib1, format = "peakview")
lib2<-canonicalFormat(lib2, format = "peakview")
lib1<-lib1[lib1$uniprot_id!="[ RT-Cal protein ]",]
lib1<-lib1[lib1$uniprot_id!="1/[ RT-Cal protein ]",]
lib2<-lib2[lib2$uniprot_id!="[ RT-Cal protein ]",]
lib2<-lib2[lib2$uniprot_id!="1/[ RT-Cal protein ]",]
lib1<-lib1[!grepl("NA",rownames(lib1)),]
lib2<-lib2[!grepl("NA",rownames(lib2)),]


dir.create("8")
setwd("8")
checkQuality(lib1,lib2)

#8corr

RT1<-lib1[,c(1,3,8)]
RT2<-lib2[,c(1,3,8)]
RT1<-RT1[RT1$modification_sequence%in%RT2$modification_sequence,]
RT2<-RT2[RT2$modification_sequence%in%RT1$modification_sequence,]
RT1<-RT1[!duplicated(RT1$modification_sequence),2:3]
RT2<-RT2[!duplicated(RT2$modification_sequence),2:3]
colnames(RT2)[1]<-"RT_detected_2"
RT<-join(RT1,RT2)
fit <- coef(lm(RT$RT_detected ~ RT$RT_detected_2))
lib2$RT_detected<-(lib2$RT_detected*fit[2])+fit[1]
lib2$iRT<-lib2$RT_detected


dir.create("8corr")
setwd("8corr")

buildSpectraLibPair(lib1, lib2, outputFile = "ExtendedLib_Myometrium.txt", plot = TRUE,
                    clean = FALSE, merge = TRUE, parseAcc = TRUE, consolidateAccession = TRUE,
                    nomod=TRUE, conf.cutoff=0.9)

file.rename(from=paste(getwd(),"/ExtendedLib_Myometrium.txt",sep=""), 
            to=paste(workDir,"/ExtendedLib_Myometrium.txt", sep=""))
setwd(workDir)

#9

conf2<-read.xlsx("170310_14_PP_GP_Myometrium__FDR.xlsx", sheet = "Search Summary", colNames = FALSE,
                 rows=33)[1,4]
lib1<-readLibFile("ExtendedLib_Myometrium.txt")
lib2<-readLibFile("170310_14_PP_GP_Myometrium.txt", clean = TRUE, intensity.cutoff = 5, conf.cutoff = conf2,
                  nomod = TRUE, nomc = FALSE, enz = "trypsin")
lib1<-canonicalFormat(lib1, format = "peakview")
lib2<-canonicalFormat(lib2, format = "peakview")
lib2<-lib2[lib2$uniprot_id!="[ RT-Cal protein ]",]
lib2<-lib2[lib2$uniprot_id!="1/[ RT-Cal protein ]",]
lib2<-lib2[!grepl("NA",rownames(lib2)),]

lib1$RT_detected<-lib1$iRT

dir.create("9")
setwd("9")
checkQuality(lib1,lib2)

#9corr

lib2a<-lib2[lib2$RT_detected<70,]

RT1<-lib1[,c(1,3,8)]
RT2<-lib2a[,c(1,3,8)]
RT1<-RT1[RT1$modification_sequence%in%RT2$modification_sequence,]
RT2<-RT2[RT2$modification_sequence%in%RT1$modification_sequence,]
RT1<-RT1[!duplicated(RT1$modification_sequence),2:3]
RT2<-RT2[!duplicated(RT2$modification_sequence),2:3]
colnames(RT2)[1]<-"RT_detected_2"
RT<-join(RT1,RT2)
fit <- coef(lm(RT$RT_detected ~ RT$RT_detected_2))
lib2a$RT_detected<-(lib2a$RT_detected*fit[2])+fit[1]
lib2a$iRT<-lib2a$RT_detected


lib2b<-lib2[lib2$RT_detected>=70,]
lib2b<-lib2b[lib2b$RT_detected<100,]

RT1<-lib1[,c(1,3,8)]
RT2<-lib2b[,c(1,3,8)]
RT1<-RT1[RT1$modification_sequence%in%RT2$modification_sequence,]
RT2<-RT2[RT2$modification_sequence%in%RT1$modification_sequence,]
RT1<-RT1[!duplicated(RT1$modification_sequence),2:3]
RT2<-RT2[!duplicated(RT2$modification_sequence),2:3]
colnames(RT2)[1]<-"RT_detected_2"
RT<-join(RT1,RT2)
fit <- coef(lm(RT$RT_detected ~ RT$RT_detected_2))
lib2b$RT_detected<-(lib2b$RT_detected*fit[2])+fit[1]
lib2b$iRT<-lib2b$RT_detected

lib2c<-lib2[lib2$RT_detected>=100,]

RT1<-lib1[,c(1,3,8)]
RT2<-lib2c[,c(1,3,8)]
RT1<-RT1[RT1$modification_sequence%in%RT2$modification_sequence,]
RT2<-RT2[RT2$modification_sequence%in%RT1$modification_sequence,]
RT1<-RT1[!duplicated(RT1$modification_sequence),2:3]
RT2<-RT2[!duplicated(RT2$modification_sequence),2:3]
colnames(RT2)[1]<-"RT_detected_2"
RT<-join(RT1,RT2)
fit <- coef(lm(RT$RT_detected ~ RT$RT_detected_2))
lib2c$RT_detected<-(lib2c$RT_detected*fit[2])+fit[1]
lib2c$iRT<-lib2c$RT_detected


lib2<-rbind(lib2a,lib2b,lib2c)

dir.create("9corr")
setwd("9corr")

buildSpectraLibPair(lib1, lib2, outputFile = "ExtendedLib_Myometrium.txt", plot = TRUE,
                    clean = FALSE, merge = TRUE, parseAcc = TRUE, consolidateAccession = TRUE,
                    nomod=TRUE, conf.cutoff=0.9)

file.rename(from=paste(getwd(),"/ExtendedLib_Myometrium.txt",sep=""), 
            to=paste(workDir,"/ExtendedLib_Myometrium.txt", sep=""))
setwd(workDir)

#10

conf2<-read.xlsx("170324_19_PP_GPlibrary_Myometrium__FDR.xlsx", sheet = "Search Summary",
                 colNames = FALSE, rows=33)[1,4]
lib1<-readLibFile("ExtendedLib_Myometrium.txt",clean=FALSE)
lib2<-readLibFile("170324_19_PP_GPlibrary_Myometrium.txt",clean = TRUE, intensity.cutoff = 5, 
                  conf.cutoff = conf2, nomod = TRUE, nomc = FALSE, enz = "trypsin")
lib2<-canonicalFormat(lib2, format = "peakview")
lib2<-lib2[lib2$uniprot_id!="[ RT-Cal protein ]",]
lib2<-lib2[lib2$uniprot_id!="1/[ RT-Cal protein ]",]
lib2<-lib2[!grepl("NA",rownames(lib2)),]
lib1$RT_detected<-lib1$iRT

dir.create("10")
setwd("10")
checkQuality(lib1,lib2)

#10corr

lib2a<-lib2[lib2$RT_detected<60,]

RT1<-lib1[,c(1,3,8)]
RT2<-lib2a[,c(1,3,8)]
RT1<-RT1[RT1$modification_sequence%in%RT2$modification_sequence,]
RT2<-RT2[RT2$modification_sequence%in%RT1$modification_sequence,]
RT1<-RT1[!duplicated(RT1$modification_sequence),2:3]
RT2<-RT2[!duplicated(RT2$modification_sequence),2:3]
colnames(RT2)[1]<-"RT_detected_2"
RT<-join(RT1,RT2)
fit <- coef(lm(RT$RT_detected ~ RT$RT_detected_2))
lib2a$RT_detected<-(lib2a$RT_detected*fit[2])+fit[1]
lib2a$iRT<-lib2a$RT_detected


lib2b<-lib2[lib2$RT_detected>=60,]
lib2b<-lib2b[lib2b$RT_detected<90,]

RT1<-lib1[,c(1,3,8)]
RT2<-lib2b[,c(1,3,8)]
RT1<-RT1[RT1$modification_sequence%in%RT2$modification_sequence,]
RT2<-RT2[RT2$modification_sequence%in%RT1$modification_sequence,]
RT1<-RT1[!duplicated(RT1$modification_sequence),2:3]
RT2<-RT2[!duplicated(RT2$modification_sequence),2:3]
colnames(RT2)[1]<-"RT_detected_2"
RT<-join(RT1,RT2)
fit <- coef(lm(RT$RT_detected ~ RT$RT_detected_2))
lib2b$RT_detected<-(lib2b$RT_detected*fit[2])+fit[1]
lib2b$iRT<-lib2b$RT_detected

lib2c<-lib2[lib2$RT_detected>=90,]

RT1<-lib1[,c(1,3,8)]
RT2<-lib2c[,c(1,3,8)]
RT1<-RT1[RT1$modification_sequence%in%RT2$modification_sequence,]
RT2<-RT2[RT2$modification_sequence%in%RT1$modification_sequence,]
RT1<-RT1[!duplicated(RT1$modification_sequence),2:3]
RT2<-RT2[!duplicated(RT2$modification_sequence),2:3]
colnames(RT2)[1]<-"RT_detected_2"
RT<-join(RT1,RT2)
fit <- coef(lm(RT$RT_detected ~ RT$RT_detected_2))
lib2c$RT_detected<-(lib2c$RT_detected*fit[2])+fit[1]
lib2c$iRT<-lib2c$RT_detected

lib2<-rbind(lib2a,lib2b,lib2c)

dir.create("10corr")
setwd("10corr")

buildSpectraLibPair(lib1, lib2, outputFile = "ExtendedLib_Myometrium.txt", plot = TRUE,
                    clean = FALSE, merge = TRUE, parseAcc = TRUE, consolidateAccession = TRUE,
                    nomod=TRUE, conf.cutoff=0.9)

file.rename(from=paste(getwd(),"/ExtendedLib_Myometrium.txt",sep=""), 
            to=paste(workDir,"/ExtendedLib_Myometrium.txt", sep=""))
setwd(workDir)

#11

conf1<-read.xlsx("160909_01_NP_147_DDA__FDR.xlsx", sheet = "Search Summary", colNames = FALSE, rows=33)[1,4]
conf2<-read.xlsx("160907_09_GPmyo_233_DDA__FDR.xlsx", sheet = "Search Summary", colNames = FALSE,
                 rows=33)[1,4]
lib1<-readLibFile("160909_01_NP_147_DDA.txt", clean = TRUE, intensity.cutoff = 5, conf.cutoff = conf1,
                  nomod = TRUE, nomc = FALSE, enz = "trypsin")
lib2<-readLibFile("160907_09_GPmyo_233_DDA.txt", clean = TRUE, intensity.cutoff = 5, conf.cutoff = conf2,
                  nomod = TRUE, nomc = FALSE, enz = "trypsin")
lib2<-canonicalFormat(lib2, format = "peakview")
lib2<-lib2[lib2$uniprot_id!="[ RT-Cal protein ]",]
lib2<-lib2[lib2$uniprot_id!="1/[ RT-Cal protein ]",]
lib2<-lib2[!grepl("NA",rownames(lib2)),]
lib1$RT_detected<-lib1$iRT

dir.create("11")
setwd("11")
checkQuality(lib1,lib2)

#11corr

lib2a<-lib2[lib2$RT_detected<25,]

RT1<-lib1[,c(1,3,8)]
RT2<-lib2a[,c(1,3,8)]
RT1<-RT1[RT1$modification_sequence%in%RT2$modification_sequence,]
RT2<-RT2[RT2$modification_sequence%in%RT1$modification_sequence,]
RT1<-RT1[!duplicated(RT1$modification_sequence),2:3]
RT2<-RT2[!duplicated(RT2$modification_sequence),2:3]
colnames(RT2)[1]<-"RT_detected_2"
RT<-join(RT1,RT2)
fit <- coef(lm(RT$RT_detected ~ RT$RT_detected_2))
lib2a$RT_detected<-(lib2a$RT_detected*fit[2])+fit[1]
lib2a$iRT<-lib2a$RT_detected


lib2b<-lib2[lib2$RT_detected>=25,]
lib2b<-lib2b[lib2b$RT_detected<40,]

RT1<-lib1[,c(1,3,8)]
RT2<-lib2b[,c(1,3,8)]
RT1<-RT1[RT1$modification_sequence%in%RT2$modification_sequence,]
RT2<-RT2[RT2$modification_sequence%in%RT1$modification_sequence,]
RT1<-RT1[!duplicated(RT1$modification_sequence),2:3]
RT2<-RT2[!duplicated(RT2$modification_sequence),2:3]
colnames(RT2)[1]<-"RT_detected_2"
RT<-join(RT1,RT2)
fit <- coef(lm(RT$RT_detected ~ RT$RT_detected_2))
lib2b$RT_detected<-(lib2b$RT_detected*fit[2])+fit[1]
lib2b$iRT<-lib2b$RT_detected

lib2c<-lib2[lib2$RT_detected>=40,]
lib2c<-lib2c[lib2c$RT_detected<80,]

RT1<-lib1[,c(1,3,8)]
RT2<-lib2c[,c(1,3,8)]
RT1<-RT1[RT1$modification_sequence%in%RT2$modification_sequence,]
RT2<-RT2[RT2$modification_sequence%in%RT1$modification_sequence,]
RT1<-RT1[!duplicated(RT1$modification_sequence),2:3]
RT2<-RT2[!duplicated(RT2$modification_sequence),2:3]
colnames(RT2)[1]<-"RT_detected_2"
RT<-join(RT1,RT2)
fit <- coef(lm(RT$RT_detected ~ RT$RT_detected_2))
lib2c$RT_detected<-(lib2c$RT_detected*fit[2])+fit[1]
lib2c$iRT<-lib2c$RT_detected

lib2d<-lib2[lib2$RT_detected>=80,]
lib2d<-lib2d[lib2d$RT_detected<95,]

RT1<-lib1[,c(1,3,8)]
RT2<-lib2d[,c(1,3,8)]
RT1<-RT1[RT1$modification_sequence%in%RT2$modification_sequence,]
RT2<-RT2[RT2$modification_sequence%in%RT1$modification_sequence,]
RT1<-RT1[!duplicated(RT1$modification_sequence),2:3]
RT2<-RT2[!duplicated(RT2$modification_sequence),2:3]
colnames(RT2)[1]<-"RT_detected_2"
RT<-join(RT1,RT2)
fit <- coef(lm(RT$RT_detected ~ RT$RT_detected_2))
lib2d$RT_detected<-(lib2d$RT_detected*fit[2])+fit[1]
lib2d$iRT<-lib2d$RT_detected

lib2e<-lib2[lib2$RT_detected>=95,]

RT1<-lib1[,c(1,3,8)]
RT2<-lib2e[,c(1,3,8)]
RT1<-RT1[RT1$modification_sequence%in%RT2$modification_sequence,]
RT2<-RT2[RT2$modification_sequence%in%RT1$modification_sequence,]
RT1<-RT1[!duplicated(RT1$modification_sequence),2:3]
RT2<-RT2[!duplicated(RT2$modification_sequence),2:3]
colnames(RT2)[1]<-"RT_detected_2"
RT<-join(RT1,RT2)
fit <- coef(lm(RT$RT_detected ~ RT$RT_detected_2))
lib2e$RT_detected<-(lib2e$RT_detected*fit[2])+fit[1]
lib2e$iRT<-lib2e$RT_detected

lib2<-rbind(lib2a,lib2b,lib2c,lib2d,lib2e)

dir.create("11corr")
setwd("11corr")

buildSpectraLibPair(lib1, lib2, outputFile = "ExtendedLib_2.txt", plot = TRUE,
                    clean = FALSE, merge = TRUE, parseAcc = TRUE, consolidateAccession = TRUE,
                    nomod=TRUE, conf.cutoff=0.9)

file.rename(from=paste(getwd(),"/ExtendedLib_2.txt",sep=""), 
            to=paste(workDir,"/ExtendedLib_2.txt", sep=""))
setwd(workDir)

#12

conf2<-read.xlsx("160909_04_NP_150_DDA__FDR.xlsx", sheet = "Search Summary", colNames = FALSE, rows=33)[1,4]
lib1<-readLibFile("ExtendedLib_2.txt",clean=FALSE)
lib2<-readLibFile("160909_04_NP_150_DDA.txt", clean = TRUE, intensity.cutoff = 5, conf.cutoff = conf2,
                  nomod = TRUE, nomc = FALSE, enz = "trypsin")
lib2<-canonicalFormat(lib2, format = "peakview")
lib2<-lib2[lib2$uniprot_id!="[ RT-Cal protein ]",]
lib2<-lib2[lib2$uniprot_id!="1/[ RT-Cal protein ]",]
lib2<-lib2[!grepl("NA",rownames(lib2)),]
lib1$RT_detected<-lib1$iRT

dir.create("12")
setwd("12")
checkQuality(lib1,lib2)

#12corr

lib2a<-lib2[lib2$RT_detected<55,]

RT1<-lib1[,c(1,3,8)]
RT2<-lib2a[,c(1,3,8)]
RT1<-RT1[RT1$modification_sequence%in%RT2$modification_sequence,]
RT2<-RT2[RT2$modification_sequence%in%RT1$modification_sequence,]
RT1<-RT1[!duplicated(RT1$modification_sequence),2:3]
RT2<-RT2[!duplicated(RT2$modification_sequence),2:3]
colnames(RT2)[1]<-"RT_detected_2"
RT<-join(RT1,RT2)
fit <- coef(lm(RT$RT_detected ~ RT$RT_detected_2))
lib2a$RT_detected<-(lib2a$RT_detected*fit[2])+fit[1]
lib2a$iRT<-lib2a$RT_detected

lib2b<-lib2[lib2$RT_detected>=55,]
lib2b<-lib2b[lib2b$RT_detected<97,]

RT1<-lib1[,c(1,3,8)]
RT2<-lib2b[,c(1,3,8)]
RT1<-RT1[RT1$modification_sequence%in%RT2$modification_sequence,]
RT2<-RT2[RT2$modification_sequence%in%RT1$modification_sequence,]
RT1<-RT1[!duplicated(RT1$modification_sequence),2:3]
RT2<-RT2[!duplicated(RT2$modification_sequence),2:3]
colnames(RT2)[1]<-"RT_detected_2"
RT<-join(RT1,RT2)
fit <- coef(lm(RT$RT_detected ~ RT$RT_detected_2))
lib2b$RT_detected<-(lib2b$RT_detected*fit[2])+fit[1]
lib2b$iRT<-lib2b$RT_detected

lib2c<-lib2[lib2$RT_detected>=97,]

RT1<-lib1[,c(1,3,8)]
RT2<-lib2c[,c(1,3,8)]
RT1<-RT1[RT1$modification_sequence%in%RT2$modification_sequence,]
RT2<-RT2[RT2$modification_sequence%in%RT1$modification_sequence,]
RT1<-RT1[!duplicated(RT1$modification_sequence),2:3]
RT2<-RT2[!duplicated(RT2$modification_sequence),2:3]
colnames(RT2)[1]<-"RT_detected_2"
RT<-join(RT1,RT2)
fit <- coef(lm(RT$RT_detected ~ RT$RT_detected_2))
lib2c$RT_detected<-(lib2c$RT_detected*fit[2])+fit[1]
lib2c$iRT<-lib2c$RT_detected

lib2<-rbind(lib2a,lib2b,lib2c)

dir.create("12corr")
setwd("12corr")

buildSpectraLibPair(lib1, lib2, outputFile = "ExtendedLib_2.txt", plot = TRUE,
                    clean = FALSE, merge = TRUE, parseAcc = TRUE, consolidateAccession = TRUE,
                    nomod=TRUE, conf.cutoff=0.9)

file.rename(from=paste(getwd(),"/ExtendedLib_2.txt",sep=""), 
            to=paste(workDir,"/ExtendedLib_2.txt", sep=""))
setwd(workDir)

#13

conf2<-read.xlsx("160909_06_TNIL_220_DDA__FDR.xlsx", sheet = "Search Summary", colNames = FALSE,
                 rows=33)[1,4]
lib1<-readLibFile("ExtendedLib_2.txt",clean=FALSE)
lib2<-readLibFile("160909_06_TNIL_220_DDA.txt", clean = TRUE, intensity.cutoff = 5, conf.cutoff = conf2,
                  nomod = TRUE, nomc = FALSE, enz = "trypsin")
lib2<-canonicalFormat(lib2, format = "peakview")
lib2<-lib2[lib2$uniprot_id!="[ RT-Cal protein ]",]
lib2<-lib2[lib2$uniprot_id!="1/[ RT-Cal protein ]",]
lib2<-lib2[!grepl("NA",rownames(lib2)),]
lib1$RT_detected<-lib1$iRT

dir.create("13")
setwd("13")
checkQuality(lib1,lib2)

#13corr

RT1<-lib1[,c(1,3,8)]
RT2<-lib2[,c(1,3,8)]
RT1<-RT1[RT1$modification_sequence%in%RT2$modification_sequence,]
RT2<-RT2[RT2$modification_sequence%in%RT1$modification_sequence,]
RT1<-RT1[!duplicated(RT1$modification_sequence),2:3]
RT2<-RT2[!duplicated(RT2$modification_sequence),2:3]
colnames(RT2)[1]<-"RT_detected_2"
RT<-join(RT1,RT2)
fit <- coef(lm(RT$RT_detected ~ RT$RT_detected_2))
lib2$RT_detected<-(lib2$RT_detected*fit[2])+fit[1]
lib2$iRT<-lib2$RT_detected

dir.create("13corr")
setwd("13corr")

buildSpectraLibPair(lib1, lib2, outputFile = "ExtendedLib_2.txt", plot = TRUE,
                    clean = FALSE, merge = TRUE, parseAcc = TRUE, consolidateAccession = TRUE,
                    nomod=TRUE, conf.cutoff=0.9)

file.rename(from=paste(getwd(),"/ExtendedLib_2.txt",sep=""), 
            to=paste(workDir,"/ExtendedLib_2.txt", sep=""))
setwd(workDir)

#14

conf2<-read.xlsx("161125_21_GPmyoDDA_TNIL_220__FDR.xlsx", sheet = "Search Summary", colNames = FALSE,
                 rows=33)[1,4]
lib1<-readLibFile("ExtendedLib_2.txt",clean=FALSE)
lib2<-readLibFile("161125_21_GPmyoDDA_TNIL_220.txt", clean = TRUE, intensity.cutoff = 5, 
                  conf.cutoff = conf2,nomod = TRUE, nomc = FALSE, enz = "trypsin")
lib2<-canonicalFormat(lib2, format = "peakview")
lib2<-lib2[lib2$uniprot_id!="[ RT-Cal protein ]",]
lib2<-lib2[lib2$uniprot_id!="1/[ RT-Cal protein ]",]
lib2<-lib2[!grepl("NA",rownames(lib2)),]
lib1$RT_detected<-lib1$iRT

dir.create("14")
setwd("14")
checkQuality(lib1,lib2)

#14corr

lib2a<-lib2[lib2$RT_detected<55,]


RT1<-lib1[,c(1,3,8)]
RT2<-lib2a[,c(1,3,8)]
RT1<-RT1[RT1$modification_sequence%in%RT2$modification_sequence,]
RT2<-RT2[RT2$modification_sequence%in%RT1$modification_sequence,]
RT1<-RT1[!duplicated(RT1$modification_sequence),2:3]
RT2<-RT2[!duplicated(RT2$modification_sequence),2:3]
colnames(RT2)[1]<-"RT_detected_2"
RT<-join(RT1,RT2)
fit <- coef(lm(RT$RT_detected ~ RT$RT_detected_2))
lib2a$RT_detected<-(lib2a$RT_detected*fit[2])+fit[1]
lib2a$iRT<-lib2a$RT_detected

lib2b<-lib2[lib2$RT_detected>=55,]
lib2b<-lib2b[lib2b$RT_detected<81,]

RT1<-lib1[,c(1,3,8)]
RT2<-lib2b[,c(1,3,8)]
RT1<-RT1[RT1$modification_sequence%in%RT2$modification_sequence,]
RT2<-RT2[RT2$modification_sequence%in%RT1$modification_sequence,]
RT1<-RT1[!duplicated(RT1$modification_sequence),2:3]
RT2<-RT2[!duplicated(RT2$modification_sequence),2:3]
colnames(RT2)[1]<-"RT_detected_2"
RT<-join(RT1,RT2)
fit <- coef(lm(RT$RT_detected ~ RT$RT_detected_2))
lib2b$RT_detected<-(lib2b$RT_detected*fit[2])+fit[1]
lib2b$iRT<-lib2b$RT_detected

lib2c<-lib2[lib2$RT_detected>=81,]

RT1<-lib1[,c(1,3,8)]
RT2<-lib2c[,c(1,3,8)]
RT1<-RT1[RT1$modification_sequence%in%RT2$modification_sequence,]
RT2<-RT2[RT2$modification_sequence%in%RT1$modification_sequence,]
RT1<-RT1[!duplicated(RT1$modification_sequence),2:3]
RT2<-RT2[!duplicated(RT2$modification_sequence),2:3]
colnames(RT2)[1]<-"RT_detected_2"
RT<-join(RT1,RT2)
fit <- coef(lm(RT$RT_detected ~ RT$RT_detected_2))
lib2c$RT_detected<-(lib2c$RT_detected*fit[2])+fit[1]
lib2c$iRT<-lib2c$RT_detected

lib2<-rbind(lib2a,lib2b,lib2c)

dir.create("14corr")
setwd("14corr")

buildSpectraLibPair(lib1, lib2, outputFile = "ExtendedLib_2.txt", plot = TRUE,
                    clean = FALSE, merge = TRUE, parseAcc = TRUE, consolidateAccession = TRUE,
                    nomod=TRUE, conf.cutoff=0.9)

file.rename(from=paste(getwd(),"/ExtendedLib_2.txt",sep=""), 
            to=paste(workDir,"/ExtendedLib_2.txt", sep=""))
setwd(workDir)

#15

conf2<-read.xlsx("OR7_20140724__FDR.xlsx", sheet = "Search Summary", colNames = FALSE,
                 rows=33)[1,4]
lib1<-readLibFile("ExtendedLib_2.txt",clean=FALSE)
lib2<-readLibFile("OR7_20140724.txt", clean = TRUE, intensity.cutoff = 5, conf.cutoff = conf2,
                  nomod = TRUE, nomc = FALSE, enz = "trypsin")
lib2<-canonicalFormat(lib2, format = "peakview")
lib2<-lib2[lib2$uniprot_id!="[ RT-Cal protein ]",]
lib2<-lib2[lib2$uniprot_id!="1/[ RT-Cal protein ]",]
lib2<-lib2[!grepl("NA",rownames(lib2)),]
lib1$RT_detected<-lib1$iRT

dir.create("15")
setwd("15")
checkQuality(lib1,lib2)

#15corr

lib2a<-lib2[lib2$RT_detected<26,]


RT1<-lib1[,c(1,3,8)]
RT2<-lib2a[,c(1,3,8)]
RT1<-RT1[RT1$modification_sequence%in%RT2$modification_sequence,]
RT2<-RT2[RT2$modification_sequence%in%RT1$modification_sequence,]
RT1<-RT1[!duplicated(RT1$modification_sequence),2:3]
RT2<-RT2[!duplicated(RT2$modification_sequence),2:3]
colnames(RT2)[1]<-"RT_detected_2"
RT<-join(RT1,RT2)
fit <- coef(lm(RT$RT_detected ~ RT$RT_detected_2))
lib2a$RT_detected<-(lib2a$RT_detected*fit[2])+fit[1]
lib2a$iRT<-lib2a$RT_detected

lib2b<-lib2[lib2$RT_detected>=26,]
lib2b<-lib2b[lib2b$RT_detected<50,]

RT1<-lib1[,c(1,3,8)]
RT2<-lib2b[,c(1,3,8)]
RT1<-RT1[RT1$modification_sequence%in%RT2$modification_sequence,]
RT2<-RT2[RT2$modification_sequence%in%RT1$modification_sequence,]
RT1<-RT1[!duplicated(RT1$modification_sequence),2:3]
RT2<-RT2[!duplicated(RT2$modification_sequence),2:3]
colnames(RT2)[1]<-"RT_detected_2"
RT<-join(RT1,RT2)
fit <- coef(lm(RT$RT_detected ~ RT$RT_detected_2))
lib2b$RT_detected<-(lib2b$RT_detected*fit[2])+fit[1]
lib2b$iRT<-lib2b$RT_detected

lib2c<-lib2[lib2$RT_detected>=50,]
lib2c<-lib2c[lib2c$RT_detected<87,]

RT1<-lib1[,c(1,3,8)]
RT2<-lib2c[,c(1,3,8)]
RT1<-RT1[RT1$modification_sequence%in%RT2$modification_sequence,]
RT2<-RT2[RT2$modification_sequence%in%RT1$modification_sequence,]
RT1<-RT1[!duplicated(RT1$modification_sequence),2:3]
RT2<-RT2[!duplicated(RT2$modification_sequence),2:3]
colnames(RT2)[1]<-"RT_detected_2"
RT<-join(RT1,RT2)
fit <- coef(lm(RT$RT_detected ~ RT$RT_detected_2))
lib2c$RT_detected<-(lib2c$RT_detected*fit[2])+fit[1]
lib2c$iRT<-lib2c$RT_detected

lib2d<-lib2[lib2$RT_detected>=87,]

RT1<-lib1[,c(1,3,8)]
RT2<-lib2d[,c(1,3,8)]
RT1<-RT1[RT1$modification_sequence%in%RT2$modification_sequence,]
RT2<-RT2[RT2$modification_sequence%in%RT1$modification_sequence,]
RT1<-RT1[!duplicated(RT1$modification_sequence),2:3]
RT2<-RT2[!duplicated(RT2$modification_sequence),2:3]
colnames(RT2)[1]<-"RT_detected_2"
RT<-join(RT1,RT2)
fit <- coef(lm(RT$RT_detected ~ RT$RT_detected_2))
lib2d$RT_detected<-(lib2d$RT_detected*fit[2])+fit[1]
lib2d$iRT<-lib2d$RT_detected

lib2<-rbind(lib2a,lib2b,lib2c,lib2d)

dir.create("15corr")
setwd("15corr")

buildSpectraLibPair(lib1, lib2, outputFile = "ExtendedLib_2.txt", plot = TRUE,
                    clean = FALSE, merge = TRUE, parseAcc = TRUE, consolidateAccession = TRUE,
                    nomod=TRUE, conf.cutoff=0.9)

file.rename(from=paste(getwd(),"/ExtendedLib_2.txt",sep=""), 
            to=paste(workDir,"/ExtendedLib_2.txt", sep=""))
setwd(workDir)

#16

conf2<-read.xlsx("OR9_201309019__FDR.xlsx", sheet = "Search Summary", colNames = FALSE,
                 rows=33)[1,4]
lib1<-readLibFile("ExtendedLib_2.txt",clean=FALSE)
lib2<-readLibFile("OR9_201309019.txt",clean = TRUE, intensity.cutoff = 5, conf.cutoff = conf2,
                  nomod = TRUE, nomc = FALSE, enz = "trypsin")
lib2<-canonicalFormat(lib2, format = "peakview")
lib2<-lib2[lib2$uniprot_id!="[ RT-Cal protein ]",]
lib2<-lib2[lib2$uniprot_id!="1/[ RT-Cal protein ]",]
lib2<-lib2[!grepl("NA",rownames(lib2)),]
lib1$RT_detected<-lib1$iRT

dir.create("16")
setwd("16")
checkQuality(lib1,lib2)

#16corr

lib2a<-lib2[lib2$RT_detected<70,]


RT1<-lib1[,c(1,3,8)]
RT2<-lib2a[,c(1,3,8)]
RT1<-RT1[RT1$modification_sequence%in%RT2$modification_sequence,]
RT2<-RT2[RT2$modification_sequence%in%RT1$modification_sequence,]
RT1<-RT1[!duplicated(RT1$modification_sequence),2:3]
RT2<-RT2[!duplicated(RT2$modification_sequence),2:3]
colnames(RT2)[1]<-"RT_detected_2"
RT<-join(RT1,RT2)
fit <- coef(lm(RT$RT_detected ~ RT$RT_detected_2))
lib2a$RT_detected<-(lib2a$RT_detected*fit[2])+fit[1]
lib2a$iRT<-lib2a$RT_detected

lib2b<-lib2[lib2$RT_detected>=70,]
lib2b<-lib2b[lib2b$RT_detected<90,]

RT1<-lib1[,c(1,3,8)]
RT2<-lib2b[,c(1,3,8)]
RT1<-RT1[RT1$modification_sequence%in%RT2$modification_sequence,]
RT2<-RT2[RT2$modification_sequence%in%RT1$modification_sequence,]
RT1<-RT1[!duplicated(RT1$modification_sequence),2:3]
RT2<-RT2[!duplicated(RT2$modification_sequence),2:3]
colnames(RT2)[1]<-"RT_detected_2"
RT<-join(RT1,RT2)
fit <- coef(lm(RT$RT_detected ~ RT$RT_detected_2))
lib2b$RT_detected<-(lib2b$RT_detected*fit[2])+fit[1]
lib2b$iRT<-lib2b$RT_detected

lib2c<-lib2[lib2$RT_detected>=90,]

RT1<-lib1[,c(1,3,8)]
RT2<-lib2c[,c(1,3,8)]
RT1<-RT1[RT1$modification_sequence%in%RT2$modification_sequence,]
RT2<-RT2[RT2$modification_sequence%in%RT1$modification_sequence,]
RT1<-RT1[!duplicated(RT1$modification_sequence),2:3]
RT2<-RT2[!duplicated(RT2$modification_sequence),2:3]
colnames(RT2)[1]<-"RT_detected_2"
RT<-join(RT1,RT2)
fit <- coef(lm(RT$RT_detected ~ RT$RT_detected_2))
lib2c$RT_detected<-(lib2c$RT_detected*fit[2])+fit[1]
lib2c$iRT<-lib2c$RT_detected


lib2<-rbind(lib2a,lib2b,lib2c)

dir.create("16corr")
setwd("16corr")

buildSpectraLibPair(lib1, lib2, outputFile = "ExtendedLib_2.txt", plot = TRUE,
                    clean = FALSE, merge = TRUE, parseAcc = TRUE, consolidateAccession = TRUE,
                    nomod=TRUE, conf.cutoff=0.9)

file.rename(from=paste(getwd(),"/ExtendedLib_2.txt",sep=""), 
            to=paste(workDir,"/ExtendedLib_2.txt", sep=""))
setwd(workDir)


#Here we start merging the 3 intermediate sublibraries, no cleaning required at this stage

#17

lib1<-readLibFile("ExtendedLib_Myometrium.txt",clean=FALSE)
lib2<-readLibFile("ExtendedLib_2.txt",clean=FALSE)

lib1$RT_detected<-lib1$iRT
lib2$RT_detected<-lib2$iRT

dir.create("17")
setwd("17")
checkQuality(lib1,lib2)

#17corr

lib2a<-lib2[lib2$RT_detected<50,]

RT1<-lib1[,c(1,3,8)]
RT2<-lib2a[,c(1,3,8)]
RT1<-RT1[RT1$modification_sequence%in%RT2$modification_sequence,]
RT2<-RT2[RT2$modification_sequence%in%RT1$modification_sequence,]
RT1<-RT1[!duplicated(RT1$modification_sequence),2:3]
RT2<-RT2[!duplicated(RT2$modification_sequence),2:3]
colnames(RT2)[1]<-"RT_detected_2"
RT<-join(RT1,RT2)
fit <- coef(lm(RT$RT_detected ~ RT$RT_detected_2))
lib2a$RT_detected<-(lib2a$RT_detected*fit[2])+fit[1]
lib2a$iRT<-lib2a$RT_detected


lib2b<-lib2[lib2$RT_detected>=50,]
lib2b<-lib2b[lib2b$RT_detected<98,]

RT1<-lib1[,c(1,3,8)]
RT2<-lib2b[,c(1,3,8)]
RT1<-RT1[RT1$modification_sequence%in%RT2$modification_sequence,]
RT2<-RT2[RT2$modification_sequence%in%RT1$modification_sequence,]
RT1<-RT1[!duplicated(RT1$modification_sequence),2:3]
RT2<-RT2[!duplicated(RT2$modification_sequence),2:3]
colnames(RT2)[1]<-"RT_detected_2"
RT<-join(RT1,RT2)
fit <- coef(lm(RT$RT_detected ~ RT$RT_detected_2))
lib2b$RT_detected<-(lib2b$RT_detected*fit[2])+fit[1]
lib2b$iRT<-lib2b$RT_detected

lib2c<-lib2[lib2$RT_detected>=98,]
lib2c<-lib2c[lib2c$RT_detected<105,]


RT1<-lib1[,c(1,3,8)]
RT2<-lib2c[,c(1,3,8)]
RT1<-RT1[RT1$modification_sequence%in%RT2$modification_sequence,]
RT2<-RT2[RT2$modification_sequence%in%RT1$modification_sequence,]
RT1<-RT1[!duplicated(RT1$modification_sequence),2:3]
RT2<-RT2[!duplicated(RT2$modification_sequence),2:3]
colnames(RT2)[1]<-"RT_detected_2"
RT<-join(RT1,RT2)
fit <- coef(lm(RT$RT_detected ~ RT$RT_detected_2))
lib2c$RT_detected<-(lib2c$RT_detected*fit[2])+fit[1]
lib2c$iRT<-lib2c$RT_detected

lib2d<-lib2[lib2$RT_detected>=105,]


RT1<-lib1[,c(1,3,8)]
RT2<-lib2c[,c(1,3,8)]
RT1<-RT1[RT1$modification_sequence%in%RT2$modification_sequence,]
RT2<-RT2[RT2$modification_sequence%in%RT1$modification_sequence,]
RT1<-RT1[!duplicated(RT1$modification_sequence),2:3]
RT2<-RT2[!duplicated(RT2$modification_sequence),2:3]
colnames(RT2)[1]<-"RT_detected_2"
RT<-join(RT1,RT2)
fit <- coef(lm(RT$RT_detected ~ RT$RT_detected_2))
lib2c$RT_detected<-(lib2c$RT_detected*fit[2])+fit[1]
lib2c$iRT<-lib2c$RT_detected


lib2<-rbind(lib2a,lib2b,lib2c,lib2d)

dir.create("17corr")
setwd("17corr")

buildSpectraLibPair(lib1, lib2, outputFile = "ExtendedLib_Myometrium.txt", plot = TRUE,
                    clean = FALSE, merge = TRUE, parseAcc = TRUE, consolidateAccession = TRUE,
                    nomod=TRUE, conf.cutoff=0.9)

file.rename(from=paste(getwd(),"/ExtendedLib_Myometrium.txt",sep=""), 
            to=paste(workDir,"/ExtendedLib_Myometrium.txt", sep=""))
setwd(workDir)


#18

lib1<-readLibFile("ExtendedLib_Myometrium.txt",clean=FALSE)
lib2<-readLibFile("ExtendedLib_1.txt",clean=FALSE)

lib1$RT_detected<-lib1$iRT
lib2$RT_detected<-lib2$iRT

dir.create("18")
setwd("18")
checkQuality(lib1,lib2)

#18corr

lib2a<-lib2[lib2$RT_detected<55,]
lib2a<-lib2a[lib2a$RT_detected>=26,]

RT1<-lib1[,c(1,3,8)]
RT2<-lib2a[,c(1,3,8)]
RT1<-RT1[RT1$modification_sequence%in%RT2$modification_sequence,]
RT2<-RT2[RT2$modification_sequence%in%RT1$modification_sequence,]
RT1<-RT1[!duplicated(RT1$modification_sequence),2:3]
RT2<-RT2[!duplicated(RT2$modification_sequence),2:3]
colnames(RT2)[1]<-"RT_detected_2"
RT<-join(RT1,RT2)
fit <- coef(lm(RT$RT_detected ~ RT$RT_detected_2))
lib2a$RT_detected<-(lib2a$RT_detected*fit[2])+fit[1]
lib2a$iRT<-lib2a$RT_detected


lib2b<-lib2[lib2$RT_detected>=54,]
lib2b<-lib2b[lib2b$RT_detected<100,]

RT1<-lib1[,c(1,3,8)]
RT2<-lib2b[,c(1,3,8)]
RT1<-RT1[RT1$modification_sequence%in%RT2$modification_sequence,]
RT2<-RT2[RT2$modification_sequence%in%RT1$modification_sequence,]
RT1<-RT1[!duplicated(RT1$modification_sequence),2:3]
RT2<-RT2[!duplicated(RT2$modification_sequence),2:3]
colnames(RT2)[1]<-"RT_detected_2"
RT<-join(RT1,RT2)
fit <- coef(lm(RT$RT_detected ~ RT$RT_detected_2))
lib2b$RT_detected<-(lib2b$RT_detected*fit[2])+fit[1]
lib2b$iRT<-lib2b$RT_detected

lib2c<-lib2[lib2$RT_detected>=99,]
lib2c<-lib2c[lib2c$RT_detected<120,]


RT1<-lib1[,c(1,3,8)]
RT2<-lib2c[,c(1,3,8)]
RT1<-RT1[RT1$modification_sequence%in%RT2$modification_sequence,]
RT2<-RT2[RT2$modification_sequence%in%RT1$modification_sequence,]
RT1<-RT1[!duplicated(RT1$modification_sequence),2:3]
RT2<-RT2[!duplicated(RT2$modification_sequence),2:3]
colnames(RT2)[1]<-"RT_detected_2"
RT<-join(RT1,RT2)
fit <- coef(lm(RT$RT_detected ~ RT$RT_detected_2))
lib2c$RT_detected<-(lib2c$RT_detected*fit[2])+fit[1]
lib2c$iRT<-lib2c$RT_detected

lib2d<-lib2[lib2$RT_detected>=119,]


RT1<-lib1[,c(1,3,8)]
RT2<-lib2d[,c(1,3,8)]
RT1<-RT1[RT1$modification_sequence%in%RT2$modification_sequence,]
RT2<-RT2[RT2$modification_sequence%in%RT1$modification_sequence,]
RT1<-RT1[!duplicated(RT1$modification_sequence),2:3]
RT2<-RT2[!duplicated(RT2$modification_sequence),2:3]
colnames(RT2)[1]<-"RT_detected_2"
RT<-join(RT1,RT2)
fit <- coef(lm(RT$RT_detected ~ RT$RT_detected_2))
lib2d$RT_detected<-(lib2d$RT_detected*fit[2])+fit[1]
lib2d$iRT<-lib2d$RT_detected


lib2<-rbind(lib2a,lib2b,lib2c,lib2d)

dir.create("18corr")
setwd("18corr")

buildSpectraLibPair(lib1, lib2, outputFile = "ExtendedLib_Myometrium.txt", plot = TRUE,
                    clean = FALSE, merge = TRUE, parseAcc = TRUE, consolidateAccession = TRUE,
                    nomod=TRUE, conf.cutoff=0.9)

file.rename(from=paste(getwd(),"/ExtendedLib_Myometrium.txt",sep=""), 
            to=paste(workDir,"/ExtendedLib_Myometrium.txt", sep=""))
setwd(workDir)


#The final library is now being loaded to remove peptide entries with less then 5 fragment ions
x<-read.delim("ExtendedLib_Myometrium.txt")
y<-as.data.frame(table(x$stripped_sequence))
y<-y[y$Freq>=5,]
x<-x[x$stripped_sequence%in%y$Var1,]
write.table(x,"ExtendedLib_Myometrium.txt",sep="\t")
```
