## Supplementary material for "The generation of a comprehensive spectral library for the analysis of the guinea pig proteome by SWATH-MS": R script 4

```
# Here, all the tissue specific libraries (created using dedicated scripts) are being combined to form a complete "total" guinea pig spectral library


library(SwathXtend)
library(plyr)
library(openxlsx)

workDir<-getwd()

#1

#Loading files, No further cleaning and cutoffs 

lib1<-readLibFile("ExtendedLib_Cerebrum.txt", clean = FALSE)
lib2<-readLibFile("ExtendedLib_Colon.txt", clean = FALSE)

#correcting the file format (removing the retension time calibrant and nonannotated peptides)
lib1<-canonicalFormat(lib1, format = "peakview")
lib2<-canonicalFormat(lib2, format = "peakview")
lib1<-lib1[lib1$uniprot_id!="[ RT-Cal protein ]",]
lib1<-lib1[lib1$uniprot_id!="1/[ RT-Cal protein ]",]
lib2<-lib2[lib2$uniprot_id!="[ RT-Cal protein ]",]
lib2<-lib2[lib2$uniprot_id!="1/[ RT-Cal protein ]",]
lib1<-lib1[!grepl("NA",rownames(lib1)),]
lib2<-lib2[!grepl("NA",rownames(lib2)),]

#creating a directory for the output files (quality measures for the first step of the library assembly)
dir.create("1")
setwd("1")

# Checking the linearity of RT correlation. If nonlinear (not the case here), the gradient is divided to linear fragments. 
# QC measures get saved in dedicated folder.

checkQuality(lib1,lib2)

#Libraries are merged. 

buildSpectraLibPair(lib1, lib2, outputFile = "Total_library.txt", plot = TRUE,
                    clean = FALSE, merge = TRUE, parseAcc = TRUE, consolidateAccession = TRUE,
                    nomod=TRUE, conf.cutoff=0.9)

# The merged library is moved to the working directory and will be used as the base library in the following steps.

file.rename(from=paste(getwd(),"/Total_library.txt",sep=""), to=paste(workDir,"/Total_library.txt", sep=""))
setwd(workDir)


#2

#The process described above is then repeated untill all the partial libraries are joined.

lib1<-readLibFile("Total_library.txt", clean = FALSE)
lib2<-readLibFile("ExtendedLib_Duoadenum.txt", clean = FALSE)
lib1<-canonicalFormat(lib1, format = "peakview")
lib2<-canonicalFormat(lib2, format = "peakview")
lib1<-lib1[lib1$uniprot_id!="[ RT-Cal protein ]",]
lib1<-lib1[lib1$uniprot_id!="1/[ RT-Cal protein ]",]
lib2<-lib2[lib2$uniprot_id!="[ RT-Cal protein ]",]
lib2<-lib2[lib2$uniprot_id!="1/[ RT-Cal protein ]",]
lib1<-lib1[!grepl("NA",rownames(lib1)),]
lib2<-lib2[!grepl("NA",rownames(lib2)),]


dir.create("2")
setwd("2")
checkQuality(lib1,lib2)

buildSpectraLibPair(lib1, lib2, outputFile = "Total_library.txt", plot = TRUE,
                    clean = FALSE, merge = TRUE, parseAcc = TRUE, consolidateAccession = TRUE,
                    nomod=TRUE, conf.cutoff=0.9)

file.rename(from=paste(getwd(),"/Total_library.txt",sep=""), to=paste(workDir,"/Total_library.txt", sep=""))
setwd(workDir)

#3

lib1<-readLibFile("Total_library.txt", clean = FALSE)
lib2<-readLibFile("ExtendedLib_Fat.txt", clean = FALSE)
lib1<-canonicalFormat(lib1, format = "peakview")
lib2<-canonicalFormat(lib2, format = "peakview")
lib1<-lib1[lib1$uniprot_id!="[ RT-Cal protein ]",]
lib1<-lib1[lib1$uniprot_id!="1/[ RT-Cal protein ]",]
lib2<-lib2[lib2$uniprot_id!="[ RT-Cal protein ]",]
lib2<-lib2[lib2$uniprot_id!="1/[ RT-Cal protein ]",]
lib1<-lib1[!grepl("NA",rownames(lib1)),]
lib2<-lib2[!grepl("NA",rownames(lib2)),]


dir.create("3")
setwd("3")
checkQuality(lib1,lib2)

buildSpectraLibPair(lib1, lib2, outputFile = "Total_library.txt", plot = TRUE,
                    clean = FALSE, merge = TRUE, parseAcc = TRUE, consolidateAccession = TRUE,
                    nomod=TRUE, conf.cutoff=0.9)

file.rename(from=paste(getwd(),"/Total_library.txt",sep=""), to=paste(workDir,"/Total_library.txt", sep=""))
setwd(workDir)

#4

lib1<-readLibFile("Total_library.txt", clean = FALSE)
lib2<-readLibFile("ExtendedLib_Kidney.txt", clean = FALSE)
lib1<-canonicalFormat(lib1, format = "peakview")
lib2<-canonicalFormat(lib2, format = "peakview")
lib1<-lib1[lib1$uniprot_id!="[ RT-Cal protein ]",]
lib1<-lib1[lib1$uniprot_id!="1/[ RT-Cal protein ]",]
lib2<-lib2[lib2$uniprot_id!="[ RT-Cal protein ]",]
lib2<-lib2[lib2$uniprot_id!="1/[ RT-Cal protein ]",]
lib1<-lib1[!grepl("NA",rownames(lib1)),]
lib2<-lib2[!grepl("NA",rownames(lib2)),]


dir.create("4")
setwd("4")
checkQuality(lib1,lib2)

buildSpectraLibPair(lib1, lib2, outputFile = "Total_library.txt", plot = TRUE,
                    clean = FALSE, merge = TRUE, parseAcc = TRUE, consolidateAccession = TRUE,
                    nomod=TRUE, conf.cutoff=0.9)

file.rename(from=paste(getwd(),"/Total_library.txt",sep=""), to=paste(workDir,"/Total_library.txt", sep=""))
setwd(workDir)

#5

lib1<-readLibFile("Total_library.txt", clean = FALSE)
lib2<-readLibFile("ExtendedLib_LargeIntestine.txt", clean = FALSE)
lib1<-canonicalFormat(lib1, format = "peakview")
lib2<-canonicalFormat(lib2, format = "peakview")
lib1<-lib1[lib1$uniprot_id!="[ RT-Cal protein ]",]
lib1<-lib1[lib1$uniprot_id!="1/[ RT-Cal protein ]",]
lib2<-lib2[lib2$uniprot_id!="[ RT-Cal protein ]",]
lib2<-lib2[lib2$uniprot_id!="1/[ RT-Cal protein ]",]
lib1<-lib1[!grepl("NA",rownames(lib1)),]
lib2<-lib2[!grepl("NA",rownames(lib2)),]


dir.create("5")
setwd("5")
checkQuality(lib1,lib2)

buildSpectraLibPair(lib1, lib2, outputFile = "Total_library.txt", plot = TRUE,
                    clean = FALSE, merge = TRUE, parseAcc = TRUE, consolidateAccession = TRUE,
                    nomod=TRUE, conf.cutoff=0.9)

file.rename(from=paste(getwd(),"/Total_library.txt",sep=""), to=paste(workDir,"/Total_library.txt", sep=""))
setwd(workDir)

#6

lib1<-readLibFile("Total_library.txt", clean = FALSE)
lib2<-readLibFile("ExtendedLib_Liver.txt", clean = FALSE)
lib1<-canonicalFormat(lib1, format = "peakview")
lib2<-canonicalFormat(lib2, format = "peakview")
lib1<-lib1[lib1$uniprot_id!="[ RT-Cal protein ]",]
lib1<-lib1[lib1$uniprot_id!="1/[ RT-Cal protein ]",]
lib2<-lib2[lib2$uniprot_id!="[ RT-Cal protein ]",]
lib2<-lib2[lib2$uniprot_id!="1/[ RT-Cal protein ]",]
lib1<-lib1[!grepl("NA",rownames(lib1)),]
lib2<-lib2[!grepl("NA",rownames(lib2)),]

dir.create("6")
setwd("6")
checkQuality(lib1,lib2)

#based on the visual inspection, the retension time correlation is non-linear
#The External library lib2 (the one that is being added to the "base library" lib1) is therefore manually divided into few, more linear fragments (lib2a-lib2b)


#6corr

#Linear fragment one
lib2a<-lib2[lib2$RT_detected<50,]

RT1<-lib1[,c(1,3,8)]
RT2<-lib2a[,c(1,3,8)]
RT1<-RT1[RT1$modification_sequence%in%RT2$modification_sequence,]
RT2<-RT2[RT2$modification_sequence%in%RT1$modification_sequence,]
RT1<-RT1[!duplicated(RT1$modification_sequence),2:3]
RT2<-RT2[!duplicated(RT2$modification_sequence),2:3]
colnames(RT2)[1]<-"RT_detected_2"
RT<-join(RT1,RT2)

# Linear regression is calculated and the coeficients used to correct the external library RT to mach the base library

fit <- coef(lm(RT$RT_detected ~ RT$RT_detected_2))
lib2a$RT_detected<-(lib2a$RT_detected*fit[2])+fit[1]
lib2a$iRT<-lib2a$RT_detected

#Linear fragment 2
lib2b<-lib2[lib2$RT_detected>=50,]

RT1<-lib1[,c(1,3,8)]
RT2<-lib2b[,c(1,3,8)]
RT1<-RT1[RT1$modification_sequence%in%RT2$modification_sequence,]
RT2<-RT2[RT2$modification_sequence%in%RT1$modification_sequence,]
RT1<-RT1[!duplicated(RT1$modification_sequence),2:3]
RT2<-RT2[!duplicated(RT2$modification_sequence),2:3]
colnames(RT2)[1]<-"RT_detected_2"
RT<-join(RT1,RT2)
fit <- coef(lm(RT$RT_detected ~ RT$RT_detected_2))
lib2b$RT_detected<-(lib2b$RT_detected*fit[2])+fit[1]
lib2b$iRT<-lib2b$RT_detected

#Linear fragments are then reassembled  

lib2<-rbind(lib2a,lib2b)

#Libraries are merged. QC measures get saved in dedicated subfolder. No cleaning is required at this step since both libraries were cleaned already.


buildSpectraLibPair(lib1, lib2, outputFile = "Total_library.txt", plot = TRUE,
                    clean = FALSE, merge = TRUE, parseAcc = TRUE, consolidateAccession = TRUE,
                    nomod=TRUE, conf.cutoff=0.9)

# The merged library is moved to the working directory and will be used as the base library in the following steps.

file.rename(from=paste(getwd(),"/Total_library.txt",sep=""), to=paste(workDir,"/Total_library.txt", sep=""))
setwd(workDir)

#7

lib1<-readLibFile("Total_library.txt", clean = FALSE)
lib2<-readLibFile("ExtendedLib_Lung.txt", clean = FALSE)
lib1<-canonicalFormat(lib1, format = "peakview")
lib2<-canonicalFormat(lib2, format = "peakview")
lib1<-lib1[lib1$uniprot_id!="[ RT-Cal protein ]",]
lib1<-lib1[lib1$uniprot_id!="1/[ RT-Cal protein ]",]
lib2<-lib2[lib2$uniprot_id!="[ RT-Cal protein ]",]
lib2<-lib2[lib2$uniprot_id!="1/[ RT-Cal protein ]",]
lib1<-lib1[!grepl("NA",rownames(lib1)),]
lib2<-lib2[!grepl("NA",rownames(lib2)),]


dir.create("7")
setwd("7")
# checkQuality(lib1,lib2)

buildSpectraLibPair(lib1, lib2, outputFile = "Total_library.txt", plot = TRUE,
                    clean = FALSE, merge = TRUE, parseAcc = TRUE, consolidateAccession = TRUE,
                    nomod=TRUE, conf.cutoff=0.9)

file.rename(from=paste(getwd(),"/Total_library.txt",sep=""), to=paste(workDir,"/Total_library.txt", sep=""))
setwd(workDir)

#8

lib1<-readLibFile("Total_library.txt", clean = FALSE)
lib2<-readLibFile("ExtendedLib_Ovaries.txt", clean = FALSE)
lib1<-canonicalFormat(lib1, format = "peakview")
lib2<-canonicalFormat(lib2, format = "peakview")
lib1<-lib1[lib1$uniprot_id!="[ RT-Cal protein ]",]
lib1<-lib1[lib1$uniprot_id!="1/[ RT-Cal protein ]",]
lib2<-lib2[lib2$uniprot_id!="[ RT-Cal protein ]",]
lib2<-lib2[lib2$uniprot_id!="1/[ RT-Cal protein ]",]
lib1<-lib1[!grepl("NA",rownames(lib1)),]
lib2<-lib2[!grepl("NA",rownames(lib2)),]


dir.create("8")
setwd("8")
# checkQuality(lib1,lib2)

buildSpectraLibPair(lib1, lib2, outputFile = "Total_library.txt", plot = TRUE,
                    clean = FALSE, merge = TRUE, parseAcc = TRUE, consolidateAccession = TRUE,
                    nomod=TRUE, conf.cutoff=0.9)

file.rename(from=paste(getwd(),"/Total_library.txt",sep=""), to=paste(workDir,"/Total_library.txt", sep=""))
setwd(workDir)

#9

lib1<-readLibFile("Total_library.txt", clean = FALSE)
lib2<-readLibFile("ExtendedLib_Pancreas.txt", clean = FALSE)
lib1<-canonicalFormat(lib1, format = "peakview")
lib2<-canonicalFormat(lib2, format = "peakview")
lib1<-lib1[lib1$uniprot_id!="[ RT-Cal protein ]",]
lib1<-lib1[lib1$uniprot_id!="1/[ RT-Cal protein ]",]
lib2<-lib2[lib2$uniprot_id!="[ RT-Cal protein ]",]
lib2<-lib2[lib2$uniprot_id!="1/[ RT-Cal protein ]",]
lib1<-lib1[!grepl("NA",rownames(lib1)),]
lib2<-lib2[!grepl("NA",rownames(lib2)),]


dir.create("9")
setwd("9")
# checkQuality(lib1,lib2)

buildSpectraLibPair(lib1, lib2, outputFile = "Total_library.txt", plot = TRUE,
                    clean = FALSE, merge = TRUE, parseAcc = TRUE, consolidateAccession = TRUE,
                    nomod=TRUE, conf.cutoff=0.9)

file.rename(from=paste(getwd(),"/Total_library.txt",sep=""), to=paste(workDir,"/Total_library.txt", sep=""))
setwd(workDir)

#10

lib1<-readLibFile("Total_library.txt", clean = FALSE)
lib2<-readLibFile("ExtendedLib_Placenta.txt", clean = FALSE)
lib1<-canonicalFormat(lib1, format = "peakview")
lib2<-canonicalFormat(lib2, format = "peakview")
lib1<-lib1[lib1$uniprot_id!="[ RT-Cal protein ]",]
lib1<-lib1[lib1$uniprot_id!="1/[ RT-Cal protein ]",]
lib2<-lib2[lib2$uniprot_id!="[ RT-Cal protein ]",]
lib2<-lib2[lib2$uniprot_id!="1/[ RT-Cal protein ]",]
lib1<-lib1[!grepl("NA",rownames(lib1)),]
lib2<-lib2[!grepl("NA",rownames(lib2)),]

#10corr

lib2a<-lib2[lib2$RT_detected<40,]

RT1<-lib1[,c(1,3,8)]
RT2<-lib2a[,c(1,3,8)]
RT1<-RT1[RT1$modification_sequence%in%RT2$modification_sequence,]
RT2<-RT2[RT2$modification_sequence%in%RT1$modification_sequence,]
RT1<-RT1[!duplicated(RT1$modification_sequence),2:3]
RT2<-RT2[!duplicated(RT2$modification_sequence),2:3]
colnames(RT2)[1]<-"RT_detected_2"
RT<-join(RT1,RT2)
fit <- coef(lm(RT$RT_detected ~ RT$RT_detected_2))
lib2a$RT_detected<-(lib2a$RT_detected*fit[2])+fit[1]
lib2a$iRT<-lib2a$RT_detected


lib2b<-lib2[lib2$RT_detected>=40,]

RT1<-lib1[,c(1,3,8)]
RT2<-lib2b[,c(1,3,8)]
RT1<-RT1[RT1$modification_sequence%in%RT2$modification_sequence,]
RT2<-RT2[RT2$modification_sequence%in%RT1$modification_sequence,]
RT1<-RT1[!duplicated(RT1$modification_sequence),2:3]
RT2<-RT2[!duplicated(RT2$modification_sequence),2:3]
colnames(RT2)[1]<-"RT_detected_2"
RT<-join(RT1,RT2)
fit <- coef(lm(RT$RT_detected ~ RT$RT_detected_2))
lib2b$RT_detected<-(lib2b$RT_detected*fit[2])+fit[1]
lib2b$iRT<-lib2b$RT_detected

lib2<-rbind(lib2a,lib2b)


dir.create("10")
setwd("10")
# checkQuality(lib1,lib2)

buildSpectraLibPair(lib1, lib2, outputFile = "Total_library.txt", plot = TRUE,
                    clean = FALSE, merge = TRUE, parseAcc = TRUE, consolidateAccession = TRUE,
                    nomod=TRUE, conf.cutoff=0.9)

file.rename(from=paste(getwd(),"/Total_library.txt",sep=""), to=paste(workDir,"/Total_library.txt", sep=""))
setwd(workDir)

#11

lib1<-readLibFile("Total_library.txt", clean = FALSE)
lib2<-readLibFile("ExtendedLib_SkeletalMusc.txt", clean = FALSE)
lib1<-canonicalFormat(lib1, format = "peakview")
lib2<-canonicalFormat(lib2, format = "peakview")
lib1<-lib1[lib1$uniprot_id!="[ RT-Cal protein ]",]
lib1<-lib1[lib1$uniprot_id!="1/[ RT-Cal protein ]",]
lib2<-lib2[lib2$uniprot_id!="[ RT-Cal protein ]",]
lib2<-lib2[lib2$uniprot_id!="1/[ RT-Cal protein ]",]
lib1<-lib1[!grepl("NA",rownames(lib1)),]
lib2<-lib2[!grepl("NA",rownames(lib2)),]


dir.create("11")
setwd("11")
# checkQuality(lib1,lib2)

#11corr

lib2a<-lib2[lib2$RT_detected<50,]

RT1<-lib1[,c(1,3,8)]
RT2<-lib2a[,c(1,3,8)]
RT1<-RT1[RT1$modification_sequence%in%RT2$modification_sequence,]
RT2<-RT2[RT2$modification_sequence%in%RT1$modification_sequence,]
RT1<-RT1[!duplicated(RT1$modification_sequence),2:3]
RT2<-RT2[!duplicated(RT2$modification_sequence),2:3]
colnames(RT2)[1]<-"RT_detected_2"
RT<-join(RT1,RT2)
fit <- coef(lm(RT$RT_detected ~ RT$RT_detected_2))
lib2a$RT_detected<-(lib2a$RT_detected*fit[2])+fit[1]
lib2a$iRT<-lib2a$RT_detected


lib2b<-lib2[lib2$RT_detected>=50,]

RT1<-lib1[,c(1,3,8)]
RT2<-lib2b[,c(1,3,8)]
RT1<-RT1[RT1$modification_sequence%in%RT2$modification_sequence,]
RT2<-RT2[RT2$modification_sequence%in%RT1$modification_sequence,]
RT1<-RT1[!duplicated(RT1$modification_sequence),2:3]
RT2<-RT2[!duplicated(RT2$modification_sequence),2:3]
colnames(RT2)[1]<-"RT_detected_2"
RT<-join(RT1,RT2)
fit <- coef(lm(RT$RT_detected ~ RT$RT_detected_2))
lib2b$RT_detected<-(lib2b$RT_detected*fit[2])+fit[1]
lib2b$iRT<-lib2b$RT_detected

lib2<-rbind(lib2a,lib2b)

buildSpectraLibPair(lib1, lib2, outputFile = "Total_library.txt", plot = TRUE,
                    clean = FALSE, merge = TRUE, parseAcc = TRUE, consolidateAccession = TRUE,
                    nomod=TRUE, conf.cutoff=0.9)

file.rename(from=paste(getwd(),"/Total_library.txt",sep=""), to=paste(workDir,"/Total_library.txt", sep=""))
setwd(workDir)

#12

lib1<-readLibFile("Total_library.txt", clean = FALSE)
lib2<-readLibFile("ExtendedLib_Small_Intestine.txt", clean = FALSE)
lib1<-canonicalFormat(lib1, format = "peakview")
lib2<-canonicalFormat(lib2, format = "peakview")
lib1<-lib1[lib1$uniprot_id!="[ RT-Cal protein ]",]
lib1<-lib1[lib1$uniprot_id!="1/[ RT-Cal protein ]",]
lib2<-lib2[lib2$uniprot_id!="[ RT-Cal protein ]",]
lib2<-lib2[lib2$uniprot_id!="1/[ RT-Cal protein ]",]
lib1<-lib1[!grepl("NA",rownames(lib1)),]
lib2<-lib2[!grepl("NA",rownames(lib2)),]


dir.create("12")
setwd("12")
# checkQuality(lib1,lib2)

buildSpectraLibPair(lib1, lib2, outputFile = "Total_library.txt", plot = TRUE,
                    clean = FALSE, merge = TRUE, parseAcc = TRUE, consolidateAccession = TRUE,
                    nomod=TRUE, conf.cutoff=0.9)

file.rename(from=paste(getwd(),"/Total_library.txt",sep=""), to=paste(workDir,"/Total_library.txt", sep=""))
setwd(workDir)

#13

lib1<-readLibFile("Total_library.txt", clean = FALSE)
lib2<-readLibFile("ExtendedLib_Stomach.txt", clean = FALSE)
lib1<-canonicalFormat(lib1, format = "peakview")
lib2<-canonicalFormat(lib2, format = "peakview")
lib1<-lib1[lib1$uniprot_id!="[ RT-Cal protein ]",]
lib1<-lib1[lib1$uniprot_id!="1/[ RT-Cal protein ]",]
lib2<-lib2[lib2$uniprot_id!="[ RT-Cal protein ]",]
lib2<-lib2[lib2$uniprot_id!="1/[ RT-Cal protein ]",]
lib1<-lib1[!grepl("NA",rownames(lib1)),]
lib2<-lib2[!grepl("NA",rownames(lib2)),]


dir.create("13")
setwd("13")
# checkQuality(lib1,lib2)

buildSpectraLibPair(lib1, lib2, outputFile = "Total_library.txt", plot = TRUE,
                    clean = FALSE, merge = TRUE, parseAcc = TRUE, consolidateAccession = TRUE,
                    nomod=TRUE, conf.cutoff=0.9)

file.rename(from=paste(getwd(),"/Total_library.txt",sep=""), to=paste(workDir,"/Total_library.txt", sep=""))
setwd(workDir)

#14

lib1<-readLibFile("Total_library.txt", clean = FALSE)
lib2<-readLibFile("ExtendedLib_heart.txt", clean = FALSE)
lib1<-canonicalFormat(lib1, format = "peakview")
lib2<-canonicalFormat(lib2, format = "peakview")
lib1<-lib1[lib1$uniprot_id!="[ RT-Cal protein ]",]
lib1<-lib1[lib1$uniprot_id!="1/[ RT-Cal protein ]",]
lib2<-lib2[lib2$uniprot_id!="[ RT-Cal protein ]",]
lib2<-lib2[lib2$uniprot_id!="1/[ RT-Cal protein ]",]
lib1<-lib1[!grepl("NA",rownames(lib1)),]
lib2<-lib2[!grepl("NA",rownames(lib2)),]


dir.create("14")
setwd("14")
# checkQuality(lib1,lib2)

#14corr

lib2a<-lib2[lib2$RT_detected<22,]

RT1<-lib1[,c(1,3,8)]
RT2<-lib2a[,c(1,3,8)]
RT1<-RT1[RT1$modification_sequence%in%RT2$modification_sequence,]
RT2<-RT2[RT2$modification_sequence%in%RT1$modification_sequence,]
RT1<-RT1[!duplicated(RT1$modification_sequence),2:3]
RT2<-RT2[!duplicated(RT2$modification_sequence),2:3]
colnames(RT2)[1]<-"RT_detected_2"
RT<-join(RT1,RT2)
fit <- coef(lm(RT$RT_detected ~ RT$RT_detected_2))
lib2a$RT_detected<-(lib2a$RT_detected*fit[2])+fit[1]
lib2a$iRT<-lib2a$RT_detected


lib2b<-lib2[lib2$RT_detected>=20,]
lib2b<-lib2b[lib2b$RT_detected<40,]

RT1<-lib1[,c(1,3,8)]
RT2<-lib2b[,c(1,3,8)]
RT1<-RT1[RT1$modification_sequence%in%RT2$modification_sequence,]
RT2<-RT2[RT2$modification_sequence%in%RT1$modification_sequence,]
RT1<-RT1[!duplicated(RT1$modification_sequence),2:3]
RT2<-RT2[!duplicated(RT2$modification_sequence),2:3]
colnames(RT2)[1]<-"RT_detected_2"
RT<-join(RT1,RT2)
fit <- coef(lm(RT$RT_detected ~ RT$RT_detected_2))
lib2b$RT_detected<-(lib2b$RT_detected*fit[2])+fit[1]
lib2b$iRT<-lib2b$RT_detected


lib2c<-lib2[lib2$RT_detected>=40,]
lib2c<-lib2c[lib2c$RT_detected<90,]

RT1<-lib1[,c(1,3,8)]
RT2<-lib2c[,c(1,3,8)]
RT1<-RT1[RT1$modification_sequence%in%RT2$modification_sequence,]
RT2<-RT2[RT2$modification_sequence%in%RT1$modification_sequence,]
RT1<-RT1[!duplicated(RT1$modification_sequence),2:3]
RT2<-RT2[!duplicated(RT2$modification_sequence),2:3]
colnames(RT2)[1]<-"RT_detected_2"
RT<-join(RT1,RT2)
fit <- coef(lm(RT$RT_detected ~ RT$RT_detected_2))
lib2c$RT_detected<-(lib2c$RT_detected*fit[2])+fit[1]
lib2c$iRT<-lib2c$RT_detected

lib2d<-lib2[lib2$RT_detected>=90,]

RT1<-lib1[,c(1,3,8)]
RT2<-lib2d[,c(1,3,8)]
RT1<-RT1[RT1$modification_sequence%in%RT2$modification_sequence,]
RT2<-RT2[RT2$modification_sequence%in%RT1$modification_sequence,]
RT1<-RT1[!duplicated(RT1$modification_sequence),2:3]
RT2<-RT2[!duplicated(RT2$modification_sequence),2:3]
colnames(RT2)[1]<-"RT_detected_2"
RT<-join(RT1,RT2)
fit <- coef(lm(RT$RT_detected ~ RT$RT_detected_2))
lib2d$RT_detected<-(lib2d$RT_detected*fit[2])+fit[1]
lib2d$iRT<-lib2d$RT_detected

lib2<-rbind(lib2a,lib2b,lib2c,lib2d)

buildSpectraLibPair(lib1, lib2, outputFile = "Total_library.txt", plot = TRUE,
                    clean = FALSE, merge = TRUE, parseAcc = TRUE, consolidateAccession = TRUE,
                    nomod=TRUE, conf.cutoff=0.9)

file.rename(from=paste(getwd(),"/Total_library.txt",sep=""), to=paste(workDir,"/Total_library.txt", sep=""))
setwd(workDir)

#15

lib1<-readLibFile("Total_library.txt", clean = FALSE)
lib2<-readLibFile("ExtendedLib_Myometrium.txt", clean = FALSE)
lib1<-canonicalFormat(lib1, format = "peakview")
lib2<-canonicalFormat(lib2, format = "peakview")
lib1<-lib1[lib1$uniprot_id!="[ RT-Cal protein ]",]
lib1<-lib1[lib1$uniprot_id!="1/[ RT-Cal protein ]",]
lib2<-lib2[lib2$uniprot_id!="[ RT-Cal protein ]",]
lib2<-lib2[lib2$uniprot_id!="1/[ RT-Cal protein ]",]
lib1<-lib1[!grepl("NA",rownames(lib1)),]
lib2<-lib2[!grepl("NA",rownames(lib2)),]


dir.create("15")
setwd("15")
# checkQuality(lib1,lib2a)


lib2a<-lib2[lib2$RT_detected<18,]
lib1a<-lib1[lib1$RT_detected>19,]
lib1a<-lib1a[lib1a$RT_detected<20.5,]

RT1<-lib1[,c(1,3,8)]
RT2<-lib2a[,c(1,3,8)]
RT1<-RT1[RT1$modification_sequence%in%RT2$modification_sequence,]
RT2<-RT2[RT2$modification_sequence%in%RT1$modification_sequence,]
RT1<-RT1[!duplicated(RT1$modification_sequence),2:3]
RT2<-RT2[!duplicated(RT2$modification_sequence),2:3]
colnames(RT2)[1]<-"RT_detected_2"
RT<-join(RT1,RT2)
fit <- coef(lm(RT$RT_detected ~ RT$RT_detected_2))
lib2a$RT_detected<-(lib2a$RT_detected*fit[2])+fit[1]
lib2a$iRT<-lib2a$RT_detected


lib2b<-lib2[lib2$RT_detected>=18,]


RT1<-lib1[,c(1,3,8)]
RT2<-lib2b[,c(1,3,8)]
RT1<-RT1[RT1$modification_sequence%in%RT2$modification_sequence,]
RT2<-RT2[RT2$modification_sequence%in%RT1$modification_sequence,]
RT1<-RT1[!duplicated(RT1$modification_sequence),2:3]
RT2<-RT2[!duplicated(RT2$modification_sequence),2:3]
colnames(RT2)[1]<-"RT_detected_2"
RT<-join(RT1,RT2)
fit <- coef(lm(RT$RT_detected ~ RT$RT_detected_2))
lib2b$RT_detected<-(lib2b$RT_detected*fit[2])+fit[1]
lib2b$iRT<-lib2b$RT_detected

lib2<-rbind(lib2a,lib2b)

buildSpectraLibPair(lib1, lib2, outputFile = "Total_library.txt", plot = TRUE,
                    clean = FALSE, merge = TRUE, parseAcc = TRUE, consolidateAccession = TRUE,
                    nomod=TRUE, conf.cutoff=0.9)

file.rename(from=paste(getwd(),"/Total_library.txt",sep=""), to=paste(workDir,"/Total_library.txt", sep=""))
setwd(workDir)

#The final library is now being loaded to remove peptide entries with less then 5 fragment ions
x<-read.delim("Total_library.txt")
y<-as.data.frame(table(x$stripped_sequence))
y<-y[y$Freq>=5,]
x<-x[x$stripped_sequence%in%y$Var1,]
write.table(x,"Total_library.txt",sep="\t")
```
